# Listening with generative models

**DOI:** 10.1101/2023.04.27.538626

**Authors:** Maddie Cusimano, Luke B. Hewitt, Josh H. McDermott

## Abstract

Perception has long been envisioned to use an internal model of the world to explain the causes of sensory signals. However, such accounts have historically not been testable, typically requiring intractable search through the space of possible explanations. Using auditory scenes as a case study, we leveraged contemporary computational tools to infer explanations of sounds in a candidate internal model of the auditory world (ecologically inspired audio synthesizers). Model inferences accounted for many classic illusions. Unlike traditional accounts of auditory illusions, the model is applicable to any sound, and exhibited human-like perceptual organization for real world sound mixtures. The combination of stimulus-computability and interpretable model structure enabled ‘rich falsification’, revealing additional assumptions about sound generation needed to account for perception. The results show how generative models can account for the perception of both classic illusions and everyday sensory signals, and provide the basis on which to build theories of perception.

## Introduction

Our perception of the world results from patterns of energy transduced by sensory receptors. But these sensory inputs alone do not specify the distal structure in the world that we perceive. For instance, many different three-dimensional objects are consistent with the same two-dimensional image, and many different combinations of sound sources are consistent with any observed sound waveform. Illusions show that out of many possible ways to interpret sensory inputs, human observers tend toward particular percepts, revealing constraints on perception that help resolve the ill-posed nature of perceptual problems. A key goal of perceptual science is to characterize these constraints and understand how they enable our real-world perceptual competencies.

Gestalt principles were an early approach to characterizing constraints on visual perception (1, 2), with analogues in auditory perception (3, 4). Nowadays these principles are taken to describe properties of a stimulus (“cues”) that determine its perceptual interpretation. For instance, in auditory perception, the principle of “common onset” states that sound components that begin at the same time tend to be grouped together. In modern extensions of the Gestalt approach (5), grouping cues are conceived as regularities that result from the causes of sensory data in the world. For example, the principle of common onset reflects the intuition that a single event would produce many frequency components simultaneously – so if such components are detected, they should be grouped.

These verbally-stated principles led to computational accounts of perception in which cue features are detected in sensory signals and used to determine stimulus grouping (3, 6–13). Models based on grouping cues have been used to explain targeted perceptual phenomena, such as contour grouping, figure/ground assignment, or grouping of tone sequences, but in practice do not come close to comprehensively accounting for the perceptual interpretation of real images or sounds. These limitations reflect the difficulty in specifying features that are predictive of perception. For instance, local features tend to be ambiguous, and can be interpreted differently depending on the surrounding context (14–19).

An alternative possibility is that perceptual systems might more directly utilize constraints on how sensory data is generated (20, 21). For instance, natural images are constrained by optics, which describes how light interacts with surfaces to create shading and shadows evident as luminance gradients in the resulting image. Similarly, natural sounds are constrained by acoustics, which describes how different types of physical or biological processes generate different types of sounds. Perceptual systems could capture these constraints using an internal ‘world model’ of how causes generate signals (Figure 1A).

**Figure 1.**
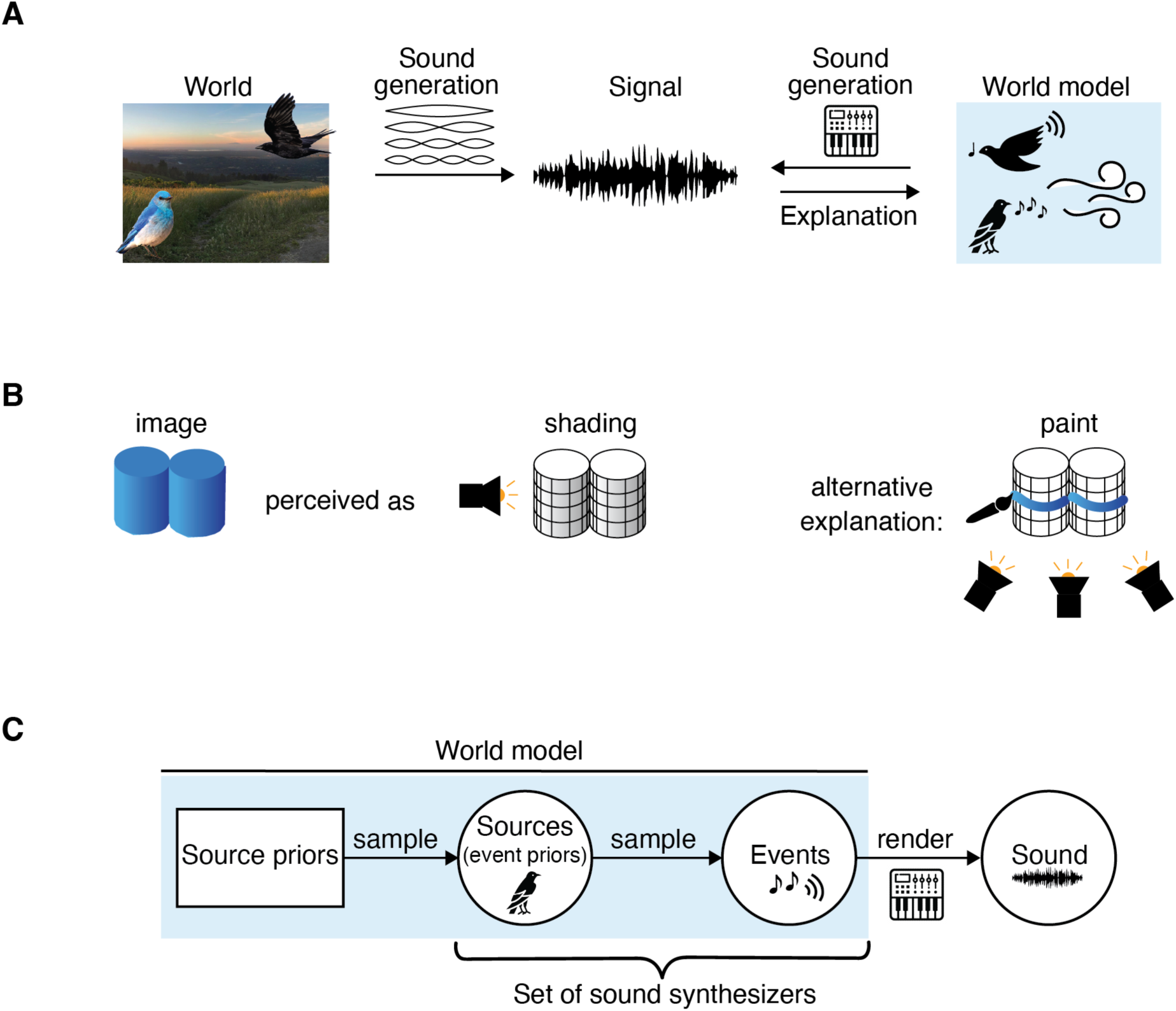
Overview of perception as inference in a world model. A) Generative processes in the world produce sound, which can be described with models of sound generation (acoustics). Perceptual systems could use an internal model of how causes generate signals (a “world model”) to explain the data, in terms of causes which generated the data. To generate signals from causes, this world model contains audio synthesizers. B) Example of inference in a visual world model (20). We perceive the image on the left to be a set of solid blue cylinders that is lit from the left. This percept can be considered an explanation of the image in terms of interacting causal variables like illumination and surface reflectance (shown in middle). But there are many alternative causal explanations for this image, for instance that the cylinders are lit uniformly and have gradients that are painted onto their surface (right). Perception could avoid this alternative explanation due to the low prior probability of surfaces being painted and illuminated in this way. C) High-level description of the proposed internal generative model of auditory scenes. Sources are sampled from a source prior distribution. Sources emit events that create sound. A source is thus an audio synthesizer. The model is expressed with a probabilistic program, which allows for scene descriptions that vary in dimensionality (the schematic here should not be mistaken for a graphical model, which would not allow for variable dimensionality). For image attributions, see acknowledgments.

In vision, this world model might resemble a model of optics that describes how illumination and three-dimensional objects (causes) interact to produce luminance patterns (signal). Perception could be a process of using such a world model to infer the causes that are likely to have generated the incoming signal (Figure 1B). In principle, probabilistic (Bayesian) inference provides the basis for inverting the world model to determine causes from signals (Figure 1A): given an input signal, the output of inference is a set of possible causes, each with an associated probability. While different causes can generate the same signal (i.e., perceptual inference is ill posed), some causes are more likely to have generated the signal than others, and a sufficiently accurate internal world model and inference procedure could enable us to arrive at an accurate perceptual interpretation most of the time (Figure 1B, alternative). The causes in the world model thus simultaneously specify what is inferred by perception and provide the constraints necessary to (probabilistically) solve the ill-posed computation.

Inference in generative models typically involves a search through the space of possible causes to find those that are plausible given the data. Despite the appeal of generative models for explaining perception (22), in practice, this search is computationally intractable for all but the simplest models, which has limited their scope as accounts of perception. With few exceptions (23, 24), prior applications of Bayesian inference to human perception have been limited to few-or fixed-dimensional domains (17, 25–33), operated on symbolic data rather than actual sensory signals (34–39), or faced intractable inference issues that prevented them from being fully evaluated (22, 40, 41).

Here we consider auditory scene analysis as a case study in perceptual organization, with which to revisit generative approaches to perception. Auditory scenes are an appealing starting point for a generative account of perception, because relatively simple generative models can synthesize a wide variety of naturalistic sounds. Nonetheless, one prevailing challenge is the difficulty of integrating structured models with raw audio signals as input. Most existing models are restricted to symbolic rather than acoustic input (37, 38, 42), and thus are only applicable to synthetic stimuli with simple symbolic descriptions (e.g. tone sequences). Because of this, we have lacked a comprehensive account of key phenomena in auditory scene analysis, despite many proposed conceptual approaches (43). It is also unclear whether putative principles for auditory perceptual organization which apply to synthetic sounds could also extend to explaining auditory scene analysis in everyday sounds (44).

To realize a theory of perception as inference in world model (20, 21), we leverage contemporary technical developments to render Bayesian inference in world models newly approachable. To build a rich, structured model of auditory scenes (Figure 1C), we use a generalization of graphical models called probabilistic programs (23, 45). To implement search, we take an “analysis-by-synthesis” approach, assessing bottom-up proposals about potential causes (“analysis”) via top-down “synthesis” using the generative model (4, 46). This strategy combines the benefits of fast pattern recognition with the explanatory power of Bayesian inference. To make analysis-by-synthesis tractable, we use deep learning to make bottom-up proposals (“amortized inference”) (47) and assess them top-down using stochastic variational inference to approximate the posterior, leveraging a differentiable generative model (48, 49). We compare our approach to deep neural networks that have had success in source separation tasks with naturalistic audio (i.e., reconstructing premixture waveforms from audio mixtures, which might be viewed as a form of auditory scene analysis).

Unlike previous symbolic models, our model can be evaluated on any sound signal, enabling it to be tested on both classic illusions and natural sounds. Experiments with human listeners show that the model inferences capture human perception for recorded environmental audio. The ability to evaluate the model on natural sounds also allows the model to be “richly falsified”, highlighting structure in natural sounds that our perceptual systems are attuned to, and that might otherwise be overlooked. Unlike neural networks for source separation, which reconstruct waveforms, our model outputs probabilities over symbolic scene descriptions that specify the number of sources, the properties of the inferred sources and the sound events that each source emits. We found that our model accounted for a variety of classic illusions in auditory perceptual organization, whereas source separation networks did not. These results show that rich Bayesian models of perception are now within reach, and can bridge traditional psychophysics with the perception of everyday sensory signals.

## Results

### Overview

To investigate the extent to which auditory scene analysis can be explained by a generative model of sound, we considered how to simply describe and render a variety of sounds. The resulting generative model provides a hypothesis for the internal model that might underlie human auditory scene analysis (Figure 1C), constituting a computational-level explanation in the sense of Marr (50). The model is accompanied by an analysis-by-synthesis inference procedure that we built to search the model space, enabling us to evaluate the model on audio signals. To test how well the generative model aligns with human perception, we first assessed whether inference in the model accounts for a set of classic auditory scene analysis illusions. These illusions are widely thought to elucidate principles of human hearing and thus provide a clear starting set of phenomena that any model should account for. We tested the role and importance of different aspects of the model by assessing the effect of various model “lesions„. To establish baseline performance on these illusions using alternative scene analysis strategies, we compared our model with an assortment of contemporary deep neural network source separation systems (as these are the main widely used alternative class of stimulus-computable model at present). Finally, we leveraged the ability to apply the model to any sound waveform, performing inference on mixtures of recorded everyday sounds. We ran experiments on human listeners to assess whether the model accounted for human perceptual organization of these real-world auditory scenes, using the model “failures” to reveal previously unappreciated constraints on human perception.

The model and inference procedure are briefly described in the following sections, with the intent of providing a high-level description that will enable the reader to understand the main results of the paper. For a full description of both the generative model and our inference procedure, see the Methods. The interested reader is also referred to Supplementary Algorithm 1 for pseudocode defining the model.

### Generative model

A generative model specifies 1) the structure of possible latent causes of sensory data, with associated prior probabilities describing how likely each cause is to occur in the world, and 2) a description of how these causes generate data, determining the likelihood of observed data under each possible cause. Together, these components provide probabilistic constraints that determine the Bayesian posterior distribution over causes given an observed sensory signal, p(cause|signal).

Our proposed model is inspired by observations of generative principles in everyday sounds, balanced by simplicity to enable tractable inference. Because everyday auditory scenes can include variable numbers and kinds of sources, we defined the model as a probabilistic program (23, 45) to allow for this variable structure. As for any generative model, the program consists of the prior and likelihood components mentioned above:

1. A sampling procedure which generates a hierarchical symbolic description of a scene, *S*, in terms of sources and the events they emit. The sampling procedure defines the prior distribution *p*(*S*) over this space of possible causes of sound, providing constraints on what sources and events are probable *a priori*.
2. A renderer which uses this symbolic scene representation *S* to generate an audio signal. The rendered sound can, in conjunction with a noise model, be used to evaluate the model likelihood *p*(*XS*) for any observed sound *X* (a noise model is needed to account for measurement error in sensory data and to allow for imperfect matches between model generated and observed data). The likelihood assesses how likely a scene description is to generate a particular sound.

Given an observed sound waveform *X*, the sampling procedure and renderer induce a posterior distribution over auditory scenes by Bayes’ rule: *p*(*S*|*X*) ∝ *p*(*S*) × *p*(*XS*). The most probable scene descriptions for an observed sound can then in principle be found via inference (searching through scene descriptions to find descriptions with high posterior probability). In practice, this search has traditionally been intractable due to the high dimensionality of the space of generative parameters. We begin by describing the qualitative structure of the sampling procedure and renderer, then the prior *p*(*S*) and likelihood *p*(*XS*), and then the inference procedure we used to make the search for probable scene descriptions more tractable.

### Generative model: Structure

Figure 2A shows examples of recorded everyday sounds. We took inspiration from such sounds to develop a flexible and widely applicable symbolic description of sound sources. The resulting model is considerably more expressive than previous generative models for auditory scene analysis, and can generate simple approximations of many everyday sounds. We nonetheless intentionally kept its structure as simple as possible so as to facilitate inference. The following three observations informed the model’s construction.

**Figure 2.**
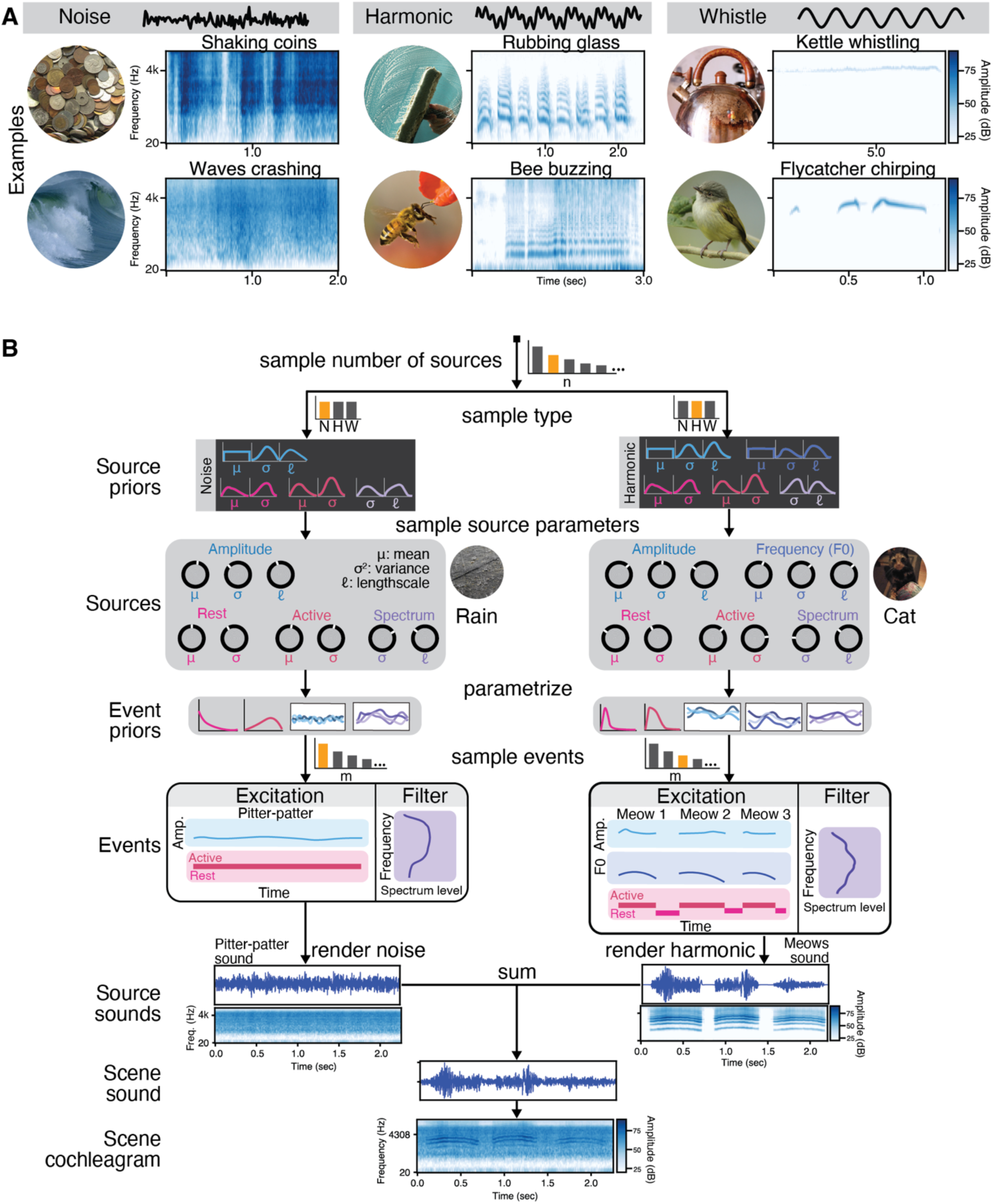
Generative model, illustrated with everyday recorded sound examples. A) The three sound types with natural sound examples. The examples demonstrate a variety of amplitude and frequency modulation, spectral shapes, and temporal patterns. Here and in other figures, sound examples are available on the project website, https://mcdermottlab.mit.edu/mcusi/bass/. B) A scene consists of any number of sources. Each source belongs to one sound type, which defines how the source is parametrized and rendered, and determines which source priors the source is sampled from. The sampled source parameters define distributions over events. A source can emit any number of events. Events consist of active and rest intervals, time-varying amplitude, and depending on the sound type, possibly a spectrum and/or time-varying fundamental frequency. Events are rendered into sound waveforms by combining an excitation and filter (cochleagrams of source sounds are shown only for visualization; they are not part of the generative model). The sounds generated by independent sources sum together to create a scene sound, which is transformed into a cochleagram for model inference. This scene cochleagram is the basis for the likelihood function. The number of sources, the type of sources, and the number of events can all change the dimensionality of the model, requiring it to be expressed as a probabilistic program. For image and sound attributions, see acknowledgments.

First, a substantial variety of everyday sounds can be described as coming from three broad classes (Figure 2A): noise, harmonic, and whistle sounds. Noise-like sounds are commonly produced by turbulence in air or fluids (e.g., waves crashing), or when large numbers of sounds superimpose to form textures (e.g., shaking coins) (51, 52); brief impact sounds are also often well described as short snippets of noise. Periodic sources produce harmonic sounds; these range from spoken vowels (53) to buzzes produced by bees’ rapid wingbeats (54) and squeaks produced by rubbing glass (55). Whistles are commonly created by air flow and resonance (e.g., in kettles) (56) and are produced in a variety of ways by animals across taxa (e.g., bird chirping) (57–59). Although some complex sound sources can involve more than one of these sound types, many natural sounds are dominated by a single sound type.

Second, many sounds can be described as being produced by multiple discrete, dynamic “events”, corresponding to when a source supplies energy to produce sound. For example, the squealing sound produced by rubbing glass starts and stops over time, corresponding to relatively discrete time intervals when force is being applied to the glass surface. Within those intervals, the sound continuously changes in fundamental frequency. The resulting events may be temporally extended over seconds (e.g., a kettle whistling) or more transient (e.g., the chirps of a bird), and can change in their properties from event to event (e.g., a dog panting in and out).

Third, the events emitted by a source reflect source-specific regularities in a variety of attributes, including event timing, fundamental frequency, amplitude, and spectral shape. For example, some sources tend to produce many regular events in quick succession (e.g., rubbing glass), while others produce long events that continuously vary (e.g., waves crashing). And while the chirps of a bird vary smoothly in fundamental frequency, the buzzing of a bee varies rapidly and erratically. Both stay in a relatively narrow range of frequency space.

These three observations are a starting point for a generative model of auditory scenes. In our model (Figure 2B), a scene description consists of sources which each emit a sequence of discrete events. A scene can contain any number of sources (Figure 2B, “sample number of sources„), each of which generates one type of sound: noise, harmonic, or whistle (Figure 2B, “sample type”). Sources of the same type can differ from each other in the attributes noted above, for example, in their spectral shape, in how loud they tend to be, or in how often they tend to emit events (characterized by the setting of the knobs in Figure 2B, which in the model are sampled from the source priors). These source parameters define the event priors, causing regularities across events produced by the same source (Figure 2B, “events”). Each source can emit any number of events. Each event is dynamic in time, e.g. changing in frequency or amplitude (Figure 2B, “Events„). Given a scene description S, the renderer synthesizes the sound emitted by each source (Figure 2B, “Source sounds„) and sums them to produce the scene sound waveform (Figure 2B, “Scene sound„).

The sound types in our model are parametrized as excitation-filter combinations commonly proposed for natural sound generation (53, 60, 61) (Figure 2B, excitation/filter split in the events panel). Sound is produced by an “excitation” that supplies sound energy and a “filter” that determines the spectral shape; in our model the excitation comes in one of three varieties determined by the sound type. Sounds produced by very different physical mechanisms (e.g. the beating wings of a bee versus the friction of rubbing glass) may still be best described by the same excitation type, and therefore can have the same parametrization in the model. A noise source produces aperiodic excitation with a pink (1/f) spectrum and an amplitude that varies over time. A harmonic source produces periodic excitation: a sum of harmonically related sinusoids with a time-varying fundamental frequency and amplitude. A whistle sound similarly produces a periodic excitation, but only generates the first harmonic. In whistle sounds, the filter can be considered fixed, and so we omit it for simplicity.

A source becomes “active” for a bounded time interval while emitting an event, and is otherwise silent while it “rests” in between events (Figure 2B, active/rest in the events panel). Events are defined by an onset when the source’s excitation begins (discretely), an offset when the excitation ends, a filter, and a time-varying excitation (Figure 2B, events panel). Each of these are sampled from event priors defined by the source parameters (represented by the knobs in Figure 2B).

### Generative model: Prior distributions

The qualitative properties of the generative model described in the preceding section are made precise with a sampling procedure for scene descriptions that defines the prior distribution *p*(*S*). To reflect the structure of the scene description (in which different sources emit events), *p*(*S*) is hierarchical (Figure 2B). There are prior distributions over source parameters for each sound type (source priors), and each sampled source defines prior distributions over events (event priors). Sources are assumed to be independent, whereas events emitted by a single source are sampled sequentially so that each event depends on the events preceding it.

This hierarchical prior can account for a variety of sound regularities. Figure 3A illustrates how the model can express sources with different temporal regularities. The short, consistent impacts of hammering nails is represented in the source parameters as a small value of µ (average duration) paired with a small value of σ^2^ (variance). By contrast, the song of a white-throated sparrow comprises mostly longer notes of more variable duration (large µ and large σ^2^). The source parameters µ and σ^2^ are sampled from the source prior; the durations of all events produced by the sampled source then follow an ‘event prior’ distribution governed by these source parameters (in particular, a log-normal distribution parametrized by µ and σ^2^). The same generative process (with different sampled source parameters) is used for the “rests” between events. Figure 3B shows the analogous hierarchical generative process for the time-varying fundamental frequency of the source excitation. Here the event priors are Gaussian processes, defined by source parameters µ (average F0), *σ* (range of F0 variation), and ℓ (pace of F0 variation). Because the Gaussian process is a prior on functions, it is depicted with samples in Figure 3B (fourth row). The amplitude of the excitation follows the same generative process but with different values defining the source priors. The Gaussian process priors for the excitation trajectories additionally account for situations where the excitation properties change from event to event (the non-stationarity illustrated in Figure 3B; see also Supplementary Figure 1). The role of the Gaussian process priors is discussed further in the *Model lesions* section below.

**Figure 3.**
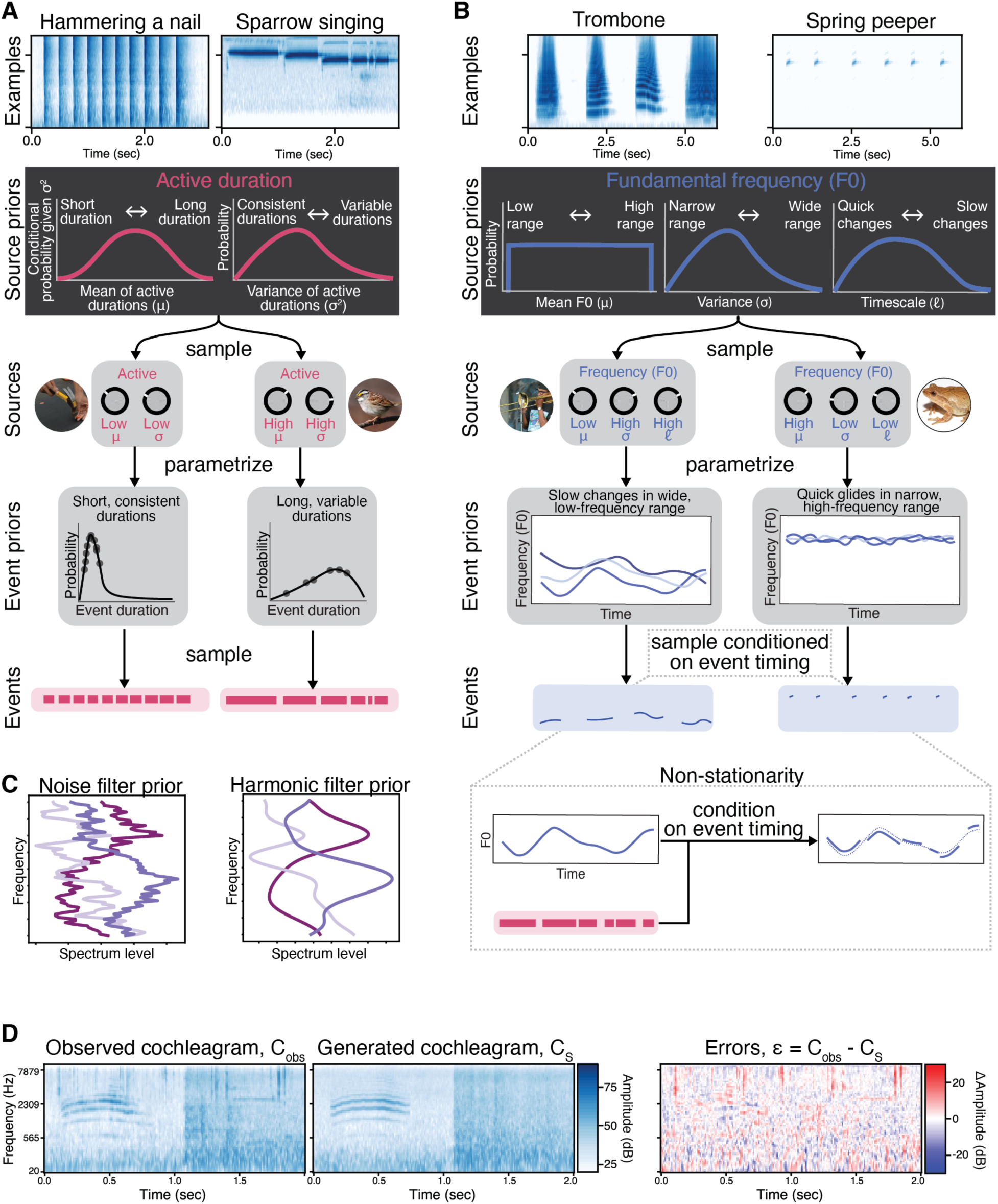
Prior and likelihood, illustrated with recorded natural sound examples. A) Hierarchical priors on event duration. Hammering nails results in short, consistent impacts, while the song of a white-throated sparrow comprises longer tones of variable duration. To capture such regularities, we use a hierarchical model. For each source, the source parameters μ (mean duration) and σ^2^ (variance) are sampled from the source priors (a normal-inverse-gamma distribution). The source parameters define the event priors (log-normal distributions) from which the event durations are sampled. Different source parameters capture different regularities: the hammer impacts can be modeled with a low mean and low variance, while the sparrow song requires a high mean and high variance. B) Hierarchical priors on fundamental frequency. The trombone’s fundamental frequency is in a low, wide frequency range, and changes slowly, while the spring peeper is in a high frequency range and changes quickly by a small amount. For each source, the source parameters μ (mean), σ (kernel variance) and ℓ (kernel lengthscale) are source parameters that are sampled from source priors. These source parameters define the Gaussian process event priors from which event fundamental frequency trajectories are sampled. Different source parameters capture different regularities: the trombone can be captured by low mean, high variance, and high lengthscale, while the spring peeper can be captured by high mean, low variance, and low lengthscale. We depict the Gaussian process priors by showing samples of potential trajectories. To account for the possibility that excitation trajectories might change in their properties from event to event (e.g., a dog panting creates different amplitudes on the in- and out-breath), we used a non-stationary Gaussian process kernel (see Supplementary Figure 1 for more detail). This required the Gaussian process event priors to be conditioned on the timing of events. C) Prior over filter shape. We used different priors for the filters of different sound types, so that the spectra of harmonic sources tended to be smoother than those of noise sources. This difference was implemented with a Ornstein-Uhlenbeck kernel for noises and a squared-exponential kernel for harmonics. We depict the Gaussian process priors by showing samples. D) To calculate the likelihood, the observed and generated cochleagrams are compared under a Gaussian noise model.

The spectrum of the source filter follows a generative process like that for the excitation, but is constant across all events. The Gaussian process priors for the spectrum of the source filter differ for noise and harmonic sources, embodying the tendency of harmonic sources to have smooth spectral envelopes (Figure 3C).

To determine the shape of the source priors, we fit each one to a dataset of sound textures (for noise) and speech, musical instruments, and birdsong (for periodic sounds; we assumed that the priors over fundamental frequency would be similar for harmonic and whistle sounds). The distributions we chose (a normal-inverse-gamma prior over the temporal variables, and Gaussian processes over the multivariate spectrum and excitation trajectory; see Methods) enable the use of differentiable sampling procedures and efficient inference platforms that have been developed for common distributions (62), thereby facilitating inference.

The overall model is depicted graphically in Figures 2 and 3, and is described in detail in the Methods. See Supplementary Algorithm 1 for pseudocode describing the full probabilistic program instantiating the generative model. Supplementary Figure 2 shows sounds sampled from *p*(*S*).

### Generative model: Likelihood

Given S, the symbolic description of the auditory scene, the renderer generates the sound waveform produced by each source. The sound waveforms corresponding to all sources are summed to produce the scene waveform, X_S_. The likelihood *p*(*X*_obs_|*S*) tells us how likely it is that a particular scene *S* will generate an observed sound, *X*_obs_. To compute *p*(*X*_obs_|*S*), the scene waveform *X*_S_ rendered from *S* must be compared to the observation *X*_obs_. To do so, the two waveforms are first converted to a cochleagram, a time frequency representation of sound that approximates the filtering properties of the human ear, and are then compared under a Gaussian noise model (Figure 3D). That is, the likelihood is the probability that the observed cochleagram *C*_obs_is a noisy measurement of the rendered cochleagram *C*_S_:

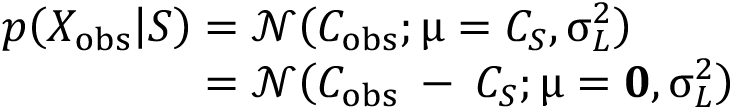

The motivation for basing the likelihood on the cochleagram rather than the sound waveform is that human discrimination is constrained by cochlear filtering (63). The cochleagram representation of sound has the additional benefit of tending to vary smoothly with respect to the continuous latent variables of our model, facilitating gradient-based inference. The assumed noise level σ_L_ is a free parameter (fixed to be the same value across all inferences).

### Inference

By Bayes’ rule, the prior *p*(*S*) and likelihood *p*(*XS*) induce a posterior distribution *p*(*S*|*X*) for an observed sound *X*, that is, *p*(*S*|*X*) ∝ *p*(*S*)*p*(*XS*). The posterior describes which hypothesized scenes are more likely explanations of the observed data. From this viewpoint, the computational goal of perception is to uncover the most likely explanation (or set of explanations) of the sound *X* given the world model.

Finding the hypotheses that are most likely under the posterior distribution involves two main challenges. First, posterior inference requires solving a difficult search problem. Because the generative model is expressive, the space of scene descriptions is vast, making it difficult to find good hypotheses to evaluate. Second, the posterior distribution is typically multimodal. Complex generative models (like ours) typically give rise to multiple plausible hypotheses (modes), such that even if we could find a region of scene space containing good hypotheses, multiple hypotheses must be compared since some may only be local optima. To successfully compare the modes, we must evaluate how much posterior probability mass corresponds to each mode (see Methods). However, because hypotheses are high-dimensional and contain complex dependencies between the various random variables, it is difficult to estimate the probability landscape surrounding a mode closely enough to ensure that all of the probability mass covered by a mode will be accounted for (64).

An inference algorithm must solve these challenges in order for us to evaluate our generative model on sound. We make no claim that the particular inference algorithm we developed provides an algorithmic-level account of perception (50). For our purposes, the most important function of our inference procedure is that it is one algorithmic approach that permits a solution to the computational challenges associated with generative models, allowing us to ask whether the generative model we proposed can explain human perception.

Nevertheless, to design our inference procedure, we took inspiration from ideas about how perceptual systems could solve these challenges, in particular analysis-by-synthesis (4, 46). Intuitively, the observed sound contains clues about likely components of a good hypothesis, pointing us to promising parts of the scene space. For instance, if we detect (using bottom-up pattern recognition) that part of a sound contains multiple frequency components, we could guess that the scene description should include an event at that time. However, the scene description often remains ambiguous given local evidence. For instance, the frequency components in question could be generated by multiple simultaneous whistle events or by a single harmonic event, and a pattern recognition system might propose both explanations as possible. To decide between these explanations we might need to consider the preceding context. These considerations motivate the analysis-by-synthesis approach. Analysis-by-synthesis incorporates the strengths of bottom-up pattern recognition to rapidly propose events to explain the sound, while precisely comparing the probability of these explanations using Bayesian inference over the full scene.

Specifically, we use a neural network to propose local event variables, which are then combined into global scene hypotheses (Figure 4, left). These steps comprise the bottom-up aspect of analysis-by-synthesis. The “event proposal” network is trained on samples from the generative model (47). The scene hypotheses are then assessed top-down with stochastic variational inference in the differentiable generative model (Figure 4, center) (48, 49), which provides a precise estimate of the posterior mass. Hypotheses are built up sequentially in time, with a set of the most promising intermediate hypotheses at each round expanded upon to explain successive intervals of sound (Figure 4, right). The details of the full inference process are given in the Methods.

**Figure 4.**
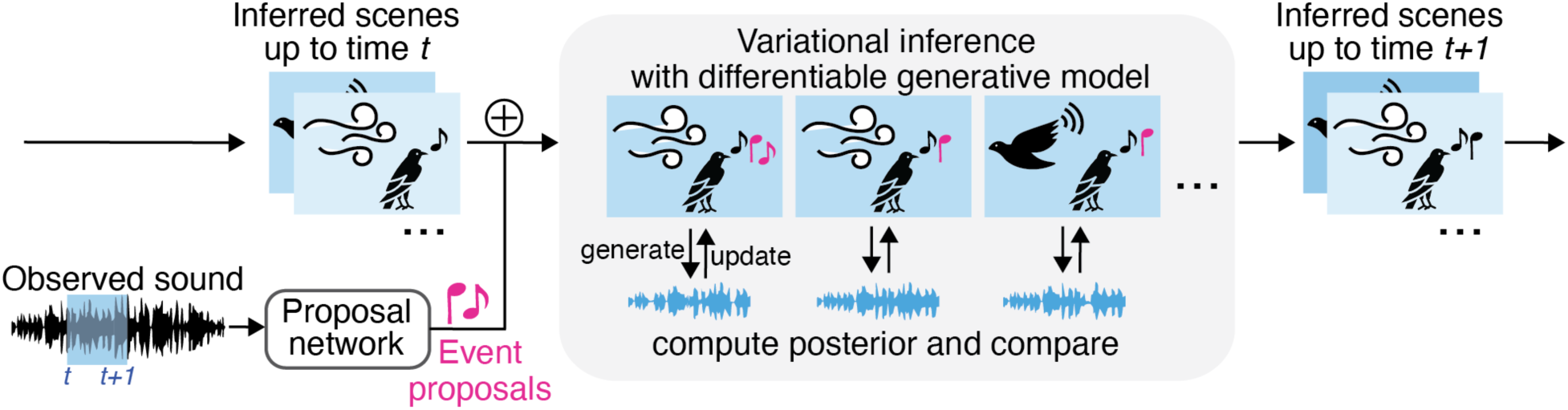
Analysis-by-synthesis inference overview. Given an observed sound, we infer a distribution over possible scenes that are likely to have generated the sound (“Inferred scenes up to time t„). This inference process proceeds sequentially, considering increasingly longer durations of audio, at each step combining a bottom-up “analysis” step with a top-down “synthesis” step. First, a deep neural network proposes events from the sound, and these events are combined into sources to create scene hypotheses (bottom-up). For each hypothesis, variational inference through the fully differentiable generative model is used to update the scene to maximize the prior and likelihood (top-down). Last, the probabilities are compared to find the best scenes given the observed sound.

This procedure results in a set of scene descriptions which best explain the full observed sound. The number of sources and events, the sound type of each source, the source parameters, and the events emitted by each source are all automatically inferred. Sequential inference is therefore general-purpose and can be applied to any audio signal, including both classic illusions and everyday sounds.

### Model results on classic auditory scene analysis illusions

The model was first evaluated on a wide range of classic auditory scene analysis illusions (Supplementary Table 1). We used the model to simulate the experiment associated with each phenomenon. Some illusions simply involve a subjective judgment of perceptual organization; in such cases, we compared the commonly reported percept with the highest probability scene hypotheses under the model (found through sequential inference). For other illusions, we simulated published psychophysical experiments for comparison with human judgments. The experiments in question often queried participants about a limited set of specific scene descriptions (e.g., in a two-alternative forced choice paradigm). When such experimental constraints were present, we replaced bottom-up search by enumeration over the experimentally defined hypotheses (“enumerative inference”). We evaluated the probabilities of each of these hypotheses with variational inference to yield an experimental response. We provide an online repository with example experimental stimuli and model inferences.^1^

**Methods Table 1.**
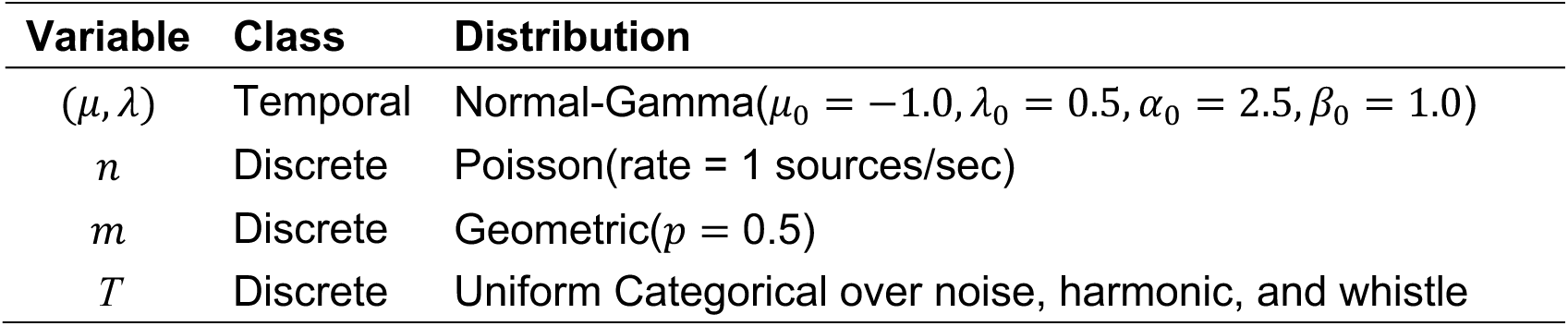
Temporal normal-gamma meta-source parameters and discrete prior parameters. The same temporal source prior is used for the rest and duration parameters.

Overall, the model qualitatively and often quantitatively reproduced human-like perception in each of the illusions. To our knowledge this provides the first demonstration that a single theory can account for this diverse set of phenomena. In the following sections we present the results for each illusion.

### Masking and filling-in

One class of illusions we tested involve “filling in”. If a less intense sound is played concurrently with a more intense sound, in some conditions the less intense sound will not be heard; this everyday occurrence is termed “masking” (65). In many such cases, the addition of the less intense sound does not alter the peripheral auditory representation to a detectable extent, rendering it undetectable. However, the perceptual interpretation can be modulated by context. For instance, a noise flanked by tones could equally well consist of two short tones adjacent to the noise, or a single longer tone overlapping the noise, that happens to be masked when the noise is present (Figure 5A). Human listeners hear this latter interpretation as long as the noise is intense enough to have masked the tone were it to continue through the noise (66). Listeners thus perceptually “fill-in” the sound that is plausibly masked, yielding a stable percept, despite physical interruption by the masker. We asked whether the model reproduces human-like filling in across a variety of contexts, inferring events that are not explicit in the sound when evidence is consistent with masking.

**Figure 5.**
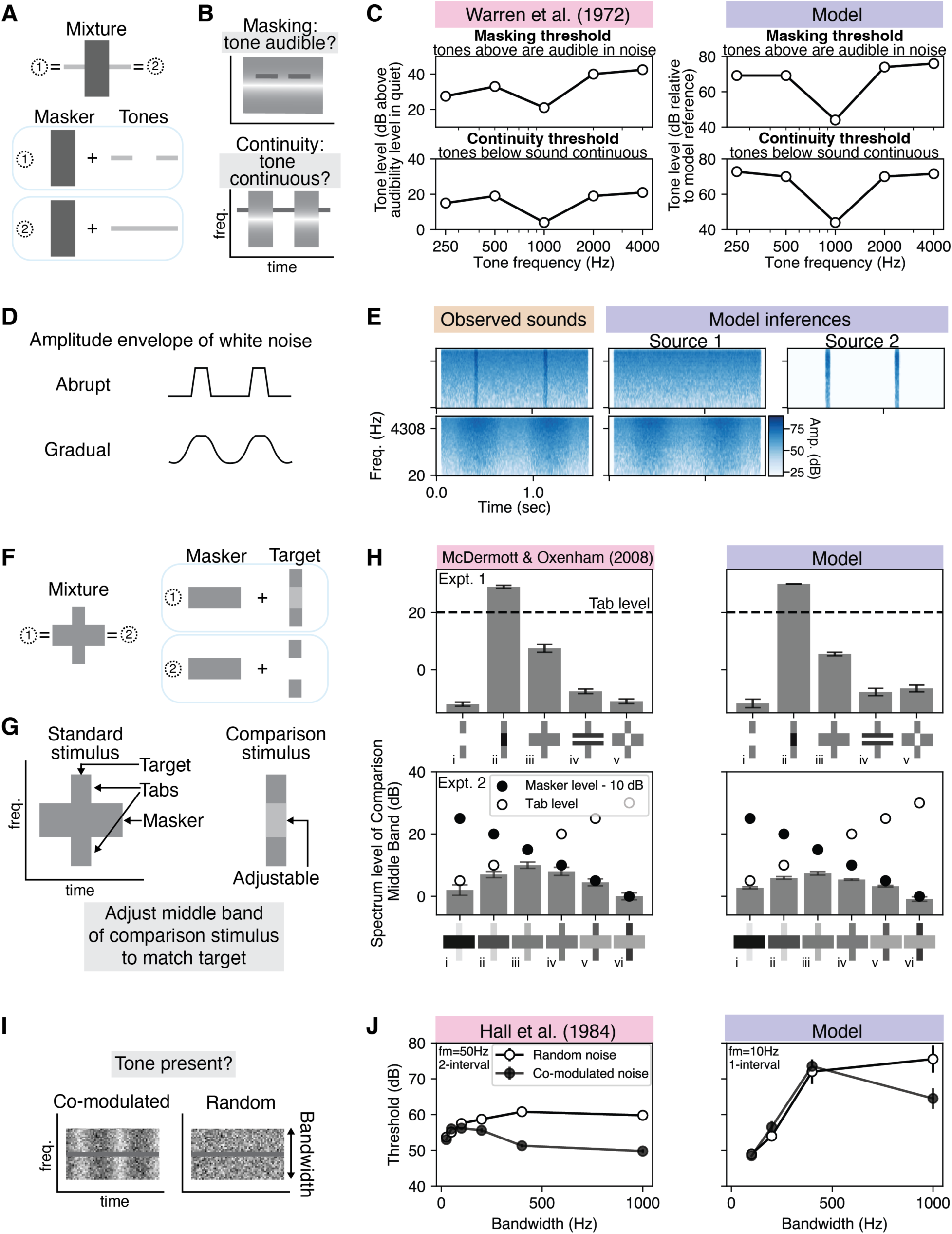
Model results for masking and filling-in illusions. A) A noise flanked by tones could be generated by summing the noise with two short tones adjacent to the noise, or a single longer tone overlapping the noise. B) Continuity illusion experimental stimuli and tasks. Thin lines: tones, grey rectangles with gradient: noise with notch at 1000 Hz. C) Left: average results for 15 human listeners from (66). Right: average model results. Like human listeners, the model’s thresholds drop at the notch. Decibel reference levels are different for humans and the model. For humans, the reference level is the level at which the tone is audible, whereas for the model it is an arbitrary value (model cochleagram is thresholded at 20 dB; for simplicity we did not include a middle ear transfer function and other factors necessary to replicate human audibility). D) Homophonic continuity. Amplitude envelope for each stimulus from (67). E) As in human perception, the model infers two sources when the amplitude envelope changes abruptly. F) A short noise target masked by a longer noise could be generated by a noise target with or without energy in the middle frequency band. Darker shade of grey indicates higher spectrum level. G) Spectral completion experiment. A comparison stimulus was adjusted until it sounded like the target in the standard stimulus, providing an indication of the target source energy inferred by human listeners. H) Left: human results from (68), averaged over 8 participants. Right: average model results. The model shows a similar pattern of results as human listeners, inferring energy in the target when it could plausibly be masked. I) Co-modulation masking release experiment. A tone is superimposed on a noise of varying bandwidth that is either co-modulated (multiplied by a time-varying envelope) or random. Participants judged whether a tone was present or not. J) Left: average results from 5 participants, as measured in (69). Right: average model results. Like human listeners, the model exhibited lower detection thresholds in co-modulated noise for sufficiently large bandwidths. Here and elsewhere, error bars depict ±1 standard error (over participants for human results, over inference replications for model results), and model results are averaged over 10 inference replications. Error bars are missing for some human plots because they were not plotted in the original publications, some of which were published many decades ago. Error bars are sometimes not visible for model results as they are smaller than the symbol size.

#### Continuity illusion

In one classic experiment (66), tones of various frequencies were either *embedded* in noise (to measure masking) or *alternated* with noise (to measure continuity, i.e., filling in) (Figure 5B). Listeners’ masking threshold corresponded to the level at which the embedded tones became audible, and their continuity threshold corresponded to the level at which the alternated tones sounded continuous. The noise was missing energy in a “notch” around 1000 Hz in order to test whether overlap in the frequency domain was necessary for listeners to perceive continuity. The results showed a close correspondence between the two types of thresholds across tones of different frequencies (Figure 5C, left). Moreover, the drop in masking and continuity thresholds for the 1000 Hz tones confirmed the importance of overlap in frequency between the tone and the masking noise. Both effects replicated in our model (Figure 5C, right).

#### Homophonic continuity

The model was also tested on a classic variant of the continuity illusion involving amplitude modulated noise (Figure 5D). When an initially soft noise undergoes a sudden rise in intensity, listeners perceive the initial source as continuing unchanged behind a distinct, louder noise burst (66). In contrast, if the amplitude modulation occurs gradually and reaches the same peak, listeners instead hear a single source changing in intensity. In accordance with human listeners, the model explained the noise as two sources when its amplitude changed abruptly (1 ms ramp), and one source when its amplitude changed gradually (250 ms ramp) (Figure 5E).

#### Spectral completion

An analogous illusion occurs over the frequency spectrum, dubbed “spectral completion” (Figure 5F). In (68), listeners heard a long masker noise, which overlapped with a brief target noise halfway through its duration (Figure 5G, left). The spectrum of the target was ambiguous because the middle band of its spectrum could be masked. Listeners were asked to adjust the level of the middle band of a comparison noise (Figure 5G, right), until the comparison perceptually matched the target. Listeners chose the level of the middle band to be well above its audibility threshold, suggesting that they perceptually filled in the middle portion of the target. Several variations on this basic stimulus configuration revealed how this perceptual “filling-in” was affected by context and masker levels (Figure 5H, left). The pattern of judgments across these stimuli was replicated in our model (Figure 5H, right).

#### Co-modulation masking release

Another masking-related grouping phenomenon occurs when masking noise is co-modulated (69). Coherently modulated noise produces lower tone detection thresholds than unmodulated noise, “releasing” the tone from masking (hence “co-modulation masking release”). The effect can be measured by comparing detection thresholds for co-modulated relative to unmodulated noise maskers as the masker bandwidth widens (Figure 5I). In contrast to unmodulated noise, for which thresholds grow and then level off (at the “critical band”) as bandwidth increases (70), co-modulated noise produces thresholds that decrease for sufficiently wide bandwidths (71) – the high and low frequencies of the noise evidently group with the noise frequencies around the tone, helping listeners perceptually separate the tone and the noise. The model shows this same qualitative effect, although the bandwidth at which the effect becomes evident is higher than that for human listeners (Figure 5J; we used a lower modulation cutoff than in the human experiment due to the limited resolution of the model’s cochleagram representation).

### Simultaneous grouping

Another set of classic illusions pertain to whether simultaneous sound components are perceived as part of the same source. We tested whether the model could account for simultaneous grouping illusions involving harmonic frequency relations and common onset (arguably the two most commonly cited grouping cues).

#### Frequency modulation

Modulating the frequency of a subset of tonal components in parallel causes them to perceptually separate from an otherwise harmonic tone (72). To test whether the model showed a similar effect, we replicated a classic illusion in which the odd-numbered tonal components had a constant fundamental frequency and the even-numbered components were frequency-modulated in parallel (Figure 6A). The model explains this sound with two distinct harmonic sources, replicating human perception (Figure 6B) (72).

**Figure 6.**
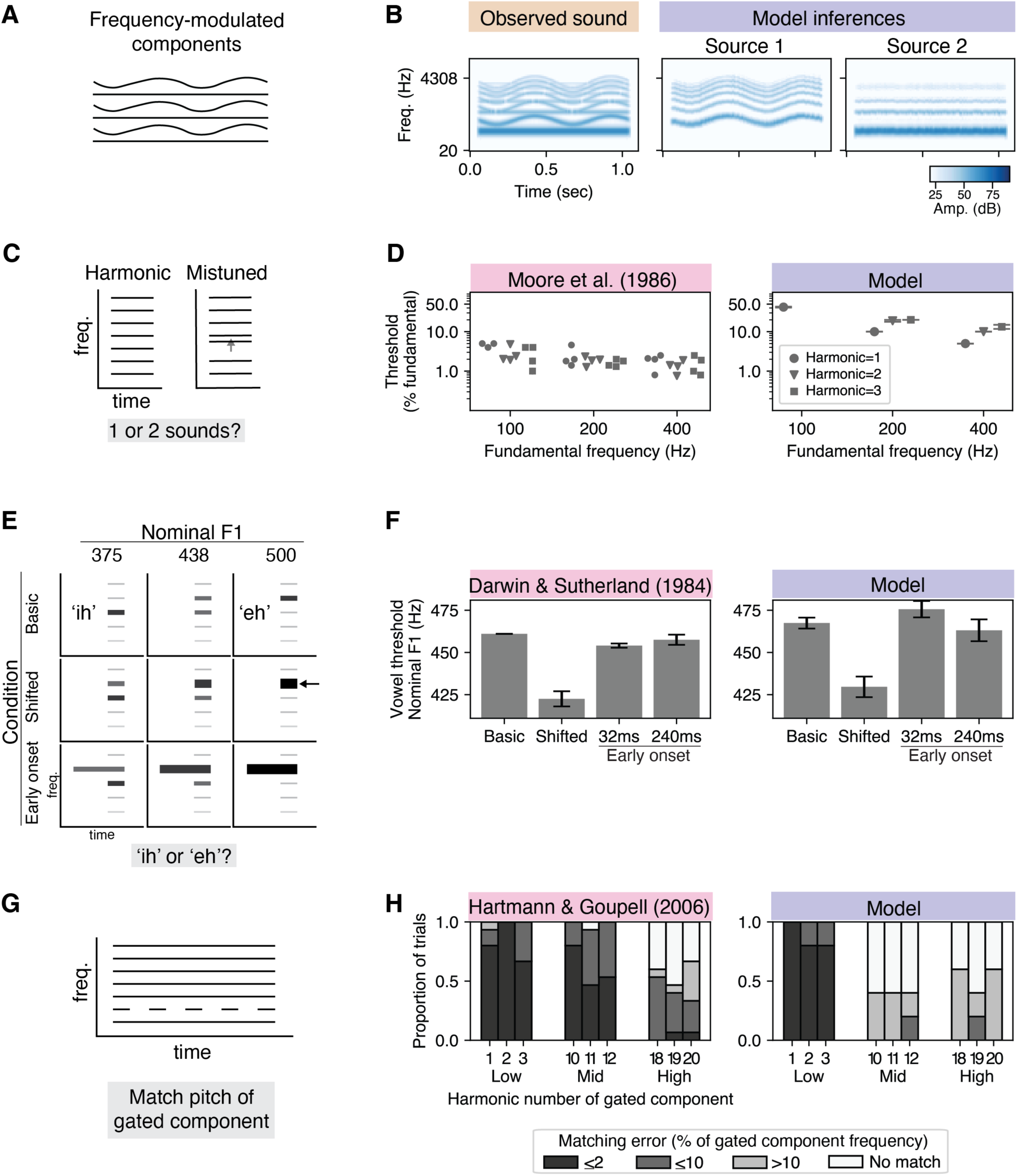
Model results for simultaneous grouping illusions. A) Frequency modulation. The stimulus is an otherwise harmonic complex tone whose even harmonics are coherently frequency modulated. Humans perceive the even and odd harmonics to belong to distinct sound sources. B) Like humans, the model infers the modulated and unmodulated components to belong to separate sources. C) Harmonic mistuning. Listeners heard an otherwise harmonic complex tone, one harmonic of which could be mistuned, and judged whether they heard one or two sounds. D) Left: mistuning detection threshold as a percent of the fundamental frequency. Each dot plots the threshold of one of the four participants in (76). Right: average model thresholds. In both human and model results plots, missing dots indicate that a threshold could not be measured below the maximum possible value of 50% (in humans, this happened in one participant for the 1^st^ harmonic of 100 Hz; in the model, this happened for the upper harmonics of 100 Hz). The model shows a mistuned harmonic effect, albeit with higher thresholds than human listeners. The higher thresholds plausibly reflect the limits of the frequency information available in the cochleagram used for inference. Here and elsewhere, error bars depict ±1 standard error, with model results averaged over 10 inference replications. E) Onset asynchrony. The schematic shows example stimuli from the experiment. The “basic” stimuli are created by shifting the first formant of the vowel from 375 to 500 Hz. The “shifted” and “early onset” stimuli are both created by adding a 500 Hz tone (at arrow) to the basic stimuli, but with different onsets (synchronous and 32 or 240 ms earlier, respectively). Participants judged whether the vowel sounds like “ih” or “eh”. F) Left: vowel boundaries averaged over 6 human listeners from (77). Right: average model vowel boundaries. In both human listeners and the model, the boundaries are lower in the shifted conditions than in the basic condition, but are restored by the onset asynchrony, indicating that the added harmonic is grouped with the harmonic tone when synchronous, but segregated as a distinct source when asynchronous. G) Cancelled harmonics. Schematic shows an example stimulus with the 2^nd^ harmonic gated. H) Left: the distribution of frequency match errors for stimuli with different gated harmonics, as measured across 3 participants in (79). To assess the match error for the model, we compare the frequency of the inferred whistle source (if present) to the frequency of the gated harmonic (see Methods). The model replicated human performance for low numbered harmonics, and showed a similar trend towards higher error at higher-numbered harmonics.

#### Mistuned harmonic

Listeners tend to hear tonal components with harmonically related frequencies as a single perceptual entity. When the frequency of one tonal component deviates from the harmonic series, the component is heard to stand out as a separate tone, demonstrating grouping via harmonicity (73–75). This effect can be quantified as the threshold at which this separation occurred, expressed as a percent of the harmonic frequency (76) (Figure 6C). Across multiple fundamental frequencies and harmonic numbers, the model also showed a measurable mistuned harmonic effect (the mistuned harmonic was reliably inferred as a separate “whistle” source; Figure 6D, right), albeit with higher thresholds than expert listeners in the experiment (plausibly due to the limited frequency information in the cochleagram representation used to compute the likelihood).

#### Asynchronous onsets

Another type of grouping illusion involves “cue conflicts”, for instance with harmonicity favoring grouping and onset asynchrony favoring segregation. To assess whether source models could provide the basis for integrating such cues, we asked whether the model resolves cue conflicts in the same way as human listeners.

One classic experiment used judgments of vowel quality (in particular, whether a sound was perceived as /I/ or /e/) to assess whether a frequency component of a harmonic sound was perceptually grouped with the others (77). Vowels differ in the frequencies at which the spectral envelope exhibits peaks (“formants”): the harmonic tone perceived as the vowel /I/ has a lower first formant than /e/. A stimulus was constructed such that an added frequency component would shift the vowel’s formant if it were grouped with the rest of the vowel, thereby changing its category, enabling vowel categorization judgments to be used to assess grouping.

A *basic* continuum of seven stimuli was constructed by progressively increasing the first formant of a 125 Hz harmonic tone from 375 Hz to 500 Hz. This continuum was perceived as morphing from /I/ to /e/ (Figure 6E, first row). A new *shifted* continuum was then constructed by adding a 500 Hz pure tone to each harmonic tone of the basic continuum (Figure 6E, arrow in second row). The phoneme boundary occurred earlier in the shifted continuum (because the added tone effectively raised the first-formant valueof the aggregate sound) compared to the basic continuum. Last, two *early-onset* continua were constructed. These continua were constructed by adding a 500 Hz pure tone to each stimulus of the basic continuum, but with the pure tone onset occurring 32 ms or 240 ms before that of the harmonic tone (Figure 6E, third row). Even though the frequencies and amplitudes were physically identical to those of the shifted continuum, listeners perceived the early-onset continua to have a similar phoneme boundary as the basic continuum (Figure 6F, left). These results indicated that the pure tone was not integrated with the harmonic tone when their onsets differed. To test this effect in our model, we added a final vowel categorization step after inference, based on empirical vowel formant distributions (78) (see Methods). Our model replicated the variation of the phoneme boundaries across conditions (Figure 6F, right), indicating human-like interactions between common onset and harmonicity on grouping.

#### Cancelled harmonics

The “cancelled harmonics” illusion (80) (Figure 6G) provides another example of cue conflict: for the duration of a harmonic tone, one of its components is gated off and on over time. Although this gated component remains harmonically related to the rest of the tone at all times, it perceptually segregates as a separate sound. To quantify this effect, listeners were asked to match the frequency of different gated harmonics within a complex tone (79). Human participants could closely match the frequency of the gated component up to the tenth harmonic (Figure 6H, left). The model qualitatively replicated this effect, as evidenced by inferring a whistle source that was well-matched in frequency to the gated harmonic for low harmonic numbers (Figure 6H, right). The quantitative discrepancy (whereby humans, but not the model, could accurately match the frequency of middle harmonics) again seems plausibly explained by limitations in the frequency information available in the cochleagram representation used for inference.

### Sequential grouping

A final set of auditory scene analysis illusions concern how sound events are grouped or segregated into sequences over time.

#### Frequency proximity

One classic demonstration of the role of frequency proximity in sequential grouping can be found in interleaved rising and falling tone sequences, producing the “X” pattern apparent in Figure 7A (81). Humans typically do not hear rising or falling trajectories when presented with the mixture. Instead, we hear the higher frequency tones as segregated from the lower frequency tones, producing two sequences that “bounce” and return to their starting points. By contrast, when the pure tones in the rising trajectory are replaced by harmonic tones, humans hear the crossing explanation (Figure 7A). The model replicates these findings, explaining the pure tone sequence as bouncing, and the sequence with harmonic tones as crossing (Figure 7B).

**Figure 7.**
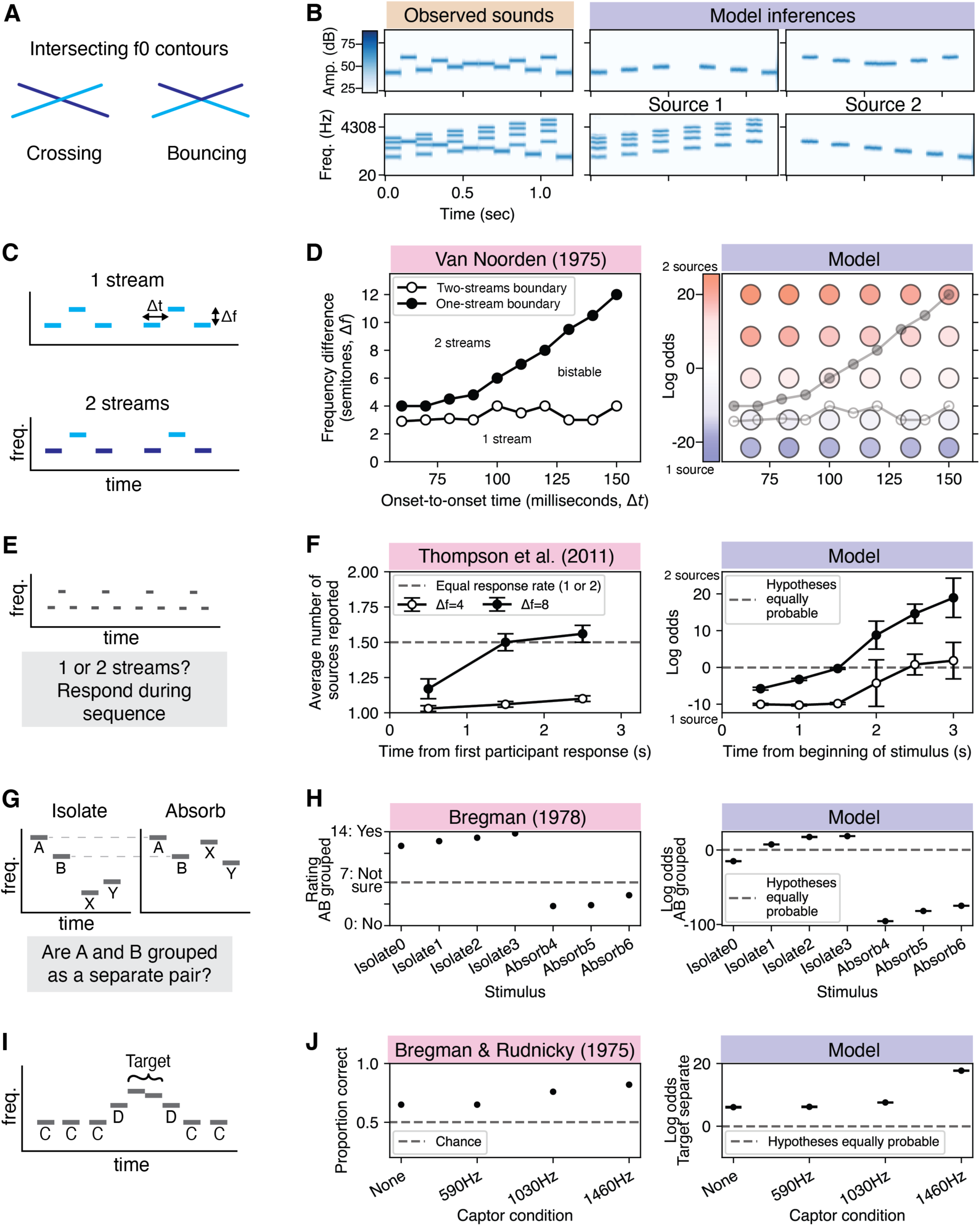
Model results for sequential grouping illusions. A) Frequency proximity. Schematic shows possible perceptual organizations for interleaved tones (81). B) The model prefers “bouncing” sources (row 1), unless there is a spectral difference between the tones in the upward and downward sweeps (row 2), matching human perception. C) The two perceptual organizations of the triplet tone sequence that were queried in (82) and (84). In the two-streams percept, listeners hear the low tones grouped together as separate from the high tones, creating two isochronous sequences. In the ‘one-stream’ percept, listeners hear all tones grouped together, creating a sequence with a galloping rhythm. D) Bistability. Left: listeners’ two-streams boundary (threshold below which listeners no longer heard two streams) and one-stream boundary (threshold above which participants no longer heard one stream) from (82). Right: each dot is average model log odds at that stimulus setting (red=prefers two sources, blue=prefers one source). The region of bistability for human listeners overlaps with the region where the model log odds are close to zero (indicating bistability in the model posterior). The human data is superimposed for reference (no model boundaries are drawn). Here and elsewhere, model results are averaged over 10 inference replications. E) Build-up of streaming. Listeners heard a triplet tone sequence as depicted in C (with a frequency difference f of four or eight semitones). Listeners could indicate at any time during the sequence if they heard one or two streams. F) Left: average responses from (84), showing that participants more often report hearing two sources with more repetitions of the stimuli. Right: average model log odds for the two-sources versus one-source explanation (positive: prefers two-sources, negative: prefers one-source), showing the same preference for two-sources with more repetitions. Here and elsewhere, error bars depict ±1 standard error. G) Effects of context, part 1. For each stimulus condition, A and B tones have the same frequency and timing, with only the X and Y tones changing across conditions. In the ‘isolate’ conditions, X and Y occupy a different frequency range to A and B. In the ‘absorb’ conditions, they occupy an overlapping frequency range. H) Left: average participant ratings for each of 7 conditions from (14). 0 indicated a confident judgment that A and B could not be heard as a separate pair, and 14 indicated a confident judgment that A and B could be heard as a separate pair. Right: average model log odds (positive=prefers A and B in their own source, negative=prefers A and B in separate sources). The log odds are more positive for the isolate stimuli, mirroring their higher ratings by human listeners. I) Effects of context, part 2. Schematic tone sequence (C: captor tones, D: distractor tones at 1460 Hz). J) Left: average participant accuracy on an interval judgment task (85). Right: average model log odds (positive=prefers target in own source, negative=prefers target grouped with distractors). The model log odds increases as the captors get closer in frequency to the distractors, similar to the effect on human accuracy.

#### Bistability

The best-known demonstration of bistability in auditory perceptual organization is arguably the classic “ABA sequence” (82) (Figure 7C). This sequence comprises a repeating set of three tones in which the first and last tone have the same frequency. Depending on the stimulus parameters, participants typically hear one of two dominant perceptual organizations: 1) all the tones are grouped together to produce a galloping rhythm (“one stream”), and 2) the A tones are grouped separately from the B tones, such that there are two sequences which each produce an isochronous rhythm (“two streams”). Increasing the frequency difference between A and B (Δ*f*), and decreasing the temporal rate of tones (Δt), both increase reports of the “two streams” organization (Figure 7D, left) (82). In the low Δ*f* region, the “one stream” percept is dominant, while for high Δ*f* and low Δt, the “two streams” percept prevails. For intermediate settings of these variables, the percept is bistable. The model accounts for these trends, as reflected in the log odds of the two explanations (the logarithm of the ratio of the probability of the two interpretations, displayed as red vs. blue in right panel of Figure 7D). The model log odds are close to zero in the intermediate region, reflecting bistability in the model posterior.

#### Buildup of streaming

Another factor affecting the preferred perceptual organization of the ABA sequence is the number of repetitions. When an ABA triplet is repeated multiple times over several seconds, listeners increasingly tend to hear two streams (83). We simulated an experiment in which listeners heard an ABA sequence and could indicate whether they heard one or two sources at any time during the sequence (Figure 7E) (84). Participant responses were averaged and binned to reveal an increase in two-stream responses as participants heard more repetitions of the triplet over time (Figure 7F, left). This trend is replicated by the model, where the posterior odds increase over time to favor “two streams” over “one stream” (Figure 7F, right).

#### Effects of context

Streaming can also be influenced by context. We simulated two complimentary experiments that show the influence of the surrounding context on whether the tones are grouped or not (Figure 7G–J) (14, 85).

In the first experiment, the frequency separation and timing between tones A and B were held constant, but they could be perceived as the same or different stream depending on the frequencies of context tones X and Y (14) (Figure 7G). There were two kinds of tone sequences: ‘isolate’ sequences where X and Y were in a separate frequency range from A and B, and ‘absorb’ sequences where all tones were in the same frequency range. Listeners rated whether they heard tones A and B as a separate pair grouped in their own stream for four ‘isolate’ sequences and three ‘absorb’ sequences; grouping into a pair was more likely in the ‘isolate’ conditions (Figure 7H, left). The model log odds compare the hypothesis when A and B are in their own source versus when they are in separate sources. The log odds differentiated the two types of sequences, qualitatively replicating listeners’ ratings (Figure 7H, right).

In the second experiment, the frequency of ‘captor’ tones affected whether listeners heard ‘distractor’ tones group with ‘target’ tones (85) (Figure 7I). Listeners judged whether the two target tones moved upward or downwards in pitch. They were more accurate when the frequency of the captor tones was close to the frequency of the distractor tones (at 1460 Hz), presumably because the distractor tones then group with the captor tones, making it easier to hear out the target tones (Figure 7J, left). We evaluated the tendency of the model to group the target and distractor tones, by measuring the model log odds comparing the hypothesis that the target tones are in their own source versus that the distractor and target tones are grouped together. The pattern of log odds across the captor conditions qualitatively mirrored the increase in accuracy for human listeners (higher for higher frequency captors; Figure 7J, right).

## Model comparisons

Figures 5-7 show that our model comprehensively replicates many classic illusions in auditory perceptual organization. However, it remains unclear what aspects of its structure are important for these results. We address this question through a series of model comparisons. First, we test “lesioned” versions of our model in order to clarify their role in obtaining these results. Second, we demonstrate the difficulty of matching human perception by evaluating an alternative model class that lacks highly structured constraints – deep neural networks trained on recorded audio (the most widely used alternative model class that can be applied to actual audio).

### Model lesions

To assess the contribution of the structure imposed by the generative model, we devised a set of model “lesions” that targeted different levels of the model’s structure. We ran each lesioned model on the full set of illusions shown in Figures 5-7. As an aggregate measure of how well each model reproduced human-like perception of the set of illusions, we calculated the dissimilarity between model and human judgments for each illusion, and then averaged this dissimilarity across illusions (see Methods). The first two lesions addressed the hierarchical priors over sources, while the second two addressed the event priors.

#### Fixed source parameters

To test whether the generative model’s hierarchical source priors were necessary, we removed the temporal source priors (Figure 3A) and the variance and lengthscale source priors over fundamental frequency, amplitude, and filter shape (Figure 3B). Instead of those distributions, each source was assumed to have the same parameters (fixed to each distribution’s mode). We found that restricting the variability in possible source parameters increased the model’s dissimilarity to human perception (Figure 8A, fixed vs. ours; p<0.01), affecting several illusions including grouping by frequency modulation and context effects in sequential grouping (Supplementary Table 2). This result suggests that an expressive distribution over sources is important to explain human perceptual organization.

**Figure 8.**
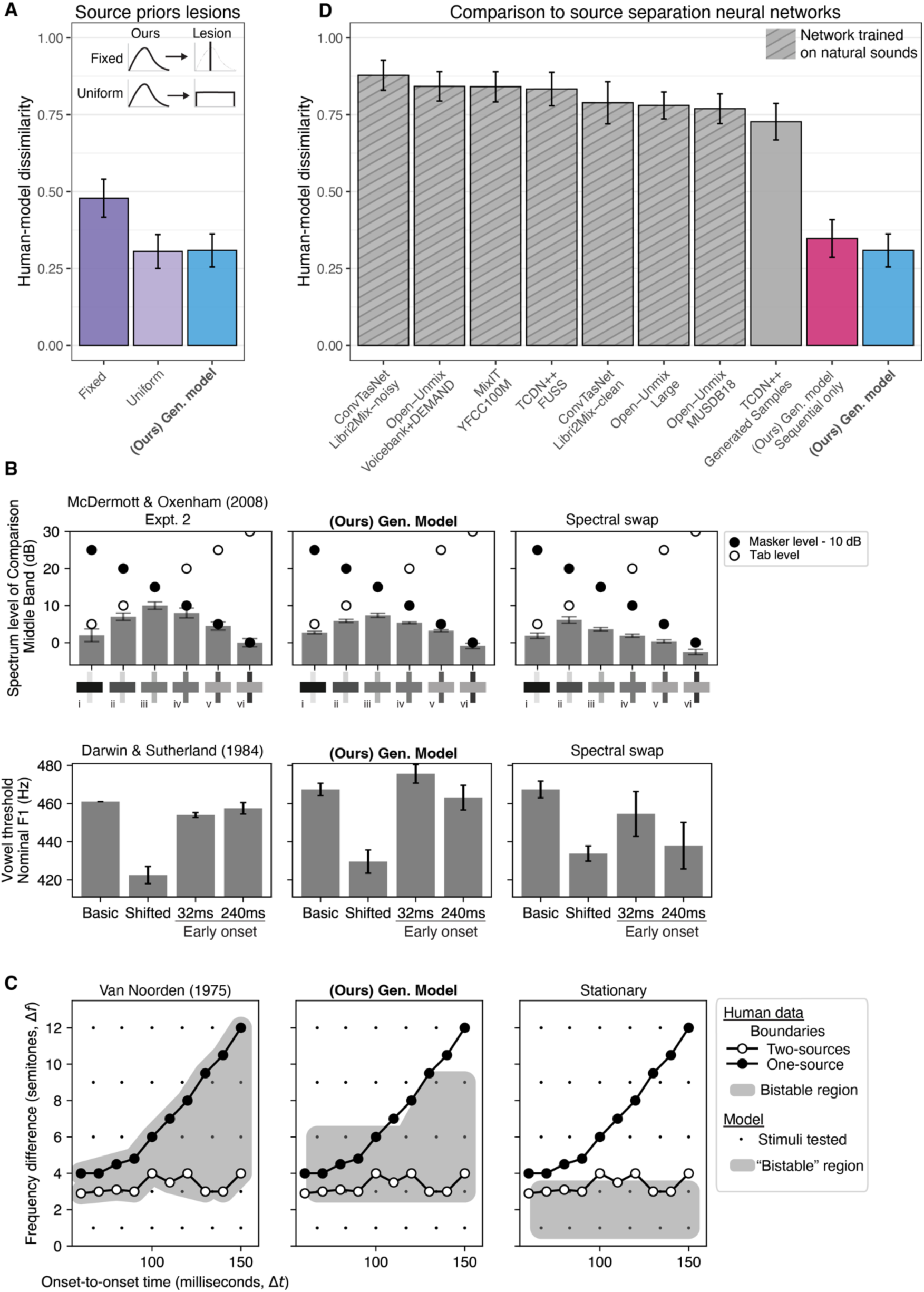
Comparisons with lesion models and neural networks. A) Human-model dissimilarity for models with alternative source priors as compared to our model. A value closer to zero indicates greater similarity to human perception. The blue bar corresponds to our main model (evaluated with both enumerative and sequential inference, as described in “Model results on classic illusions”). Fixing all sources to the mode of the source priors (Fixed) decreased dissimilarity more than changing the source priors to uniform distributions (Uniform; which increases the variability in the prior over sources). Here and in D, error bars plot ±1 standard error computed by bootstrap (see Methods). B) Example aberrant results from the model with swapped covariance kernels for harmonic and noise sources. For this alternative model, the spectral completion and onset asynchrony experiments did not qualitatively match the human results, plausibly because both involve spectral filling-in. C) Example aberrant result for a lesioned generative model with a stationary covariance kernel for frequency and amplitude trajectories. The original results of (82) are compared to those of our intact model as well as this lesioned model. Human boundaries are overlaid on each plot, with conventions following Figure 7D. Small black dots indicate stimulus parameters for which we obtained model judgments. For each model, the shaded area (‘bistable’ region) covers the stimulus parameters for which the log odds fell between -15 and 15. For all stimulus parameters, the lesioned model favours the “two-streams” explanation more than the original model. By contrast, the bistable region of the intact generative model overlaps with that for humans. D) Human-model dissimilarity for source separation neural networks as compared to our model. The hatched grey bars correspond to neural networks trained on source separation tasks using corpora of natural sounds (labelled as network/dataset). The solid gray bar, ‘TCDN++/Generated Samples’, shows the dissimilarity for a source separation network trained on samples from our generative model. We also assessed our model using purely sequential inference to obtain judgments in the same way as for the networks (pink solid bar). Our model remained more similar to human perception in this case.

**Methods Table 2.**
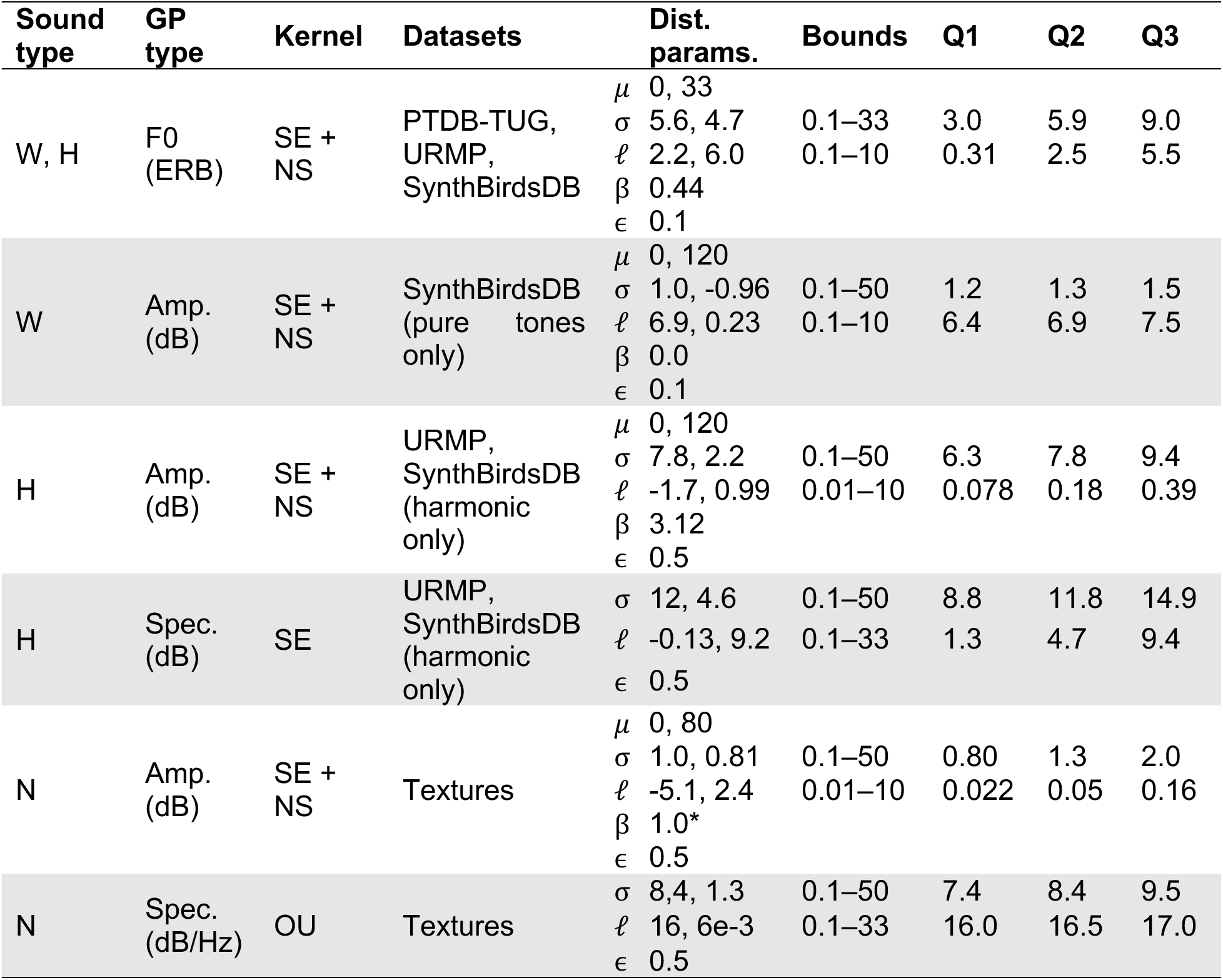
Gaussian process meta-source parameters. The mean *μ* is uniformly distributed (with units listed under GP type). The inverse softplus of the standard deviation *σ* and lengthscale ℓ are each normally distributed. Samples of *σ* and ℓ are bounded to reasonable values to avoid overflow or underflow errors (i.e., they were sampled from a truncated distribution). *σ* is in the GP-units, while ℓ is in seconds for excitation variables and in ERB for spectrum. *β* is a positive constant that implements the non-stationary kernel. As is typical for GPs, *∈*^2^ is added to the diagonal of the covariance matrix for numerical stability. The last three columns indicate the first, second, and third quartile of sampled *σ* and ℓ distributions based on 5000 samples. *Note: We manually set the value of *β* to 1.0 because the scenes in the textures dataset only had one event.

#### Uniformly distributed source parameters

The generative model’s variance and lengthscale source priors were originally fit to a small set of everyday sounds. As a result, sources with excitation trajectories and spectral shapes whose variances and lengthscales match those sounds are more probable under the model. The second model lesion changed these source priors to be uniform, making a wider range of source parameters equally likely and therefore making a wider range of excitation trajectories and spectral shapes equally likely. In contrast to the Fixed lesion, this Uniform alteration did not increase the model’s overall dissimilarity from human perception (Figure 8A, uniform vs. ours; p=0.30 by empirical bootstrap). This result indicates that the model’s similarity with human perception is driven more by the model’s overall structure (e.g., events with discrete onsets/offsets, smooth amplitude trajectories) than by the fine-tuning of the source priors to everyday sounds.

#### Spectral swap and stationary covariance

Two other model lesions tested the role of the event priors for fundamental frequency, amplitude, and spectrum (Figure 2B, events). We altered the covariance kernels of the Gaussian processes, which encode tendencies in how spectral shape (the filter) changes over frequency and how fundamental frequency and amplitude (the excitation) change over time.

In the original model, we used different priors for the filters of different sound types (Figure 3C). These were chosen to cause spectra of harmonic sources to be smoother than those of noise sources, based on the observation that some noise sources have filters that change abruptly with frequency (see Methods: Equation 19). We swapped the kernels underlying these priors to examine their impact. We found this swap reduced the model’s match to human perception for two illusions: spectral completion and onset asynchrony (Figure 8B). These illusions both involve the inference that one sound masks part of another sound’s spectrum, with spectral “filling-in” occurring as a result. However, one involves a noise source while the other involves a harmonic source. The effect of the kernel swap suggests that filling-in may depend on relatively specific assumptions about the spectra of different types of sounds.

The final lesion involved the covariance kernels for the excitation trajectories. In the original model this kernel included a non-stationary term (see Methods: Equation 15), intended to enable the model to express a single source that modifies its sound-generating process between events, even if these events occur in immediate succession (e.g., a dog panting in and out; Supplementary Figure 1). When this non-stationary term was removed, we found a reduced match to human perception for sequential grouping illusions: this lesioned model consistently favored two streams even when human listeners do not (Figure 8C). This result highlights one possible function of the event level priors in explaining perceptual organization, namely, representing discontinuities in sound-generating processes.

In summary, these lesions demonstrate the necessity of the different components of the generative model. Comprehensively explaining many perceptual illusions in a single model required a hierarchical, expressive model of sources and events. The model also provides a refined understanding of key assumptions about sound generation, as highlighted by the kernel lesions.

### Neural network source separation comparisons

We next evaluated an assortment of contemporary source-separation networks on the set of classic auditory scene analysis illusions in the previous section. This type of model is arguably the most widely used alternative model class. In addition, since previous work has demonstrated that deep neural networks can reproduce key aspects of human behaviour and brain representations in tasks such as sound localization, pitch perception, and word recognition (86–89), such models might reasonably serve as a baseline for matching human judgments. We used seven published models trained on mixtures of natural sounds, chosen to span a diversity of architectures, tasks, supervised and unsupervised training regimes, and natural sound datasets (90–95).

Rather than inferring latent symbolic scene structure as in the generative model, source separation networks take in a mixture waveform and output ‘premixture’ waveforms. For each network, we obtained its premixture estimates for each experimental stimulus. To run a network on a psychophysical task requiring a choice between alternative scene descriptions, we took its judgment to be the scene description whose rendering produced the minimum mean squared error from the networks’ output, as measured on a cochleagram (see Methods).

To evaluate our model in the same way that we evaluated the source separation networks, we inferred sources for all the experimental stimuli (rather than estimating the probability of the experimenter-specified hypotheses, which is not possible using a conventional source separation model). For each experimental stimulus, we rendered source sounds from the inferred scene and analyzed them in the same way as for the neural networks.

As shown in Figure 8D, none of the source separation networks could replicate the generative model’s match to human perception across the broad range of illusions tested here (neural networks: grey bars, model: blue bar; p<0.01 for each network). This advantage was evident even when our model was evaluated in the same way as the source separation networks (without the benefit of enumerative inference; Figure 8D, pink bar; p<0.01 for each network). The generative model’s advantage was present for all three classes of illusions (filling in, simultaneous grouping, and sequential grouping; Supplementary Figure 3). These results illustrate the difficulty of explaining human perception of classic illusions – human-like results do not fall out of contemporary source separation systems trained on natural sounds.

We also assessed whether a source separation network would produce better results when trained on samples produced by the generative model (Figure 8D; gray bar without hashing). The resulting network also had a worse match to human perception than the generative model (p<0.01). This result demonstrates the difficulty of fully “amortized” inference – even if a generative model is used for training, it is difficult to replicate perception exclusively using a contemporary neural network, highlighting the utility of our analysis-by-synthesis approach. It also suggests that the dissimilarity of the source separation networks to human perception is not wholly explained by the data these networks were trained on.

## Model results on everyday sounds

Unlike many previous models of auditory illusions, our generative model can be applied to any sound waveform, allowing us to test it on everyday sounds. This allowed us to ask: can the same generative principles that explain auditory scene analysis illusions also explain the perceptual organization of naturalistic sounds? This question is critically important for linking classical perception research to real-world competencies – a theory of audition must be able to account for both. Because the various alternative models considered in the previous section were unable to account for the human perception of illusions, and thus ruled out as accounts of perception, we focused our investigations of everyday sounds on our main generative model.

We evaluated the generative model on naturalistic sound mixtures from the Free Universal Sound Separation dataset (FUSS) (91). As the basis for quantifying the generative model’s match to human perception, we identified four ways that the model’s inferred sources could deviate from human perceptual organization:

1. **Unrecognizability**: the model infers a source which people do not hear in the mixture
2. **Absence**: the model omits a source that people do hear in the mixture
3. **Over-segmentation**: the model segregates sounds into distinct sources when people hear these sounds as coming from a single source
4. **Over-combination**: the model combines sounds into a single source when people hear these sounds as coming from distinct sources

We aimed to understand how often and in what circumstances the generative model’s inferences deviated from listeners’ perceptual organization in each of these ways. We note that our goal was to investigate whether the model captured perceptual organization rather than whether it reproduced fully naturalistic sounds. Samples from the model were typically not fully naturalistic in appearance, but might nonetheless in principle capture the structure inferred by human listeners.

Experiment 1 addressed the first deviation type, identifying which model-inferred sources were unrecognizable to human listeners in the mixtures. Experiment 2 addressed the other three deviations: absence, over-segmentation, and over-combination. We encourage the reader to listen to the online repository of rendered model inferences.^2^

### Experiment 1: unrecognizability deviations

On each trial, participants heard three sounds (Figure 9A). They first heard a two-second sound mixture. They then listened to two potential sources and chose which was part of the mixture. In the “recorded” condition, both potential sources were recorded premixture sounds, one of which was part of the mixture presented in the trial, with the other being an unrelated premixture sound. In the “model” condition, both potential sources were inferred sources rendered via the generative model: one was a source inferred from the trial mixture, while the other was inferred from a different mixture.

**Figure 9.**
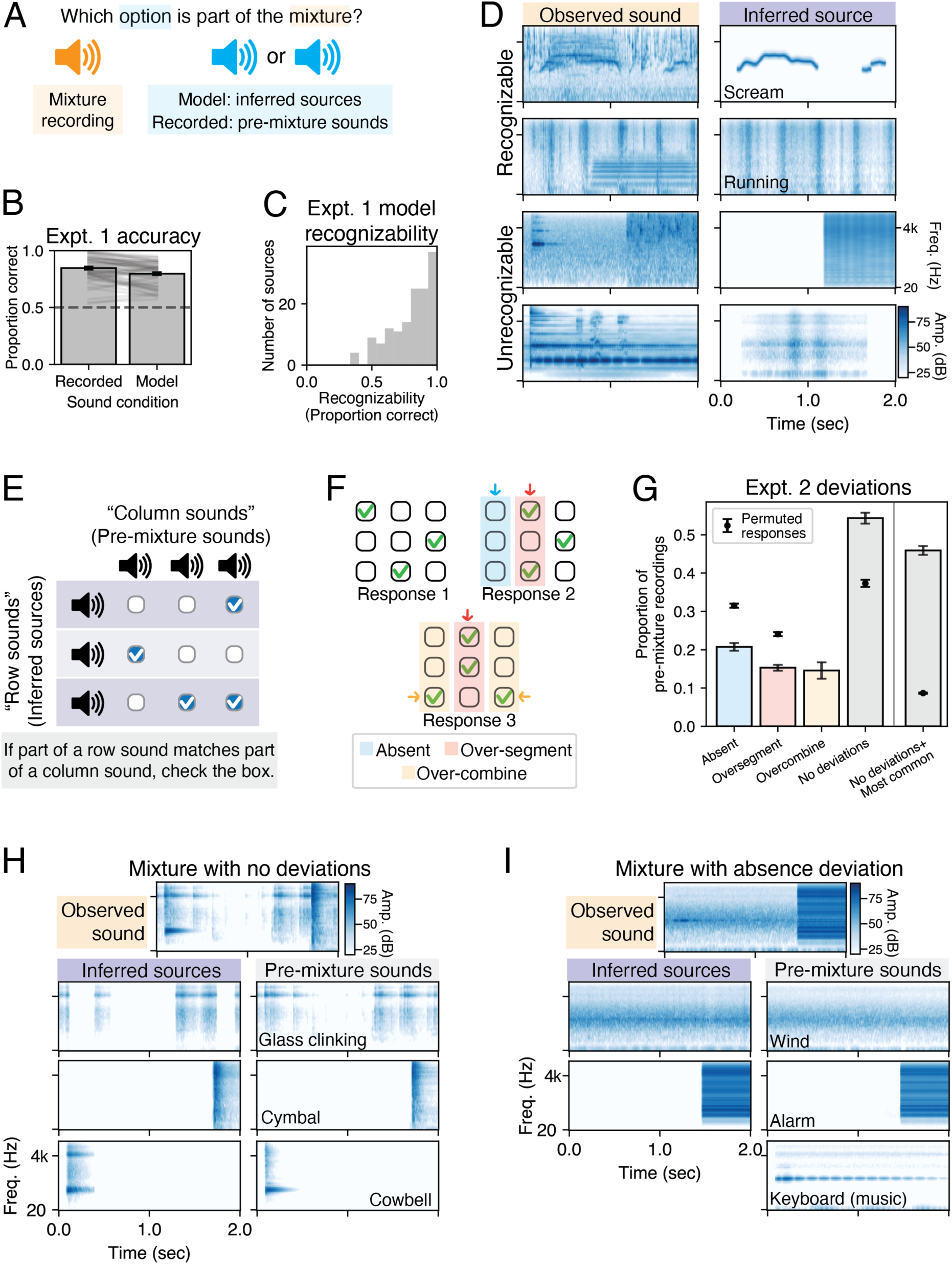
Model inferences for everyday sounds. A) Experiment 1 tested whether the model’s inferred sources were recognizable to humans as having occurred in the mixture. B) Mean results by condition for Experiment 1 (n=64). Accuracy in the model condition is well above chance. C) Histogram of proportion correct responses for each inferred source in the ‘model’ condition. Histogram is skewed left, indicating that unrecognizability errors occurred for a minority of inferred sources. D) Examples of inferred sources with high and low recognizability in Experiment 1. The premixture sounds that compose each observed sound, by row: 1) stream (water), scream, tearing; 2) running, pots and pans, brass instrument; 3) ocean waves, woodwind instrument, water from faucet; 4) organ, giggle, glass ringing. E) Experiment 2 paradigm to measure absence, over-segmentation, and over-combination deviations. F) Examples of possible responses. Response 1 has no deviations tallied for any columns (each column corresponds to a premixture sound). Response 2 has an absence deviation for column 1, an over-segmentation deviation for column 2, and no deviation for column 3. Response 3 has an over-combination deviation each for columns 1 and 3, and an over segmentation deviation for column 2. G) Mean proportion of pre-mixture recordings with each deviation type, averaged across participants (n=25). Black circles: estimates of deviations that would be expected by chance, obtained from random permutations of participant responses, constrained to have at least one checkmark per row. H) Example of scene for which most participants reported no deviations from the premixture sounds. I) Example of scene with an absence deviation from the premixture sounds.

We found that most of the model’s inferred sources were recognizable in the mixture, with performance in the model condition far better than chance, and only slightly worse than the recorded condition (Figure 9B; recorded 95%CI=[0.82, 0.87], model 95%CI=[0.77, 0.82]). For each inferred source, we computed the proportion of participants that correctly recognized the source as present in the mixture. A histogram of this proportion is skewed left (Figure 9C), indicating that unrecognizability errors were limited to a modest number of inferred sources. Examples of unrecognizable and highly recognizable source inferences are shown in Figure 9D, with additional recognizable examples in Figure 10 (also see “Qualitative investigation” section below). These results indicate that the model often successfully infers sources from everyday sound mixtures that are recognizable to humans.

**Figure 10.**
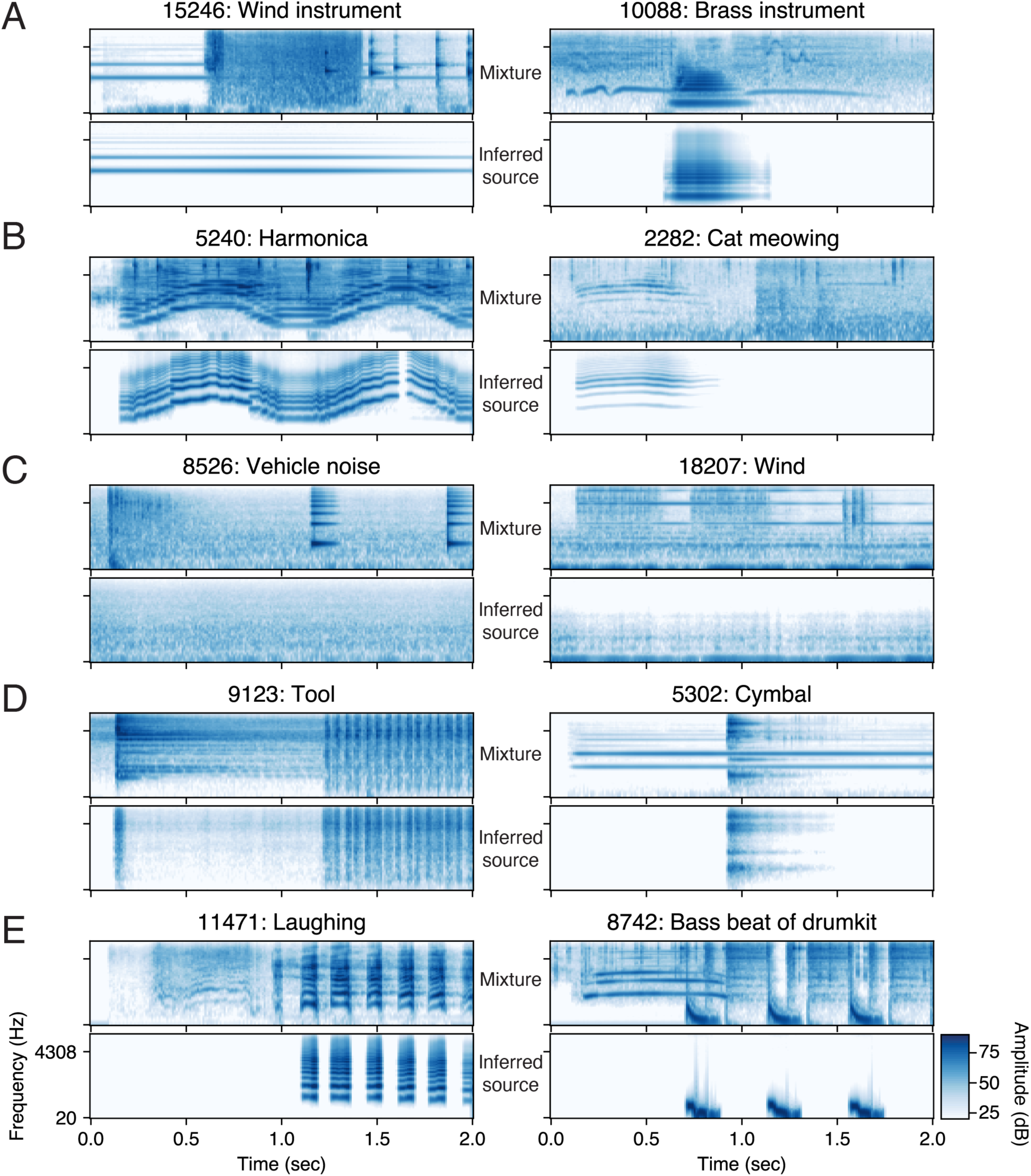
Examples of recognizable source inferences in Experiment 1. A) Harmonic sources, with static fundamental frequencies. B) Harmonic sources with dynamic fundamental frequencies. C) Background noises with relatively static amplitude. D) Broadband amplitude-modulated noises. E) Sound sequences. Listen to the audio examples at https://mcdermottlab.mit.edu/mcusi/bass/. Title numbers refer to FUSS filename and are provided to aid with website navigation.

### Experiment 2: absence, over-segmentation, and over-combination deviations

The purpose of Experiment 2 was to identify when inferences in the model deviated from human perceptual organization via absence, over-segmentation and over-combination of sound sources. We used a matching task between the inferred sources and the recorded premixture sounds, depicted in Figure 9E. Although premixture sounds are not guaranteed to predict human perceptual organization of the sound mixture, they typically formed a coherent perceptual entity in the mixture, and provided a convenient proxy for human perceptual organization.

On each trial, participants listened to inferred sources and premixture sounds from the same sound mixture, which formed the axes of a response grid. Participants were instructed to place a checkmark whenever part of a “row sound” (inferred source) matched part of a “column sound” (premixture sound). We excluded both the premixture sounds and model inferences which listeners could not reliably recognize as part of the mixture in Experiment 1, and therefore required that participants place at least one checkmark for each inferred source. The response grid otherwise contained all the premixture sounds and all the inferred sources for a given mixture.

This setup allowed us to tally the different ways in which inferred sources could deviate in perceptual organization from the premixture sounds (Figure 9F). If a column did not have any checkmarks, this was tallied as an absence deviation for the corresponding premixture sound. If a column had more than one checkmark, this was tallied as an over-segmentation deviation for the corresponding premixture sound. If a row had more than one checkmark, this was tallied as an over-combination deviation for each checked premixture sound. Finally, if a column had only one checkmark and the row that was checked was not checked for any other sounds, no deviations were tallied for that premixture sound.

Listeners’ judgments were reliable despite the binary judgments, with inter-rater reliability of 0.63 (inter-class correlation, *ICC*(2,1) = 0.63, 95% CI=[0.60, 0.67]). The average results across participants are shown in Figure 9G. To obtain chance performance levels for each type of response, we randomly permuted each participant’s responses subject to the task constraint that there be one checkmark in each row, then averaged over participants. This constrained permutation preserves the number of extra checkmarks in rows, such that the level of over combination deviations is preserved (we thus do not have an estimate of the chance level of these deviations).

If the model never explained everyday sound mixtures in a way that corresponded to the premixture sounds, the proportion of premixture sounds with no deviations would be close to zero (below chance). By contrast, if the model inferences tended to replicate the perceptual organization implied by the premixture recordings, the proportion of premixture sounds with no deviations would be well above chance while the proportion of premixture sounds with deviations would be below chance. This latter prediction is what we observed. First, the mean proportion of premixture sounds that had a one-to-one correspondence with an inferred source was greater than would be expected by chance (Figure 9G: ‘no deviations’, p<0.001 by permutation test). Second, the mean proportions of premixture sounds with an absence or an over-segmentation were significantly less than chance (Figure 9G, p<0.001 in each case, by permutation test). And third, the mean proportion of premixture recordings with an over-combination was small (Figure 9G, 95%CI=[0.11, 0.19] by bootstrap). Figure 9H-I depict scene inferences with premixture sounds in these various response categories.

To ensure that the results were not due to participants tending to randomly choose a single inferred source for each premixture sound, we checked whether participants responded consistently. We first analyzed the premixture sounds with no deviations. We determined the most commonly chosen inferred source for each premixture sound (among participants who answered one-to-one for that premixture sound). We then computed the mean proportion of premixture sounds for which a participant chose only the most commonly chosen inferred source. We found that this proportion was much greater than would be expected by chance (Figure 9G: ‘no deviations+most common’, p<0.001 by permutation test), indicating that most of the premixture sounds with no deviations were consistently matched to a specific inferred source. The incidence of deviations for particular premixture sounds was reliable across splits of participants for each of the three types of deviations as well, providing further evidence that the responses were non random (Supplementary Figure 4; p<0.01 by permutation test for all deviation types).

Overall, these results indicate that the model often (but not always) inferred perceptual organization that mimicked the premixture sounds. Supplementary Figure 5 shows the average number of each deviation for each premixture sound.

### Qualitative investigation

To better understand the model’s successes and failures, we also qualitatively examined participants’ judgments for individual sound mixtures, linking them to acoustic structure and sound category labels. Many failures point to generative principles underlying everyday sounds that are missing from our model. A priori it was not obvious that these principles would need to be included in order to account for perception; each could form a productive direction for future research. We note that the ability to observe these failures in the interpretation of recorded audio represents an advance over previous Bayesian models of perception, which typically have not been applicable to actual sensory signals, and so could not be evaluated on arbitrary real-world stimuli (34–38). The combination of stimulus-computability with the interpretability of the model makes the model “richly falsifiable”, with failure modes that provide insight.

#### Hierarchical sources

Sources in our model are constrained to emit events with similar properties, but this assumption is violated in some everyday sounds. We observed over-segmentation deviations when a common source in the world had more than one way to produce sounds (Supplementary Figure 6), such as those produced when writing on a chalkboard (the impacts of the chalk on the board produce brief impulsive sounds, whereas the scraping of the chalk along the board produces more extended and noise-like sounds). Related over-segmentation deviations occur when a real-world sound source concurrently mixed different sound types (Supplementary Figure 7), such as breath noise concurrent with the pitched sound of a flute note. These deviations suggest that human perceptual organization has a type of hierarchical structure missing from our model, whereby humans can hear distinct sound-generating processes as a composite source.

#### Diversity of everyday sound spectra

Other failures result from limitations in the generative model of spectrum and amplitude (Supplementary Figures 8–11). For instance, we found that the model over-segmented frequency components of impact sounds that decay at different rates (Supplementary Figure 8), because it cannot explain frequency-dependent decay as being due to a single source, despite the frequent occurrence of this type of sound in the environment (96, 97). The model also sometimes over combined perceptually distinct sounds. For instance, in the example in Supplementary Figure 9, clarinet and violin sounds are combined into one source, because the model cannot represent the spectral differences that distinguish the two types of sounds. These results point to the richness of the human perception of source spectra and their variation over time.

#### Cochleagram input

One frequent feature of unrecognizable inferred sources was a mismatch in the periodicity of the inferred source and the corresponding everyday sound. For instance, the model explained the sound of rain with a harmonic source (Supplementary Figure 12). This failure is plausibly due to the cochleagram used for the likelihood representation – this representation was chosen for computational efficiency, but its low spectral and temporal resolution limits the resolution with which periodicity can be measured. This low-resolution input representation likely also contributes to absence deviations, for example with quiet tones in noise (Supplementary Figure 13).

## Discussion

Inspired by everyday sounds, we built a probabilistic generative model that describes sources which emit events to produce auditory scenes. We sought to test whether such a generative model could account for illusions that illustrate principles of auditory grouping, and whether the same model could explain human perception of real-world sound mixtures. To address the challenges inherent to Bayesian inference of causes from sensory signals, we combined deep learning with a differentiable generative model: a neural network made event proposals, which were sequentially combined into sources and evaluated with stochastic variational inference in the generative model. The model qualitatively replicated human perception across a variety of classic auditory scene analysis illusions. By contrast, contemporary source separation neural networks could not account for human perceptual organization, whether trained on datasets of recorded audio or samples from the generative model, revealing the difficulty of matching perception comprehensively in the absence of a generative model. Selective model “lesions” revealed the importance of a distribution over sources that allows for variation in source properties. Despite the model’s simplicity, experiments with human listeners showed that the model could also explain the perceptual organization of many natural sound mixtures. The model failures were instructive, revealing the need for specific additional generative principles (and the likelihood that such principles are implicit in human perception). Some deviations pointed to perceptual phenomena that have not been extensively studied (e.g. the perception of hierarchical structure in sound sources) and thus identify directions for future work. Our model represents an advance in the scope of Bayesian models of perception, being able to bridge between the perceptual organization of experimental and everyday signals. Although the results do not prove the necessity of generative models in accounting for perception, they illustrate the many benefits of generative models and show one way it is now possible to instantiate and test the overall approach.

## Relation to prior models

The idea that perception can be explained with Bayesian inference has a long history. But the difficulties of specifying a rich world model and performing inference from actual sensory signals has limited application of Bayesian inference to stimulus-computable models of human perception (40, 98). To meet these challenges, we applied modern tools to integrate and extend key elements of previous work into a unified and functional computational system.

### Structured models

Enabled by modern computational tools, we were able to automatically infer scenes from raw audio while using a world model is that substantially more expressive than in prior work. The basic generative structure of our model posits that parametrized sources produce sound via discrete events. Among previous models that explicitly considered discrete events, most treated them as pre-determined symbolic input to be grouped (37, 38, 42). These models typically were intended to explain the sequential grouping of tones, and cannot be applied to raw audio (where the events are not explicit), and thus cannot be evaluated on natural sounds. One previous model used Gaussian processes to model raw audio for scene analysis but relied on inferring a point estimate of the posterior distribution (32). This approximation prevented automatic inference of the underlying sources (which relies on integrating densities of varying dimensionalities) as well as the inference of bistability. Earlier work on computational auditory scene analysis also attempted to compute detailed scene descriptions from raw audio (4, 22). We built on this work by specifying a richer generative model (e.g., source priors, source-specific kernels, non-stationarity between events) and applying modern techniques to enable inference.

### Clustering

Some computational approaches to auditory scene analysis instead frame perceptual organization as clustering “bottom-up” features of audio (99, 100). This approach has often been combined with bottom-up features inspired by neurophysiology (11, 13, 101–103), but it remains difficult to identify features whose clustering is comprehensively predictive of perceptual organization. Our approach shares some abstract similarities in combining a bottom-up processing component with a top-down inference stage, but a key difference is that the bottom up component of our analysis-by-synthesis algorithm preserves uncertainty, outputting proposals for event latent variables that can be refined or rejected during inference. Maintaining uncertainty over the event proposals was critical to robust inference, particularly with everyday sounds, because the event proposal network frequently detected candidate events that were ultimately discarded from the most probable scene description.

### Machine hearing

We compared our model to source separation neural networks from machine hearing. These particular networks were not developed with the intention of replicating human perception, but other types of neural networks have had success in explaining human auditory perception in various other domains (86–89) (though see (104)), and analogous models trained on source separation (i.e., trained to reconstruct a set of premixture sounds from a mixture sound) might be envisioned to account for aspects of human auditory perceptual organization. We found that a suite of such models with varied training datasets, supervision, and architectures did not comprehensively match human perception as well as our model did, even when trained on samples from our generative model. It is possible that source separation provides the wrong task constraints, or that currently available neural networks are not good enough at source separation to sufficiently constrain their representations. But the results underscore the challenge of accounting for human perceptual organization purely with feedforward neural networks, and suggest the benefits of coupling a generative model with neural networks.

In addition to better accounting for perception in the domain examined here, generative models confer several desirable traits as compared to pure neural network systems. First, our model explains human perception with a set of interpretable principles. As shown in the everyday sounds experiments, this interpretability helps to identify what is missing in the model and how to address its shortcomings. Since the model is composed of meaningful parts, we can augment it by adding more meaningful parts. For example, to create a composite source that emits both noises and harmonics, we could combine the noise and harmonic renderer and define how their latent variables co-vary. It would also be possible to add entirely new generative modules if needed, such as reverberation (with an additional stage of filtering applied to the source sounds). Second, generative models provide a language for specifying structured scene descriptions. Without an explicit generative model to create input data labelled with latent variables, human annotators are required to create data needed for supervised training, which can be inefficient. Moreover, the labels that are convenient for humans to annotate may be limited, e.g. to verbal class labels. In contrast, our generative model uses multiple source-level variables along with a sequence of events that the sources generate, which each include their own generative parameters (e.g. temporal trajectories). The source-level latent variables alone (e.g., variance on fundamental frequency; Figure 3) would likely be difficult for a human annotator to label. In addition, as demonstrated with model lesions (Figure 8), we can manipulate the probability distribution over scenes to help understand the assumptions that underlie the model’s inferences. Such manipulations of training data can be paired with neural network models of perception, but are facilitated by a model to generate the data (88).

## Limitations

### Model structure

The generative model is richer than previous models, and sufficed to account for many classic auditory illusions, but is nonetheless limited by a relatively simplistic model of spectrum and amplitude. The model lacks event-linked structure in amplitude (e.g., with amplitude decay after an impact (97)), time-varying spectra (e.g. frequency-dependent decay (96)), and the tendency for sounds to contain repeated motifs. These limitations were apparent in the everyday sound experiments, and also prevented us from modeling some classical results, notably the asymmetry between onset and offset asynchrony (12, 77) and the perceptual segregation caused by repetition (105). Although these are limitations of the model, the fact that the model can be tested on arbitrary sound stimuli exposes the importance of the missing generative structure for perception.

### Inference

Inference is the central challenge to making generative approaches to perception work. Many of the model limitations reflect choices made to facilitate inference. Moreover, we cycled through multiple iterations of models and inference procedures in attempt to resolve various inference difficulties encountered at different stages of this project. Yet despite these efforts, inference was still not completely robust. The everyday sound experiments revealed three classes of inference failures. First, variational inference sometimes could not recover when the event proposals from the neural network were far from correct, for example proposing the wrong sound type or missing a proposal for a whistle in noise (Supplementary Figure 12–13). Second, gradient descent could get caught in local minima, for example, making it difficult to adjust the fundamental frequency of a harmonic tone or continue a quiet sound behind a masker. Third, the set of hypotheses maintained during sequential inference can become very similar to one another over time, a problem known as ‘degeneracy’. Discarding alternative explanations too early in the timecourse of a sound may set up inference to fail later, when subsequent evidence would favor those explanations.

In addition to not working perfectly, our inference procedure required substantial amounts of computation, and it remains unclear whether this amount of computation is realistic for a biological system (see Methods). We thus do not consider our inference algorithm to provide a mechanistic or algorithmic-level explanation of perceptual organization. Rather, it is a means to implement the computational-level explanation provided by our model (50). Inference is nonetheless a bottleneck for this type of modeling, and an important open scientific issue in perception and neuroscience. The combined efficiency and efficacy of perception demands explanation from both an algorithmic and neuroscientific perspective (see Future Directions).

### Likelihood

A major limitation of our model is the cochleagram likelihood representation. We found that the model quantitatively deviated from human results for illusions that plausibly depend on either a high-resolution representation of frequency (e.g. harmonic mistuning) or time (co-modulation masking release), which likely reflects the limited resolution of the cochleagram. This deficiency was especially apparent for periodic sounds, which also limited the model’s performance with many everyday sounds. Some previous work supplemented a cochleagram with an explicit periodicity representation for this reason (e.g., (13, 22, 101)). We deviated from this previous work for reasons of efficiency (speed and memory for iterative computation). Some alternative representations that might better capture periodicity include the correlogram (106), sparse periodicity acoustic features (107), wavelet scattering transforms (108), wefts (109) or multi-scale cochleagrams (110). Another possibility would be to use representations learned by a neural network to extract periodicity from a high-fidelity simulated cochlear representation (87). But the challenge of defining appropriate mid-level representations for the likelihood extends beyond periodicity. For example, the cochleagram also seems poorly suited for generative inference on sound textures (52), the details of which are often inaccessible to human listeners (111) but which affect cochleagram-based likelihood. We view this as an important open theoretical issue in perceptual science.

## Future directions

One of the contributions of this work lies in providing a model that both has interpretable structure and can be applied to actual audio, and thus richly falsified. The failures are as instructive as the successes, and identify a number of opportunities for modeling advances as well as empirical characterization of perception.

### Hierarchical organization

Our model results on everyday sounds expose the importance of additional levels of hierarchical structure in human perceptual organization. Consider a rhythm played by the bass, snare and cymbal of a drumkit, or a breathy flute note with simultaneous periodic and aperiodic components. The components within such sounds can be simultaneously perceived as distinct and linked. These scenarios are akin to visual hierarchical grouping (34, 112, 113). Sound ontologies and synthesis methods have advocated for multi-layered scene descriptions to describe such scenarios (110, 114–116) and one perceptual model has posited hierarchical structure in tone sequences (37), but we still know relatively little about the perception of such hierarchical structure, especially in naturalistic sounds.

Our model also exposes the question of whether perceptual organization in these cases is based on physical/acoustic causal interpretations. For example, our model produces an over segmentation for the sounds of a ball that bounces on the floor and then strikes a metallic surface (Supplementary Figure 7E). The acoustic structure of the two impact events is dissimilar, but humans hear an integrated causal sequence. One way to account for such examples is with “schema-based” auditory scene analysis in which the auditory system learns to group patterns of sound which recur in the environment (117, 118), as is thought to aid the streaming of music and speech (119, 120). But it is also possible that the auditory system constructs more specific causal models, for instance representing sounds from physical events using internal models of physics (115, 121, 122), as is thought to occur in vision (30, 123, 124). Physics-based sound synthesis should enable additional work in this direction (97, 125, 126).

### Sound textures

Another notable model shortcoming occurred for sound textures (52) (Supplementary Figure 10– 11). In one example, the model explained a mixture of temporally overlapping textures (running water and brushing teeth) as a single noise source, whereas humans tend to hear two separate streams (Supplementary Figure 10A). The model is limited both by a noise model that does not capture all of the statistical regularities that differentiate natural textures (52), and by a likelihood function that is applied to the cochleagram rather than to a statistical representation akin to that thought to determine human texture perception (111). But this limitation also exposes the need to better understand the role of texture in auditory scene analysis. The auditory system is known to ‘fill-in’ textures when they are masked (127), indicating that multiple textures can be heard at the same time, and to selectively average sound elements attributed to a texture (128). But we know little about the conditions in which humans segregate texture mixtures, pointing to another direction for future work.

### Alternative scene descriptions

The limitations imposed by the model’s scene descriptions raise the question of what an ideal model’s descriptions should contain, which in turn raises open questions about the content of human perceptual experience. Our model’s scene descriptions were relatively abstract and signal-based, with a division into sound types that has historically been common in sound synthesis (22, 51, 129, 130). These could be enriched and extended in many ways. One question is whether the source models of human perception are closer to a flat hierarchy of many source models that each describe a fairly specific set of sounds (impacts, speech, woodwind instruments, textures, modern electronic sounds, etc.), as opposed to a deep, unified hierarchy in which a modest number of primitives are composed to create more complex sounds. In addition to more descriptive source models, an ideal model of scenes should include other aspects of sound generation, such as reverberation and spatial effects (96, 131–134).

### Learning

An alternative to designing source models is to learn them from sound data. One possibility is to combine learned statistical components into a structured generative model. For example, one could swap out our classic Bayesian priors (Gaussian processes) over excitation and spectra for autoregressive neural networks (135) trained to synthesize natural sounds through the model’s renderer. The discrete sound types could also be swapped out for a more flexible learned representation. Integrating learned components into a structured model would allow the model to capture a richer class of sounds accurately while still maintaining interpretability (136).

### Inference

If Bayesian models are to be taken as mechanistic explanations of human perceptual systems, then its key algorithmic formalism – search through a hypothesis space – must be shown to be plausible. We found that everyday sound mixtures and experimental stimuli provided complementary tests of our search procedure, in part because they tended to have distinct inferential demands. Typically, psychophysical experiments are designed to be ambiguous with respect to just a few reasonable perceptual hypotheses, in order to isolate the effect of a single variable. In experiments, and in most classic illusions, listeners are instructed to report one of these hypotheses, and are given practice to help them get accustomed to doing so. Because the hypotheses are delimited to a small set, the inferential difficulty lies in accurately estimating the posterior distribution: it may be difficult to estimate the relative probability mass between the two hypotheses because they can differ in subtle ways (e.g., the presence of a quiet tone, or the placement of a formant). In contrast, when inferring explanations for everyday sound mixtures, there are many more plausible hypotheses in a model rich enough to explain them. It appears as if the local information is less diagnostic in such settings, but the overall global scene interpretation is less ambiguous (as noted for visual inference (46)). This means that adjudicating between hypotheses is easier for everyday sound mixtures than for typical experimental stimuli, but that search is much harder. We thus think that testing models on everyday sounds is critical for future progress, as it exposes challenges that are less apparent in classical experimental settings.

We believe that solving the challenge of narrowing search while maintaining multiple hypotheses is central for a mechanistic account of perceptual organization. For our model, using an amortized inference network to propose events was necessary for tractability – the network narrows the search space to a small subset of all possible events. But because these event proposals were combined into scene hypotheses, assessing all of their combinations was intractable when there were many event proposals (as was typically the case for everyday sound mixtures). Search relied on simple heuristics to prioritize smaller scene descriptions, as well as substantial parallel computing resources. If perception is solving a similar search problem in real-time, it seems likely to utilize more efficient procedures to search this combinatorially large space. We suggest that more plausible algorithmic accounts could be discovered by learning procedural knowledge. For instance, over many inference trials, we could observe which combinations of events proposals are successful versus unsuccessful. These could then be used to learn how to prioritize the assessment of certain combinations (137, 138), in place of the simple hand-designed heuristics used here. Other aspects of the search algorithm could also be replaced by learnable components (e.g. the stochastic gradient descent used for hypothesis optimization (139)). An ambitious future direction would be to replace the entire search algorithm with a fully learned procedure, as has been done for simple compositional graphics programs (140).

### Time

Our model was designed to perform joint inference over the sources and events that produced an entire observed sound (2 seconds in our everyday sound experiments). However, human perceptual judgements are sensitive to context within a local window, and do not integrate evidence from an arbitrarily long time-horizon (128, 141). Such integration windows could be incorporated by explicitly including memory representations. One possibility is to modify sequential inference such that variables eventually stop being actively inferred and become fixed as a memory trace, no longer affected by further observations but potentially informing future variables. Such an approach could enable investigation of the timescales for which inference improves in complex scenes, which could clarify whether the temporal extent of memory is adapted to the scale of temporal dependencies in natural scenes, or whether it mainly reflect resource constraints.

### Attention

Another aspect of auditory experience that is missing from our current model is the ability to attend to a particular sound source. Attention can be construed as a mechanism that selectively refines parts of a perceptual hypothesis in order to deal with the intractability of inference in generative models (142), but the difficulty of inference in such models has prevented such ideas from being tested. Our model offers an opportunity to explore such computational accounts of attention. One possibility could be to vary the dimensionality of the approximate variational posterior over parts of a scene hypothesis as a function of attention. For example, increasing the time-resolution in the variational posterior over the excitation trajectories would enable inference of finer-grained detail in those trajectories. Along these lines, attention could be instantiated in a model like ours by increasing/decreasing the complexity of the approximate variational posterior distribution for attended/unattended sources, respectively. Implementing attention in this way could help explain psychophysical benefits of attention (143, 144) in a normative framework, and provide a formal hypothesis for the targets of object-based attention (145, 146).

### Illusion generation

A generative model of perception enables both analysis and synthesis of sound. One intriguing application is to use the model to generate illusions by inferring the stimulus properties that would produce a desired percept (147). One could formalize common illusion paradigms, such as multistability or conflicting global and local percepts, into objectives for optimization, providing a powerful additional model test.

### Other sensory modalities

Many of the principles implemented in this paper can be traced to classical ideas in vision, where inference in generative models has long been proposed to underlie perception, but where computational systems that embody this approach have historically been challenging to make work. Our results demonstrate the utility of auditory perception as a case study in perceptual inference. Audition has the advantage that relatively simple generative models can account for many everyday signals, making the approach tractable. However, the general principles and questions explored here are not modality-specific, and the time seems ripe to reapply this framework to other perceptual domains, and to multi-sensory perception.

## Methods

### Generative model: Overview

We begin by specifying the overall probabilistic structure of the model (Figure 2B). A full scene description *S* is composed of a number of sources *n*. Each source description *s_i_* has a sound type *T_i_*, a set of source parameters *Θ_i_*, and a number of events *m_i_*. A source generates a set of *m* event descriptions, *E_i_*, from its event priors defined by *Θ_i_*.

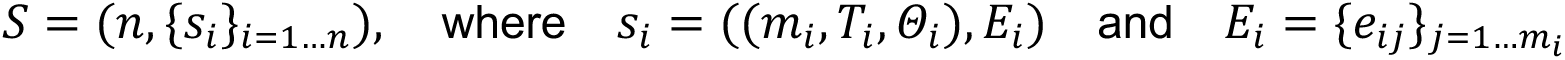

The sound type determines the structure of the source (e.g., whether it has a fundamental frequency) as well as how its sound is synthesized. For each source, the event descriptions are input to a renderer that synthesizes a source sound (to be described more in the Likelihood section below). The source sounds are summed to create a mixture.

The sources are independent, meaning 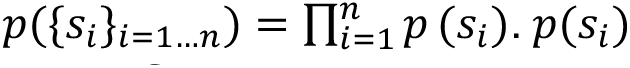 further factorizes, yielding the following hierarchical model for the scene *S*:

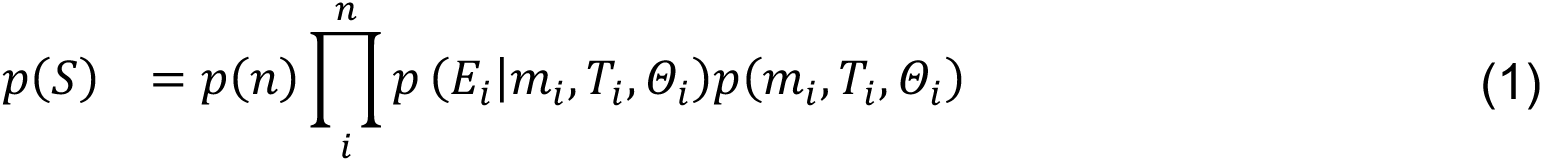

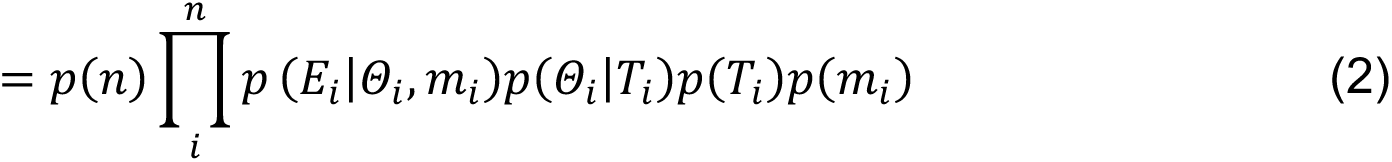

The hierarchical factorization into event priors *p*(*E_i_*|*Θ_i_*, *m*_)_) and source priors *p*(*Θ_i_*|*T_i_*) allows different sources to have different tendencies when emitting events (Figure 3A-B). The next section will cover sources and events, followed by a section covering source priors.

For the sampling procedure that defines the entire generative model, see Algorithm 1 in the Supplementary Information. This algorithm reflects the actual probabilistic program we use, which uses *for* statements to express hypotheses that vary in dimensionality (e.g. a variable number of sources, *n*) and *if..else* statements to express how categorical random variables can change the structure of the hypothesis (e.g., sound type determines whether events have a fundamental frequency).

## Generative model: Sources and events

A source emits events, which are input into a synthesizer to produce a sound waveform (Figure 2B). From the perspective of sound synthesis, the events define the *excitation* that provides sound energy, which in the case of harmonic or noise sources, is additionally passed through the source’s *filter* to generate sound. The filter is fixed across events emitted by a given source, on the grounds that this is common in sound generation (for instance, where a filter corresponds to the instrument body). The excitation can change in time, turning on or off abruptly as well as changing continuously (e.g., in amplitude). From a probabilistic perspective, the source parameters *Θ_i_* parametrize a prior distribution over events, *p*(*E_i_*|*Θ_i_*, *m*_)_) (Figure 3A-B). First, we will define the random variables that make up the event, specifically the excitation. We will return to the filter (which is shared across events) at the end of this section.

As depicted in Figure 2B, each event *e_ij_*. consists of its onset and offset **τ***_ij_*., as well as time-varying excitation trajectories (amplitude *a_ij_*.(*t*) and, if periodic, fundamental frequency **f***_ij_*.(*t*)) which depend on the onset and offset of each event (**τ***_i_*).

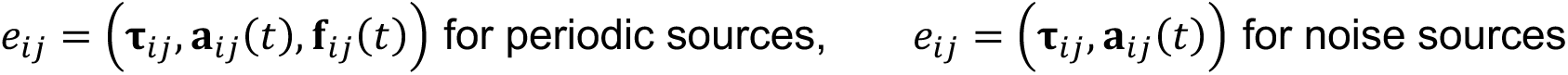

Events within a source are not independent from each other. Rather, events are sampled sequentially, such that each event depends on the events before it. For example, the full event prior for a periodic source *s_i_* is:

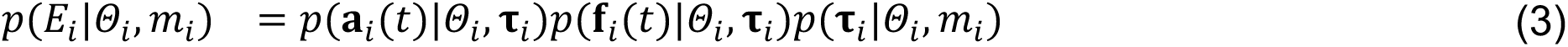

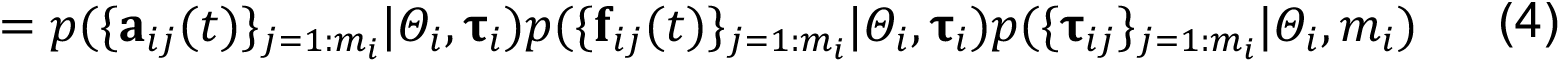

with individual events conditioned on the preceding events (the first event has an onset sampled uniformly from the duration of the scene, and trajectories sampled from an initial Gaussian process described below):

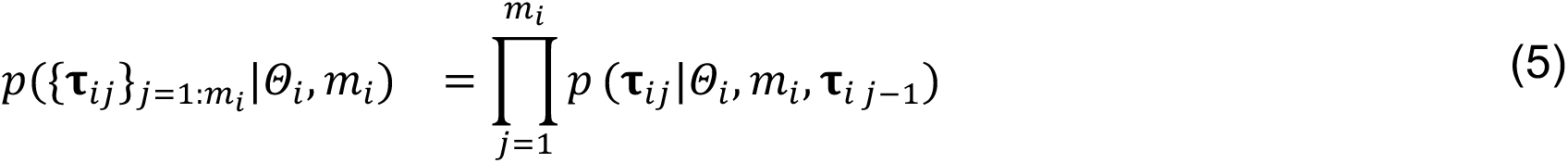

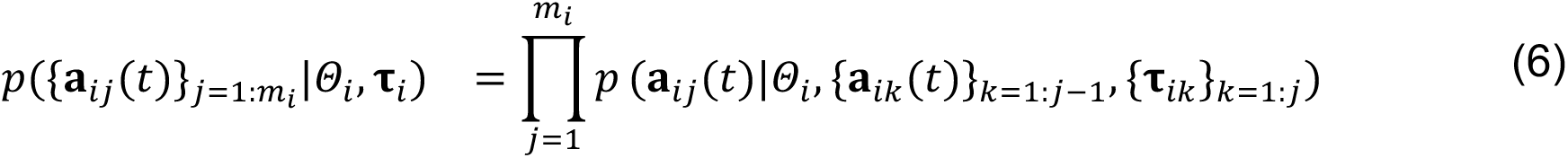

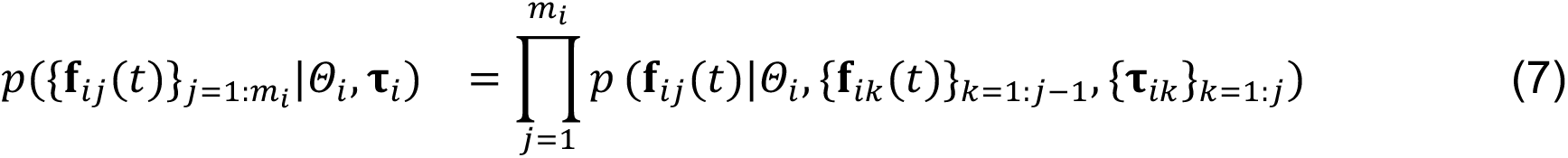

Note that the onset and offset of an event (**τ***_ij_*.) are only dependent on when the previous event occurred (**τ***_ij_*_.-1_), but the trajectories of event *j* (*a_ij_*, **f***_ij_*) are dependent on its onset/offset (**τ***_ij_*.) because the onset and offset specify where the trajectories are sampled (i.e., between the onsets and offsets of events).

We now describe the event onset and offset and how they are sampled from sources. To constrain the events to be non-overlapping in time, we reparametrize an event’s onset 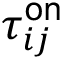 and offset 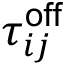 as the silent “rest” interval 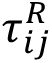 preceding *e_ij_*. and its active interval 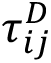. If the durations 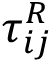 and 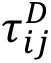 are non-negative then the events will not overlap. The event priors on these variables are Log Normal, and we sample the event rest and duration as:

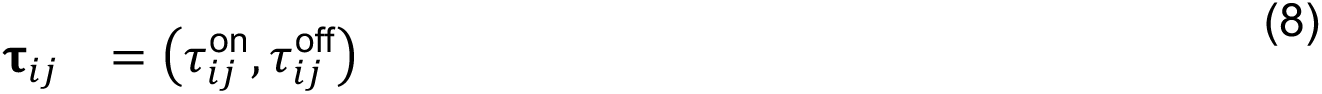

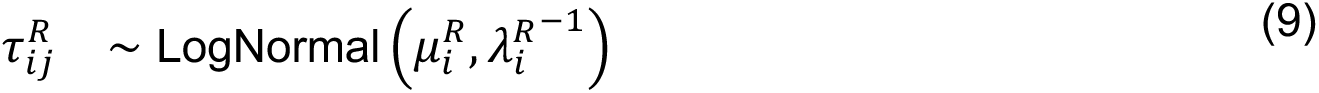

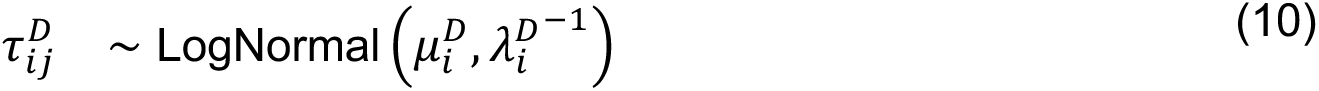

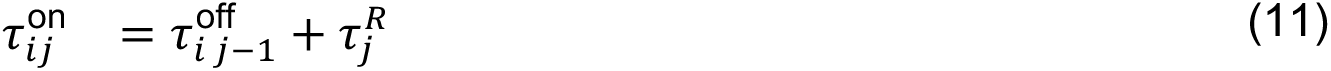

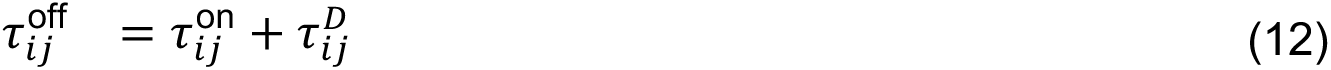

The only exception is the first onset which is sampled uniformly from the duration of the scene.

The sampled event timings will reflect source regularities as captured by the parameters of the log-normal event priors. These parameters are source parameters: 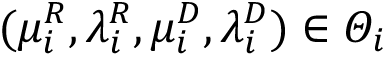 (Figure 2B). The source prior of each (*μ_i_*, *λ_i_*) pair is a normal-gamma conjugate prior which is shared across all sound types (see source priors below; Methods Table 1). These source priors allow the model to account for a wide range of temporal regularities, as occur in natural sounds (Figure 3A; note λ=σ^-2^): for example, a source could tend to have long events (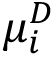 is high), events occurring in rapid succession (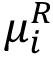 is low), or a wide variety of event durations (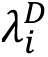 is low).

In addition to the onset and offset (**τ***_ij_*), events have excitation trajectories. The event excitation amplitude **a**(**t**) and frequency **f**((**t**)) trajectories are vector variables that represent a time-varying function, and thus require an event prior over functions. Therefore, we sample the time-varying excitation trajectories from one-dimensional Gaussian Process (GP) priors (148). GPs are distributions over functions *g*(*x*), for which any finite set of function values *g*(*x*_+_), …, *g*(*x*_-_)} yield a multivariate normal distribution. GPs are thus characterized by their mean function *μ*(*x*) and a kernel function *κ*(*x*, *x*′) that specifies the prior covariance between *g*(*x*) and *g*(*x*′). Given thesefunctions and a vector of timepoints (**t**) which lie within an event (i.e., 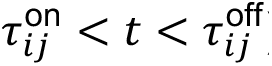), we can sample the trajectory of the time-varying excitation variables **a**(**t**) and **f**((**t**)). To instantiate source regularities for the excitation trajectories, we induce temporal correlations both within and between events in the kernel function (see below).

We used mean functions that were constant, reflecting the tendency of sources to be high or low in amplitude or fundamental frequency. The entire time-varying excitation trajectory is sampled from a GP with a constant mean function (e.g., for amplitude 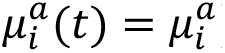) and non-stationarykernel function *k_i_*(*τ_i_*); we call this *g*_1:*m*_. Taking the excitation amplitude *a_i_*((**t**)) across all events of source *i* as an example:

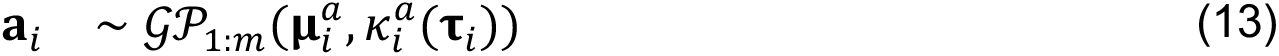

Since inference can proceed sequentially, we write the following to indicate the same overall distribution for convenience in notation (as used in Algorithm 1):

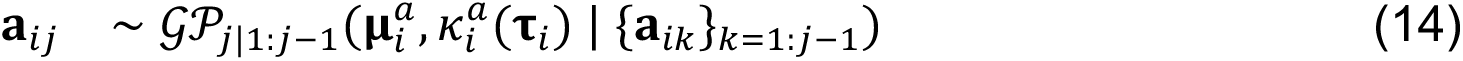

Given that the Gaussian Processes are event priors, their parameters (specifying the mean and kernel functions) are source parameters that define the regularities in the source’s events (Figure 3C). The mean constant *μ_i_* determines the central tendency of the trajectory (e.g. quiet vs. loud). The kernel *k_i_* determines the shape of the trajectories which are likely under the GP prior.

The first term of *κ* is a standard squared exponential kernel (SE) that instantiates a prior favoring smoothly-varying excitation trajectory both within and across events, which is characteristic of many natural sounds (149). The second term is a non-stationary kernel (NS) which specifies higher covariance for trajectory values that occur within the same event, compared to across events. This non-stationary kernel can express a single source that slightly modifies its sound generating process between events. In natural sounds, this occurs in a variety of ways: for instance, a dog panting is quieter on the in-breath than the out-breath, and a flute can discretely switch pitches between notes (Supplementary Figure 1). Therefore, we implement a non stationary kernel for all excitation trajectories, as follows:

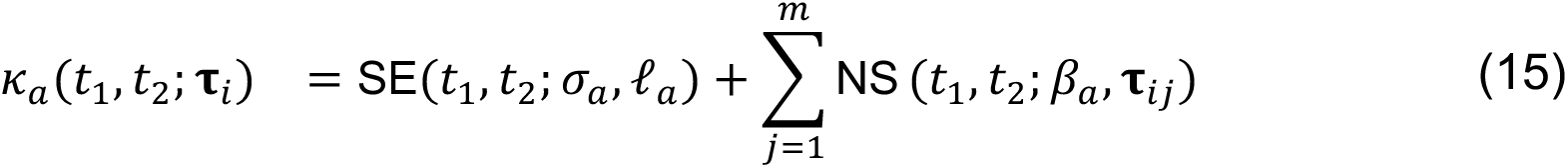

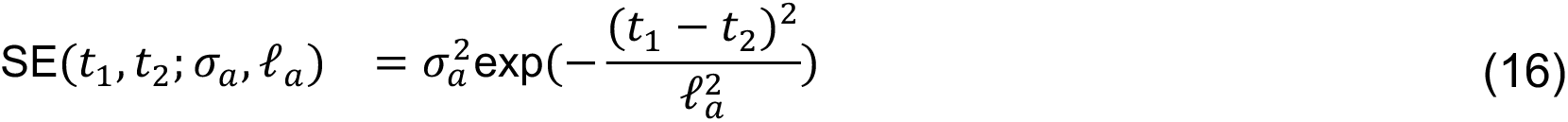

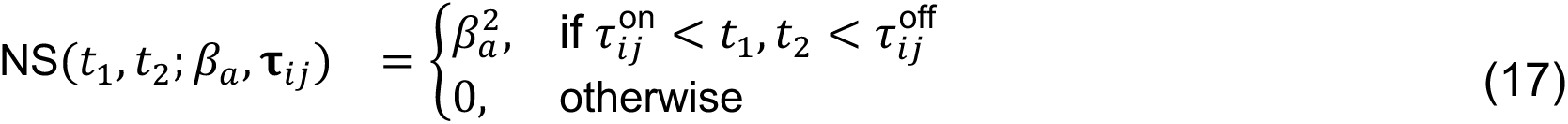

We turn now to the filter, which is defined analogously to the excitation trajectories. The filter shape, which is a function of frequency (Figure 2B), has a one-dimensional stationary GP prior. Like the excitation trajectories, the prior over the spectra of harmonic sources uses an SE kernel, exhibiting a tendency to vary smoothly. Because of the prevalence of bandpass noise with sharp cutoffs, we used an Ornstein-Uhlenbeck (OU) kernel for noise source spectra. Like the squared exponential kernel, the Ornstein-Uhlenbeck kernel exhibits a tendency to revert to the mean, but differs in that it does not tend to be smooth. For all sound types, the sampled filter **H** is shared across all events emitted by a source.

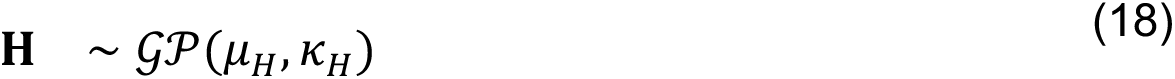

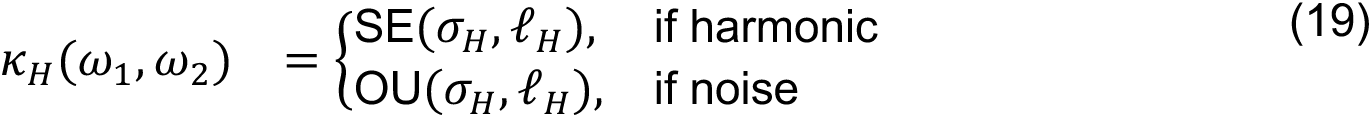

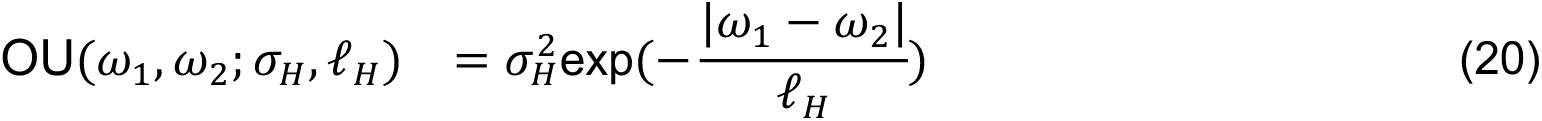

The sampled excitation trajectories and filters reflect source regularities that are defined by the parameters of the GPs. The mean *μ* as well as the kernel variables *σ* and ℓ of each GP are source parameters (see source priors below and Methods Table 2). The means are sampled from an appropriately scaled uniform distribution. The inverse softplus of *σ* and ℓ are distributed normally (to maintain positivity and numerical stability; a Log Normal distribution would provide a similar functionality but was numerically unstable during gradient descent for variational inference). These source priors allow the model to account for a wide range of regularities in the excitation trajectories and the filters, as observed in natural sounds. For example, a source could have a flat spectrum (low *σ_H_*) or a spectrum with widely spaced peaks (high *σ_H_* and high ℓ*_H_*).

## Generative model: Source priors

In the preceding section, we described how event distributions are parametrized by source parameters, and how different values of the source parameters allow the model to account for different regularities in sounds (Figure 3). These source parameters are themselves sampled from distributions called source priors. For clarity, we use the term “meta-source parameters” for the parameters that define the source priors. The meta-source parameters that define the source priors are listed in Methods Tables 1 and 2.

### Temporal source priors

The duration and rest source parameters, 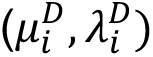 and 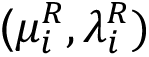 espectively, are sampled from normal-gamma source priors. The meta-source parameters of these source priors were chosen to be only weakly informative, covering a large range (see Methods Table 1). We restricted the rest and duration source priors to be the same as each other, and the same across all sound types. In practice, we found the specific settings of these parameters had little effect on the results.

### Gaussian process source priors

Each Gaussian process has source parameters defining its mean and kernel. Since the Gaussian process kernel source parameters encode relatively complex constraints, we fit their source priors to natural sounds. We first describe the natural sound datasets used to fit the source priors. Then, we describe the specific form of the source priors and the procedure for fitting the source priors to the natural sound datasets.

We chose publicly available datasets of recorded sounds for which the dominant sound type was obvious and for which annotations of the secondary, event-level variables were available (onset, offset and fundamental frequency where applicable). We used these annotations to facilitate stable inference of the meta-source parameters (described more below). For noises, we used a subset of the sound texture dataset used by (52) which were obviously aperiodic. For harmonic and whistle sources, we used three different datasets that covered a variety of sounds: speech, music, and bioacoustics. For speech, we used the Pitch Tracking Database from Graz University of Technology (PTDB-TUG), which provided recorded audio of speakers saying English sentences and pitch trajectories extracted from corresponding laryngograph signals (150). For music, we used the University of Rochester Multi-Modal Music Performance (URMP) dataset, which contained recorded audio of various instruments playing classical music along with corresponding MIDI and pitch tracks (151). For bioacoustics, we used the Synth Birds Database (SynthBirdsDB) which contained recorded audio from a variety of bird species, with either harmonic or pure tone vocalizations (152). This dataset was compiled to develop pitch-tracking for bird vocalizations, so we used a subset of the data for which accurate pitch tracks were provided.

Each source prior was fit to one or more of the datasets, with the goal of maximizing the variability in the set of sounds for each source prior, while ensuring that the model was appropriate for the included sounds. The amplitude and spectrum source priors for the noise sound type were simultaneously fit to the sound textures dataset. The amplitude and spectrum source priors for the harmonic sound type were simultaneously fit to the music and bird datasets (URMP and SynthBirdsDB). We omitted the speech dataset (PTDB-TUG) here because the time-varying formants of speech could not be well-modeled with the model’s constant spectrum constraint. The amplitude source prior for the whistle sound type was fit to sound clips from SynthBirdsDB hand selected to contain bird vocalizations with a single dominant frequency component, omitting harmonic speech and music. Last, the fundamental frequency source prior was fit on the speech, music, and bird datasets, and was shared between the whistle and harmonic sources. To use all three datasets, we directly fit a GP to the pitch track metadata rather than to the recorded audio.

To fit a given source prior, we simultaneously inferred scene descriptions for the set of natural sounds described above. Each scene description was constrained to use a single source of the specified sound type. We conditioned the model on the annotated secondary event variables to facilitate stable inference of the meta-source parameters (even though the datasets were chosen to feature a dominant source type, a single source was not always sufficient to explain the audio; e.g., due to the presence of background noise). The meta-source parameters (which in the normal use of the model were fixed) were allowed to vary, instantiating a learnable distribution over the corresponding source parameters. The meta-source parameters describing this distribution were inferred by variational inference, along with the source parameters and unconditioned event variables, to maximize the marginal likelihood of the full set of sounds. The inferred meta-source parameters were then fixed as constants for the experiments described in this paper (both those with classic illusions and those with natural sounds).

Specifically, we fit an inverse softplus normal source prior for each of the variance *σ* and lengthscale ℓ kernel parameters of a GP prior. For time-varying excitation trajectories with an additional non-stationary component in the kernel (Methods: Equation 15), we also fit a single value of the non-stationary parameter *β*, shared across all sounds. Due to the computation required for fitting many scenes simultaneously, we randomly selected a subset of two-second clips from the appropriate dataset(s) to fit each set of source priors (approximately one minute of audio in total). This small amount of data was sufficient given that we fit only 2-3 meta-source parameters per source prior.

The source priors over the GP means were uniform distributions. We manually set the bounds of these uniform distributions to cover a natural range of values (fundamental frequency could span the range of frequencies audible to humans and below the Nyquist limit; amplitude was bounded below by the quietest audible amplitude) and to capture the range of means inferred during source prior fitting. The spectrum means were fixed at zero (because the spectrum and amplitude trajectory were summed, the means were redundant, and we removed this redundancy by fixing the spectrum mean to zero and allowing the amplitude mean to vary).

In our model lesion experiments, we found that using uniform source priors, instead of source priors fit to natural sounds, resulted in a negligible difference in the model’s similarity to human perception (see Results: Model lesions and Figure 8A). Although it seems likely that using a model based on real-world sounds is important for capturing human perception, this result suggests that the structural constraints in the model (that were inspired by observations of natural sounds) may be more important than the specific shape of the priors as defined by the underlying meta-source parameters.

## Generative model: Discrete priors

There are three discrete priors: the prior over the number of sources *p*(*n*), the prior over the number of events in a source *p*(*m*_)_), and the prior over the sound type of a source *p*(*T*_)_). Methods Table 1 shows the parameters that define these priors.

To generate scenes with any number of sources and events, *p*(*n*) is a Poisson distribution and *p*(*m_i_*) is a Geometric distribution. Because sources are independent, we use a Poisson distribution *p*(*n*) to specify the probability of some number of sources occurring during a known time interval. Because events in a source are not independent of each other, we use a Geometric distribution *p*(*m_i_*) to describe the probability that *m_i_* events are emitted in total before source *s_i_* ceases. It was critical for the generative model to have the flexibility to allow the number of sources and events to vary but we found the specific parameters of these distributions had little effect on the results.

The sound types are equally likely, with *p*(*T_i_*) as a uniform categorical distribution over the whistle, noise, and harmonic classes. Without the inclusion of a separate whistle class, we would have required a bimodal prior on the filter for periodic sounds (sharply peaked at one frequency versus smooth), essentially amounting to a categorical difference. For simplicity and clarity we opted to instantiate this difference at the level of the sound types.

## Generative model: Likelihood

To compute the likelihood, the sampled scene description *S* must be rendered into the resulting sound *X* (Figure 3D). The scene description was sampled as described in the Priors section above (also see Algorithm 1 in Supplemental Information). The amplitude and frequency trajectories (**a** in dB, **f** in ERB) were sampled at 10 ms intervals for illusion stimuli and at 20 ms for the everyday sounds (to increase memory efficiency; the everyday sounds generally required more memory because they often contained events that extended over seconds). The filter shape was sampled at 0.3 ERB intervals (**H** in dB/Hz for noise sources and dB for harmonic sources). To render the sounds specified by these sampled latent variables, we first synthesized the sound corresponding to each sampled event in each source. We then concatenated the events and silent intervals to construct each source sound. Finally, we summed the source sounds to produce the final auditory scene (Figure 2B, bottom).

To enable variational inference, events of different sound types were differentiably rendered. Each event type was generated by combining an initial excitation with spectral and/or temporal amplitude modulation (filter). For whistle events, we generated a pure tone that was frequency modulated according to **f**. To amplitude-modulate the tone, it was windowed with half-overlapping cosine windows. Each window was scaled by the corresponding amplitude in **a**, followed by overlap-add to synthesize the tone. For noise events, we generated a pink noise sample as excitation. Given the degeneracy of the excitation and filter, the pink spectrum is an arbitrary choice. This noise sample was frozen with a fixed random seed to enable differentiability. The noise was then amplitude-modulated following the same procedure used for tones. Finally, a log spaced, cosine-shaped, half-overlapping filterbank scaled by **H** was multiplicatively applied in the frequency domain. For harmonic events, we generated a set of 200 pure tones at the harmonic frequencies specified by **f**. Any samples of the pure tones that exceeded the Nyquist limit were set to zero to prevent aliasing. We scaled the amplitudes of each tone so that they were pink with respect to the fundamental, and then summed them to produce a complex tone that served as the excitation. The sampled spectrum **H** was shifted in frequency at each timepoint so that its first channel aligned with the fundamental frequency **f**. Then, spectral and amplitude modulations were applied as for noise sources. For all sound types, sigmoid on-ramps and off-ramps were applied to each event (duration=18 ms), with the sampled onsets and offsets corresponding to the maximum of the ramps. Sigmoid ramps were chosen for their differentiability. We rendered all sounds at 20 kHz.

We used a differentiable implementation of gammatone-based spectrograms for our likelihood representation (Figure 3D) (153). This particular version of a cochlear-inspired representation was chosen because of the speed of computation during stochastic variational inference, which requires thousands of iterations. A conventional spectrogram was calculated using FFT, after which the frequencies bins were pooled into channels that approximated the tuning of a gammatone filter bank (Glasberg & Moore, 1990) . We used a window size of 25 ms, hop size of 10 ms, 64 filters with a half-ERB width, and a lower threshold of 20 dB relative to a fixed internal reference level. The sampled scene gammatonegram was compared to the observed gammatonegram under an isotropic Gaussian noise model. The standard deviation on the noise model was fixed to *σ* = 10 across all inferences. The noise standard deviation was manually chosen instead of being directly fit to human data (which would have been computationally prohibitive).

## Inference: overview

We maintain a separation between our model and the inference procedure that infers explanations of audio waveforms under the model: the former encapsulates our scientific hypotheses about perception while the latter is an engineering component necessary to assess whether the model can explain aspects of perception. We believe there is more than one way to engineer a working inference algorithm for this model (e.g., instead of variational inference, it might be possible to utilize fully Monte Carlo based inference, or fully amortize inference). The particular method presented here is not intended as a hypothesis of how perception works, though the challenges of inference that are exposed by our algorithm remain central challenges for complete theories of perception.

We used two modes of inference – sequential and enumerative inference – depending on how an illusion was evaluated in human listeners. Both involve optimizing and comparing hypotheses, but they differ in how they determine the hypotheses to evaluate in the first place. Here we give an overview of these two modes of inference.

Inspired by analysis-by-synthesis, we combined bottom-up and top-down inference processes. In sequential inference, the first two *bottom-up* steps go from the observed sound to a set of hypotheses. The last two *top-down* steps use the generative model to refine and evaluate these hypotheses. Given an observed sound, the steps of sequential inference are as follows (Figure 4):

1. **Events proposal**: A neural network operating on the cochleagram proposes candidate events, which initialize event variables in the hypotheses (described in the Methods section above, Generative model: Sources and events). This “amortized” inference (47, 154) allows us to benefit from existing state-of-the-art machine learning architectures (155) to find good hypotheses. Although the neural network is trained on data produced by the generative model, it may misdetect events or detect multiple alternative event explanations for the same sound, often requiring the generative model to assess the candidate events (in the steps that follow).

Then, to build up the hypotheses sequentially in time beginning with the earliest candidate event, steps 2-4 are iterated until all candidate events have been assessed.

1. **Source construction**: Inspired by sequential Monte Carlo approaches (41) and similar to some previous models of auditory scene analysis (22, 42), candidate events corresponding to the current timestep are combined into scene descriptions through three update actions (add candidate event to existing source, create new source with candidate event, leave out candidate event). This results in a set of hypotheses for the sound up to a particular moment in time. By building up hypotheses sequentially, we avoid searching a combinatorially large number of combinations of events into sources. Nevertheless, there may still be more hypotheses than are efficient to search, so they are prioritized for hypothesis optimization with a set of heuristics.
2. **Hypothesis optimization**: The hypotheses are refined using gradient-based optimization. Specifically, for each hypothesis, a guide distribution is optimized to best approximate a mode of the posterior distribution using variational inference. Variational inference allows us to benefit from a fully differentiable generative model, jointly optimizing all of the continuous latent variables in a hypothesis in order to fit each mode as closely as possible.
3. **Scene selection**: The posterior probabilities (approximated with importance sampling based on the optimized guide distributions) are used to compare the alternative hypotheses. A set of hypotheses with the highest posterior probability are selected for the next round of source construction. The last two steps comprise the top-down component of sequential inference.

This process results in a set of scene descriptions, each with an associated probability. Sequential inference is appropriate for modeling unconstrained listening with illusions and everyday sounds, because it automatically infers the number of sources and events, each source’s sound type and source parameters, as well as the events emitted by each source.

In enumerative inference, which is more appropriate for the constrained setup of psychophysical experiments, we directly assess the probability of competing experimenter-defined hypotheses (using only steps 3 and 4 above). Sequential inference can also be used in psychophysical experiments, and we relied on it in this way for one of the alternative models in Figure 8. This alternative model demonstrated that sequential inference generally resulted in similar overall human-model similarity as enumerative inference (see Results: Model comparisons).

We first explain hypothesis optimization and scene selection, because it is common to both modes of inference (steps 3-4). Then we explain sequential and enumerative inference in turn, in particular, how they determine the hypotheses to evaluate.

### Inference: hypothesis optimization and scene selection

Instantiating perception as inference requires determining the posterior probability of different hypotheses. We use variational inference to obtain an approximation of the posterior distribution. We were able to utilize variational inference because we designed the generative model to be fully differentiable. In this section we first clarify what hypotheses correspond to in our model, and then explain how we compare the posterior probabilities of hypotheses.

We assume that a hypothesis *H* corresponds to a region containing a “mode” of the full posterior. There are two motivations for this assumption. The first reflects the potential uncertainty over the latent variables defining a hypothesis. For instance, a particular hypothesis could specify a single source with one high and one low whistle, but there may be some variance around the exact frequencies of the whistle. A hypothesis is thus characterized by a specific setting of the structural variables (e.g. the number of sources and events), as well as an approximate setting of continuous variables (e.g. source parameters, onset timings). This corresponds to the organization of specific events into sources, while only approximately specifying the low-level features of the events. To compute the posterior probability of this hypothesis requires marginalizing over the continuous variables.

The second reason to associate hypotheses with regions rather than points is to allow comparison of hypotheses that differ in dimensionality. Scenes with potentially different dimensionalities cannot be compared by their posterior densities (which have different units), making point estimates of the posterior mode insufficient. Instead, we need to integrate each hypothesis into a posterior probability mass so that we can compare across dimensionalities.

Formally, each hypothesis *H_i_* corresponds to a region of scene space (*H_i_: S_i_*), and the hypotheses are compared based upon their posterior odds ratio (i.e., relative posterior mass contained within the regions). For two hypotheses and an observed sound *X*, the posterior odds is:

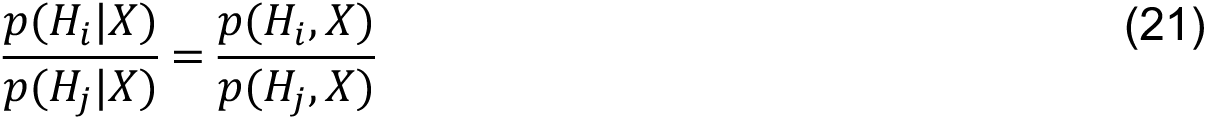

Since *H_i_* contains many specific scenes *S_i_*, computing the marginal probability *p*(*H_i_*, *X*) requires integrating over *S_i_*.

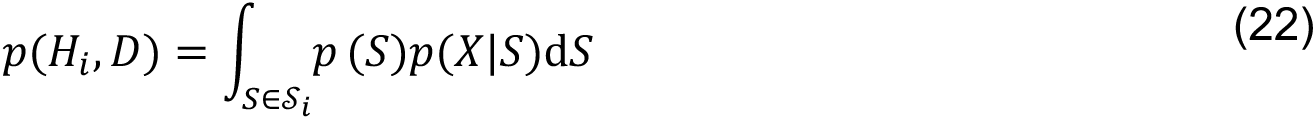

To approximate this integral, we use importance sampling. In importance sampling, we use a guide distribution *q_i_* corresponding to hypothesis *H_i_* to take samples.

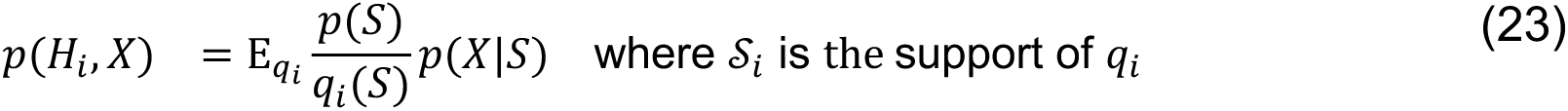

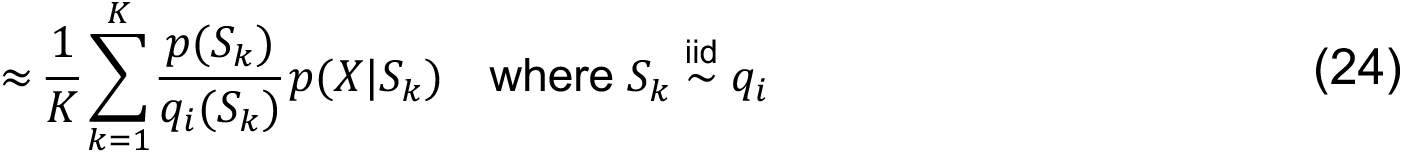

The success of this method is determined by how closely the guide distribution *q_i_* approximates a mode of the posterior. In order to derive a “good” guide distribution, we start with an initial guide distribution which is either 1) based on the amortized neural network output in sequential inference (specifically, the means of the guide distribution were set to the neural network’s output values; see Source Construction section below) or 2) based on experimenter-defined hypotheses in enumerative inference (see Enumerative Inference section below). In both cases, this initial guide distribution is then refined using stochastic variational inference. The form of the guide distribution is mean-field, except for the vector variables (e.g. time-varying amplitude) which have a Gaussian Process prior. For these vector variables, we use a variational inducing point framework (156).

To refine the guide distribution, during each iteration of gradient descent we optimize *q_i_* with respect to the standard variational objective,

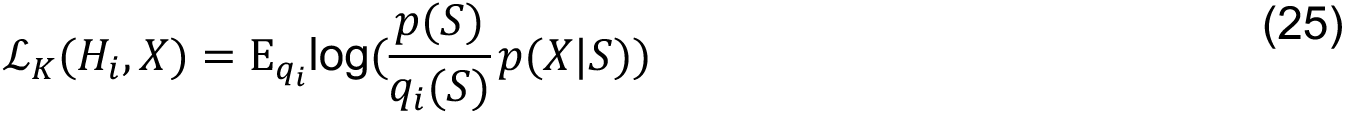

This constitutes a lower bound on the log marginal probability, ℒ*_K_*(*H_i_*, *X*) ≤ log *p*(*H_i_*, *X*). We used Adam to implement stochastic gradient descent. Each iteration of gradient descent used a batch of 10 samples from *q*. The learning rates for the different latent variables were scaled so that the gradient steps taken in any direction were approximately the same size. We also used a scheduler to automatically decrease the learning rate if the variational lower bound plateaued.

## Sequential inference

We now describe the unique aspects of sequential inference, starting with event proposal (step 1) and then source construction (step 2). We provide examples of sequential inference for two different sound mixtures in Supplementary Figures 14 and 15.

### Event proposals via amortized inference – segmentation neural network

A segmentation neural network proposed candidate events from the observed sound for use in later stages of sequential inference. This network was adapted from a previously published image segmentation network to work with sounds (155). It takes in a sound represented as a cochleagram image and outputs a set of (soft) image masks over the cochleagram. However, events in our model are not equivalent to cochleagram regions, rather, they comprise a set of latent variables as described above (Methods, Generative model: Sources and events). Therefore, along with each mask, the network outputs a set of event variables (e.g., onset, offset, amplitude trajectory). We call each set of event variables a “candidate event„. These are only proposals, and may or may not be utilized in later stages of inference (i.e., the inclusion of a candidate event may lower the posterior probability of a hypothesis and thus would be discarded).

Candidate events are depicted in Supplementary Figures 14A and 15A, where they are illustrated as a red mask overlaid on the sound mixture cochleagram. Although not directly depicted in the figures, a candidate event also included its estimated sound type and event variables (onset, offset, amplitude, f0, spectrum). During the subsequent sequential inference rounds, these event variables were combined into full scene hypotheses (shown in Supplementary Figures 14B and 16B/C). We aimed to generate a moderate number of candidate events with the neural network: too many proposals cause sequential inference to become computationally intractable, while too few proposals could increase the possibility that search misses important parts of hypothesis space.

In the sections that follow, we explain how we adapted the image segmentation network to work with sound and to output latent variables in addition to masks.

#### Architecture

The segmentation neural network was based on a publicly available image segmentation network, *Detectron2* (155). We use the standard *Detectron2* Generalized R-CNN/FPN architecture, but modified the training and test procedures to adapt *Detectron2* for use with sounds and in the context of sequential inference, as explained below.

#### Training dataset

We sampled a dataset of input/output training pairs from the generative model. The base representation for the input is a scene cochleagram, *C_S_*. The outputs are event variables {*e_ij_*.} and a set of binary masks of the cochleagram for each event *M_e_*}. Because data generation for training the segmentation network does not need to occur iteratively, we used a more computationally expensive and detailed cochlear model to compute *C_S_* and *M_e_*} (104).

The input to the network was composed of three channels: 1) *C_S_* scaled to image values 0-255 (scaling from 20-180 dB relative to the internal model reference), 2) a binary channel indicating where *C_S_* had energy above the 20 dB threshold, and 3) a binary channel with a value of one for all pixels in the cochleagram. This third channel functioned to encode the edges of the cochleagram. The top and bottom edges corresponded to the ends of the frequency range. Without this third channel, we found that the network would assign events at these edges of the cochleagram, presumably due to spurious features created by zero-padding.

The dataset was composed of 100000 sounds, with durations uniformly sampled from between 0.5 and 4 seconds long. Rather than sampling from the model with source priors as defined by the natural sound datasets, we use uniform priors over all source prior parameters to increase data diversity and thereby improve transfer (domain randomization (157–159)).

#### Training objective

Using these three channels, the network was trained to recover the set of binary event masks *M_e_*} and the sound type and event latent variables {*e_ij_*.} for each event. The loss function used to train the network was the sum of the standard composite segmentation loss and a custom loss based on prediction error for the event latent variables {*e_ij_*.}. This additional component of the loss function was based on the following set of latent variable predictions. For fundamental frequency, we predicted the cochleagram frequency bin with the closest center frequency to the fundamental. For tone amplitude, we predicted the continuous value from the generative model for each time point. For noise and harmonic events, we predicted a continuous value for each time-frequency bin corresponding to the sum of the spectrum and amplitude values from the generative model. For binary-valued outputs (*M_e_*}, frequency) we used a binary cross entropy loss, as is standard for image segmentation. For continuous-valued outputs (tone amplitude, spectrum+amplitude) we used a mean-squared error loss. We defined the left and right edges of the target bounding box to equal the onset and offset of the event. The lower and upper edges of the bounding box equaled the frequencies at the 1^st^ and 99^th^ percentile of the pixels selected by the binary mask.

*Detectron2* also gives a confidence score for each output event proposal. This network confidence score (or associated rank) is listed above each event mask in Supplementary Figure 14A. We trained the network with *Detectron2*’s default optimizer (stochastic gradient descent with learning rate = 0.001, momentum factor = 0.9) for 100000 iterations with a batch size of 20.

#### Test procedure

At test time, we input a scene cochleagram into the segmentation network to determine a list of candidate events for the entire sound. To address resource limitations in downstream sequential inference, we reduced the number of proposed events by excluding some of the events detected by the segmentation network. Given *Detectron2*’s initial outputs, we first applied a custom threshold on the confidence scores that depended on sound type (whistle=0.1, noise=0.1, harmonic=0.7). This reflected our observation that the network more often outputted erroneous proposals of harmonic sounds, impairing inference. Even with these custom thresholds, the network still sometimes detected more events than seemed optimal, in particular with multiple candidates that were near-duplicates (as is common in machine vision segmentation algorithms). Machine vision segmentation algorithms typically use a heuristic technique termed “non-maximum suppression” (NMS) to exclude duplicate objects: if the intersection-over-union (IoU) of a pair of bounding-boxes exceeds some threshold, the lower-confidence candidate is rejected (for a discussion, see (160)). Accordingly, given the initial outputs of the network, we first applied *Detectron2*’s built-in bounding-box non-maximum suppression threshold. Then, for the remaining proposals, we applied a custom non-maximum suppression-inspired “mask threshold”, which instead used the sigmoid outputs for the binary masks of events *e_i_* and *e_j_* in the following IoU formula:

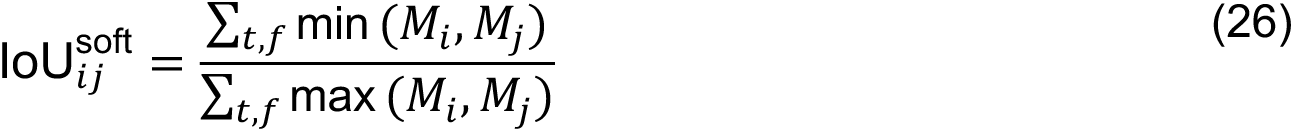

We ran this procedure with both bounding-box and mask thresholds equal to 0.5. This resulted in a list of candidate events with corresponding latent variables for an entire sound, ordered by network confidence.

For some sounds, the procedure described so far resulted in fewer than ten candidate events. In these cases, it was computationally feasible and desirable for inference to include additional candidates to be considered during sequential inference. Therefore, we supplemented the first set of candidate events with an additional set. The additional candidates were computed from the same network, but (1) a more lenient bounding-box non-maximum-suppression threshold (0.9) was used for all sound types, (2) when considering a pair of harmonic candidate events, the mask threshold was replaced with an equivalent IoU computed for the estimated fundamental frequencies (allowing harmonic proposals that differed in fundamental frequency despite overlapping in the cochleagram), and (3) new candidate events were rejected if they duplicated candidate events from the first set. This additional step could never decrease the total number of candidates or alter existing candidates.

In summary, our custom adaptations of *Detectron2* are:

- Self-supervised training on generative model samples, using domain randomization
- Custom input channels to indicate the edges of the cochleagram
- Losses to include the estimation of sound type and event latent variables
- Custom non-maximum suppression inspired mask thresholds
- Flexible procedure to accommodate additional candidate events to be considered in sequential inference

### Source construction

Source construction is the second step of bottom-up inference in analysis-by-synthesis (Step 2 in Inference overview section). In source construction, candidate events are combined to create scene descriptions that each completely specify a set of sources and the sequence of events they emit. These scene descriptions are used to initialize variational inference as described in the above section *Hypothesis optimization and comparison*. Alternations of source construction and variational inference proceed until the entire duration of the sound is observed.

Inspired by sequential Monte Carlo (41, 161), each round *j* of the inference procedure considers a progressively longer duration of the observed sound. On each round, inference provides a set of likely scene descriptions that explain the sound up to time *t_j_*. The result of the first round of source construction is depicted in Supplementary Figure 14B (“Initial hypotheses”). Each initial hypothesis is a full scene, with one or more sources composed of candidate events, along with initial source parameters.

One round of source construction involves 1) any candidate events that have estimated onsets *t_j_*_-1_ < *τ^on^* ≤ *t_j_* (Supplementary Figures 14A and 15A) and 2) the existing scene descriptions selected on the previous round. To create a new scene description, one or more of these candidate events may be added to an existing scene or used to start a new scene. An event can be added to a new source, or it can be added to a pre-existing source as long as the sound type matches and it only minimally overlaps with the other events in the source (IoU ≤ 0.05). The new scene description is constructed and then passed onto the hypothesis optimization step. Specifically, the scene description *S_i_* is used to initialize the means of the guide distribution *q_i_* described above in the above section *Hypothesis optimization and comparison*, corresponding to hypothesis *H_i_*. Supplementary Figure 14B depicts this process over several rounds. Similarly, Supplementary Figure 15C depicts how the existing scene descriptions selected on Round 1 are combined with the last remaining event proposal on Round 2 (see +*e*_4_ in Supplementary Figure 15C) to create a new set of initial hypotheses.

These simple update actions can result in a large number of scene hypotheses if there are many candidate events within the interval [*t_j_*_-1_, *t_j_*]. On some rounds of sequential inference, heuristics were necessary to limit which scene hypotheses were optimized for further consideration. These heuristics essentially favored smaller scene descriptions or ranked hypotheses based on the outputs of the segmentation network (IoU and confidence). They were as follows:

1. Only consider a maximum number of candidate events on each round (max=5), ranked by network confidence.
2. Only consider a maximum number of scenes per round (max=100 for illusions, max=15 for everyday sounds). Prefer scene hypotheses for which events have high network confidence and low IoU between the old and new event(s).
3. After the first round, only consider hypotheses which utilize the events inferred in previous rounds.
4. Do not consider scene hypotheses with more than a maximum number of sources (max=4).
5. After the first round, do not consider scene hypotheses which add more than a maximum number of new sources (max=2).
6. Do not consider scene hypotheses which add more than a maximum number of new events (max=3) on a single round.

We found these heuristics to be essential to making inference tractable when applied to everyday sounds.

### Hypothesis optimization and scene selection (sequential inference details)

After each iteration of the source construction procedure described above, the “initial hypotheses” *qi* were optimized with variational inference resulting in “optimized hypotheses” which had optimized event variables and source parameters. Supplementary Figure 15A depicts this process of hypothesis optimization (the change to the event variables is evident in the change in the rendered cochleagrams before and after optimization). Because there could be many hypotheses to optimize, we first performed a partial optimization to find the most promising hypotheses and abandon the rest. We first used 250 iterations of variational inference as an initial assessment of the hypotheses. We then ranked the hypotheses by their variational lower bound (Methods: Equation 25) and kept only the top five (not shown in figures). Then, for these five hypotheses, we ran additional steps of variational inference. As discussed in the Discussion, illusions typically required additional steps of inference in order to distinguish the competing hypotheses, which often differed in subtle ways. Therefore, variational inference was run for 2000 total iterations for everyday sounds and 4000 total iterations for illusions. Finally, the next round of source construction required the best hypotheses of this round to build upon; the top two hypotheses were selected to be used in the next round of source construction (Supplementary Figures 14B and 15B). This number of hypotheses was constrained by computational limitations. Supplementary Figure 14B shows optimized and selected hypotheses over several rounds of sequential inference (comprising both source construction and hypothesis optimization), demonstrating how the final top hypothesis for the observation was successively built up over several rounds.

### Cleanup proposals

After completing inference on everyday sounds, we generated a few extra sets of hypotheses (automatically, from the hypotheses produced by the inference procedure). These hypotheses were meant to alleviate posterior degeneracy and test alternative minima that were difficult to reach through local gradient-based search. They were generated by (1) removing a source, (2) removing an event, (3) changing a source’s sound type, (4) merging sources, and (5) merging events. In practice, we found that these cleanup proposals had only a modest effect. This is possibly because the discovered hypotheses had already undergone multiple rounds of optimization during sequential inference, making it unlikely that new hypotheses could be optimized as effectively.

### Computational resources

Sequential inference required considerable computational resources. Over the course of a single sound, hundreds of hypotheses could be tested. All hypotheses were optimized on a single GPU over a duration ranging from 2 minutes to an hour. Therefore, sequential inference for a single sound could potentially take on the order of tens or hundreds of GPU hours. This computationally intensive inference was enabled by parallel computing.

## Enumerative inference

In enumerative inference, rather than proposing hypotheses sequentially based on the data, we directly defined hypotheses corresponding to the choices provided to a human participant in a psychophysical experiment. In some cases an experiment required a participant to choose between multiple explanations of sound, corresponding to structurally distinct hypotheses in our model:

- Continuity illusion: a single long whistle vs. several short whistles
- Tone sequences (bistability, build-up of streaming, and the two context experiments): the arrangement of events into one or more sources (streams)
- Co-modulation masking release, Mistuned harmonic: the presence or absence of a whistle event

Two other experiments involved judgments of sound properties that our model treats as continuous latent variables:

- Spectral completion: the spectrum level of part of the target sound
- Onset asynchrony: peaks in the spectrum (used to make a vowel judgment)

Whether the final judgment was based on the structure of the hypotheses or on continuous latent variables, we performed enumerative inference in the same way. We first created hypotheses corresponding to each distinct “structure” to be considered in the experiment (such as the arrangement of elements into sources). For each of these structural hypotheses, we then optimized the continuous latent variables corresponding to the sources and events defined by the hypothesis (such as the spectrum of each source in the hypothesis). Specifically, as described above in the Methods section on Hypothesis optimization and scene selection, we use gradient descent to optimize a “guide distribution” to approximate the posterior distribution over continuous latent variables. As this optimization can land in local optima, we repeated it from multiple initializations (corresponding to different settings of the continuous latent variables). Finally, we use these optimized guide distributions to make the relevant psychophysical judgment. To choose between different structural hypotheses, we compared them based on their marginal probability, which we estimated via importance sampling through the optimized guide distributions (we took the marginal probability to be the maximum over all initializations, to avoid being biased by local minima). To extract continuous latent variables from a hypothesis, we simply took the expectation of the optimized guide distribution (weighting initializations by their importance-sampled marginal probability, which in practice was often similar between initializations).

### Experimenter-defined hypotheses

For each hypothesis, we initialized the continuous event latent variables so that the initial rendered scene sound would be a relatively close match to the observed sound. This was done on a case-by-case basis based on our knowledge of the stimulus parameters and the generative model. For instance, the onsets of the source events were set equal to the onsets of the stimulus components, and the frequencies of the events were set equal to the frequencies of the stimulus components. However, the rendered stimulus was typically not exactly faithful to the original stimulus, because it was often not obvious how to set the generative model parameters by hand to exactly replicate aspects of the stimulus. A simple example of this is the fact that the model used fixed duration onset and offset ramps for events, whereas these varied somewhat in shape and duration in the experimental stimuli. The best fitting onset and offset times thus varied somewhat depending on these ramps and the initialization was not always optimal. It was thus critical to optimize these hypotheses to give the model the best chance of explaining the experimental stimulus. In addition, in order to estimate the probability associated with each hypothesis, we had to optimize the continuous source parameters and estimate posterior variances on all latent variables. For brevity, we omit the long list of generative parameters used for the hypothesis initializations in each experiment, but they are available in the Github repository for the experiments, which will be made public upon publication.

### Hypothesis optimization (enumerative inference details)

In enumerative inference, we used additional steps of variational inference (compared to what was used in sequential inference) to help ensure precise estimates of the posterior distribution for pairs of hypotheses that sometimes differed in subtle ways (see Discussion). Specifically, we used 8000 iterations of variational inference for a single hypothesis (in comparison to the 4000 iterations used during sequential inference).

### Computational resources

Similar to sequential inference, enumerative inference required considerable computational resources. Depending on the experiment, there were 1-10 hypotheses for each sound. All hypotheses were optimized on a single GPU over a duration ranging from 30 minutes to two hours. Therefore, enumerative inference to test all the hypotheses for a single sound could take on the order of tens of GPU hours. Again, this computationally intensive inference was enabled by parallel computing.

## Classic auditory scene analysis illusions

### Overview

We simulated a set of classic psychophysical experiments and illusions in our generative model. For the results reported in the main text figures, we used a mix of enumerative or sequential inference. For illusions that were evaluated subjectively, we used sequential inference to simulate unconstrained listening. For all psychophysical experiments except one, we used enumerative inference, on the grounds that the experiments explicitly asked participants to choose between different perceptual interpretations. The one exception was the cancelled harmonics experiment, where it seemed more appropriate to use sequential inference (see below).

To simulate an experimental setting in which multiple participants complete the same experiment, we repeated enumerative inference multiple times for the same experiment. Specifically, we ran the enumerative inference procedure on all hypotheses for a stimulus 10 times with different random seeds. Each run of the procedure yielded a distribution over the continuous variables for each hypothesis. This distribution could be used to estimate the “perceived” value of one of the continuous variables (e.g. the spectrum level of part of a source). The distribution could also be integrated out to estimate the marginal probability of that hypothesis (e.g., to choose between alternative hypotheses). For each run of the inference procedure, we computed the quantity to be plotted (this varied from experiment to experiment, see below). We then calculated the mean response across inference runs and calculated standard error bars. With sequential inference (intended to simulate unconstrained listening conditions), we ran the inference procedure with a single random seed.

Some experiments (continuity illusion, co-modulation masking release, onset asynchrony, and mistuned harmonic) required estimating the threshold of a stimulus-generation parameter at which the model preferred one explanation to the other. While it would be natural to define this threshold as the point at which the posterior probability crosses 50%, this estimator is not robust to the noisy probability estimates produced by our stochastic inference procedure (and does not necessarily produce a unique value). Instead, we defined the threshold more robustly based on the integral of the posterior with respect to the stimulus-generation parameter. Specifically, if H0 is to be preferred for low parameter values and H1 for high parameter values, we define the threshold τ (uniquely) with respect to stimulus-generation parameter values ν*_X_* such that:

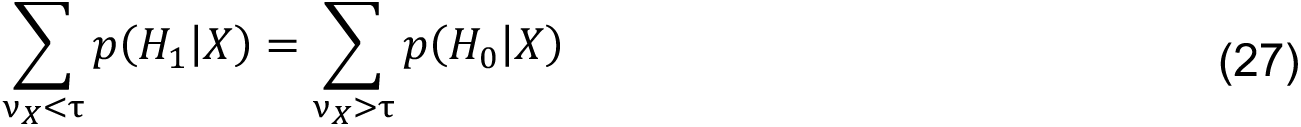

All sounds were generated at 20kHz. Stimulus levels for the model experiments are specified in dB relative to an arbitrary model reference value.

### Continuity illusion

#### Stimuli

We adapted the experimental stimuli from Experiment 3 of ref. (66). In the original experiment, participants adjusted the level of a tone until it was just audible in noise (to measure masking thresholds) or until it just sounded discontinuous, if alternating with noise bursts (to measure continuity thresholds). In both conditions, noise was played throughout the adjustment process. The experiment included 15 participants who had not participated in previous experiments in the paper.

Because our model was not set up to actively adjust the parameters of the experimental stimuli like participants could, we instead generated stimuli spanning a range of tone levels (44-80dB in 4dB increments) and then simulated model responses at each level. We tested model responses for pure tones at 250, 500, 1000, 2000 and 4000 Hz, replicating all the frequencies in the experiment except 8000 Hz (due to its presence on the very edge of the cochleagram representation we used). For both conditions, we generated pink noise with an order-206 FIR filter with a one-octave notch. The filter’s 3dB down points were at 722 and 1367 Hz, and it reached its maximum attenuation of 40dB between 900-1200 Hz. After filtering, the level of the noise was set to 80 dB.

To generate the stimuli for the masking condition, three pure tones were embedded in a noise masker. The masker was a single noise burst of duration 1.8-s with 10-ms raised cosine onset and offset ramps. Each tone was 300-ms long with 10-ms raised cosine ramps. The tones were separated by 300-ms interstimulus intervals. The first tone began 150-ms after the onset of the noise burst, such that the last tone ended 150-ms before the offset of the masker. To generate the stimuli for the continuity condition, the tones were alternated with short noise bursts. Five 300-ms noise bursts were generated with an interstimulus interval of 300-ms. The first 150-ms tone began 150-ms after the offset of the first noise burst. The next two noise bursts were alternated with 300-ms tones. The last tone started immediately after the fourth noise burst and ended 150-ms before the onset of the last noise burst. The noise and the tones each had 10-ms onset and offset ramps, and overlapped in the region where they were ramped (i.e., were cross-faded). All stimuli were padded with 50ms of silence. In total, there were 50 stimuli resulting from this process (5 frequencies ✕ 10 levels). The above process was replicated 10 times with different exemplars of noise for each replication of inference, increasing the stimulus set size to 500 (10 exemplars of 50 stimuli).

#### Analysis (enumerative inference)

We used a one-interval, 2AFC task to measure thresholds. For each stimulus in the masking condition (corresponding to a tone level/frequency pair), the model compared the hypotheses that tones were present versus absent. The tone-absent hypothesis was initialized with a noise with a similar spectrum to the stimulus masking noise (as described above, we attempted to set the parameters of the generative model to match the noise, but this did not produce an exact match, and was optimized during the model inference to match the stimulus as best possible). The tone-present hypothesis initialized this noise along with the tones at the appropriate level, frequency and timing for the stimulus. For each stimulus in the continuity condition, the model compared the hypothesis that there was a continuous single tone versus four discontinuous tones. Each hypothesis was initialized with noise bursts of appropriate duration and spectrum, and tones of constant level. For each condition, this resulted in a log odds curve as a function of tone level for each tone frequency, which we used to calculate thresholds.

### Homophonic continuity

#### Stimuli

We extracted two 1.5-s clips from Track 32 of (67), titled “Homophonic continuity and rise time”. Both clips were composed of continuous bandpass white noise (0-8 kHz), but had different amplitude envelopes. The initial amplitude of both sounds was a quarter of the peak amplitude, and both sounds rose and fell in amplitude twice. The first clip was gradually modulated, rising linearly to a peak over 252 ms and then falling to the initial amplitude over 252 ms. The second clip was abruptly modulated, rising to peak amplitude in 1 ms, maintaining peak amplitude for 32 ms, and then falling to the initial amplitude in 1 ms. We downsampled the original clips to a sampling rate of 20kHz. Each sound was padded with 50-ms of silence.

#### Analysis (sequential inference)

The test of whether the model succeeds in this illusion is whether the abruptly modulated noise is interpreted as two separate sources despite the gradually modulated noise being interpreted as a single source. Therefore, we only used sequential inference for these sounds and visually assessed the results.

### Spectral completion

#### Stimuli

We reproduced the experimental stimuli from Experiments 1 and 2 of ref. (68). In the original experiment, participants adjusted the spectrum level of the middle frequency band of a short comparison noise until it sounded as similar as possible to the target noise. The starting level of the middle band of the comparison noise was set randomly between -10 and 30 dB spectrum level. The final spectrum level of the middle band was reported as an average across participants. The experiments included 8 participants between the ages of 18 and 30 years old.

Stimulus generation was identical to that in the original experiments. Each stimulus was generated by combining bandpass white noise bursts of various spectrum levels and pass bands. The standard stimulus consisted of a “masker” and a “target”. The masker was 750 ms in duration and had a pass band of 500-2500 Hz. The target contained a lower tab (100-500 Hz) and an upper tab (2500-7500 Hz). The target was 150 ms in duration, and started 300 ms after the onset of the masker. To generate each noise burst, we first set all spectral domain magnitude coefficients outside the pass band to zero, and then performed an inverse fast Fourier transform. Each noise burst had 10-ms raised cosine ramps. We additionally padded each stimulus with 100ms of silence.

For Experiment 1, we generated five stimuli. The tabs in Experiment 1 were set to a spectrum level of 20 dB/Hz. Stimuli i and ii only contained the short target sound. In stimulus i, the middle band was silent. In stimulus ii, the middle band had a spectrum level of 30 dB/Hz. Stimulus iii was the standard stimulus described above. Stimulus iv had a masker with a spectral gap that extended from 600-2080 Hz and its spectrum level was increased so that its overall level was equal to that of the masker in the standard stimulus. Stimulus v had a masker which stopped at the tab onset and began again at the tab offset.

For Experiment 2, we generated six variations on the standard stimulus in which the spectrum level of the tabs and maskers were varied in opposite directions. The tab and masker levels for these six stimuli were (5,35), (10, 30), (15, 25), (20, 20), (25, 15), and (30, 10) dB/Hz.

There were a total of 11 stimuli across both experiments. We replicated the stimulus generation process 10 times with different exemplars of noise for each replication of inference, increasing the stimulus set size to 110 (10 exemplars of 11 stimuli).

We also generated each comparison stimulus with a range of spectrum levels for the middle band. For Experiment 1, we generated the comparison stimulus with middle band spectrum levels spanning -15-40 dB/Hz in steps of 2.5 dB/Hz, and tab spectrum levels at 20 dB/Hz. This resulted in a total of 22 comparison stimuli. For Experiment 2, we generated each comparison stimulus with spectrum levels spanning -5-12.5 dB/Hz in steps of 1.25 dB/Hz. For each comparison stimulus, the tab level matched that of the corresponding target stimulus. This resulted in 15 comparison stimuli for each of 6 target stimuli, for a total of 90 comparison stimuli. We replicated this process 10 times with different exemplars of noise for each replication of inference, increasing the comparison stimulus set size to 900 (10 exemplars of 90 comparison stimuli across both experiments).

#### Analysis (enumerative inference)

For each stimulus, we optimized a single structural hypothesis based on the stimulus generation parameters. For the first two stimuli in Experiment 1, the structural hypothesis contained a single noise source corresponding to the target. For the rest of the stimuli in Experiment 1 and all stimuli in Experiment 2, the structural hypothesis contained two noise sources that corresponded to masker and target. For stimulus v in Experiment 1, the source corresponding to the masker contained two events; otherwise, each source contained one event. We note that these structural hypotheses accord with the number of sources and events found by sequential inference except in the case of stimulus iv in Experiment 1, where the masker is accounted for by two simultaneous sources. Nevertheless, for stimulus iv in Experiment 1, the inferred middle band spectrum level is comparable for both enumerative and sequential inference.

The structural hypothesis was initialized to match the masker and tab level in the stimulus, but with multiple settings of the middle band spectrum level of the target. This emulated the variable starting level of the middle band in the experiment. For Experiment 1, the initial spectrum levels of this middle band were -20, 0, 10, 20, and 30 dB/Hz. For Experiment 2, the initial spectrum levels were -5, -2.5, 0, 5, 10, 12.5, 20 dB/Hz. After variational inference, for each initialization, we selected the batch of sampled scenes with the best score (see Methods: Hypothesis optimization and comparison). We averaged the target spectrum over the batch. Then, we took a weighted average of the target spectra (arising from the different initializations) using their importance sampled marginal probability. Then, we computed the mean squared error between this average spectrum and the inferred latent spectra for each comparison stimulus. The model’s judgment was selected to be the spectrum level of the comparison stimulus which minimized the error.

Averaging over runs of inference with different random seeds provided the plotted value and standard error bars.

### Co-modulation masking release

#### Stimuli

We adapted the experimental stimuli from Experiment 1 of ref. (69). In the original experiment, participants were asked to detect a tone in bandpass noise that varied in bandwidth. The different spectral regions of the noise were either co-modulated or had random amplitude envelopes. Participants’ thresholds were measured using a two-interval, 2AFC procedure. The experiment included five highly experienced participants with two hours of training in all experimental conditions.

For each bandwidth, we generated a co-modulated noise burst and a random noise burst. The random noise burst was generated from white noise (with power from 0-10 kHz, i.e. up to the Nyquist frequency). To generate co-modulated noise, we multiplied the white noise by a low-pass noise with power between 0-10 Hz. Then, we bandpass filtered both noises to the appropriate bandwidth, centered on 1000 Hz, using a fourth-order Butterworth filter and forward-backward filtering. After filtering, we confirmed that the absolute spectrum level of the co-modulated and random noises were within 1 dB/Hz of 40 dB/Hz within the passband. We trimmed each noise to 400-ms and applied 50-ms raised cosine ramps. We then generated a set of 1 kHz tones with a range of levels (40-85 dB in steps of 5 dB). The tones were 400-ms with 50-ms raised cosine ramps. We added the tones to each noise, for a total of 20 stimuli for each noise bandwidth (2 noise conditions ✕ 10 tone levels). We repeated this process for 100, 200, 400, and 1000 Hz bandwidths, resulting in 80 stimuli. Finally, we repeated this process ten times with different exemplars of noise for each replication of inference, increasing the stimulus set to 800 sounds.

The major difference between the original experiment stimuli and our model simulation is that the frequency cutoff of the low-pass noise for the model experiment was 10 Hz, instead of the original 50 Hz. We used slower amplitude modulations in the stimuli for the model experiment because the relatively coarse temporal resolution of the cochleagram representation limited the rate of modulations that could be resolved. We found subjectively that the masking release was still obvious at this setting, consistent with previous work (71). We also omitted the 25 and 50 Hz bandwidth conditions due to the frequency resolution of our cochleagram.

#### Analysis (enumerative inference)

For each stimulus, we compared the hypotheses that a tone was present versus absent (one-interval, 2AFC task). For all hypothesis initializations, we initialized the noise with the appropriate amplitude envelope. The tone absent hypothesis only contained the noise burst. For the tone present hypothesis, we made two initializations in order to aid in finding the best setting of the continuous latent variables for this hypothesis. One initialization had the tone matching the tone level in the stimulus. The other had a “quiet” tone initialized at 0 dB. For each bandwidth and noise condition, we computed the log odds as a function of tone level then computed the detection threshold.

### Frequency modulation

#### Stimuli

We based our stimulus design on the classic demonstration first described in ref. (72). We generated a 1-s complex tone in which the odd harmonics had a steady f0 and the even harmonics were coherently frequency modulated. The even harmonics began in a harmonic relationship with the odd harmonics, but were immediately frequency modulated at a rate of 2 Hz, with a maximum frequency change of 70 Hz. The fundamental frequency of the odd harmonics was 300 Hz. The stimulus contained harmonics 1-12, with levels of 75, 63, 56, 52, 48, 48, 42, 46, 38, 45, 35, 39 dB respectively. The entire stimulus had 20-ms raised cosine ramps, and was padded with 50-ms of silence.

#### Analysis (sequential inference)

The test of whether the model succeeds in this illusion is whether the modulated components are discovered as a separate source. Therefore, we only used sequential inference for these sounds and visually assessed the results.

### Mistuned harmonic

#### Stimuli

We reproduced a subset of the experimental stimuli from ref. (76). In the original experiment, participants were presented with a complex tone and asked to indicate whether they heard a single sound (with one pitch) or two sounds (a complex tone and a pure tone). On half of trials, the tone was harmonic and on the other half one frequency component was mistuned. The authors then calculated the degree of mistuning which was necessary to detect two sounds. The experiment included four participants, the authors and one volunteer, who were all highly experienced with psychoacoustics and with the task of detecting mistuning in harmonic complexes.

We generated 400-ms complex tones with a fundamental frequency of 100, 200 or 400 Hz. The complex tones had equal amplitude harmonics, each with a level of 60 dB. Tones at 100 and 200 Hz included harmonics 1-12 and tones at 400 Hz included harmonics 1-10. Each tone was given 10-ms raised cosine onset and offset ramps. Based on these harmonic complex tones, we created complex tones where one component was mistuned. We mistuned harmonics 1-3 by 5, 10, 20, 30, 40, or 50% of the fundamental frequency. Including the in-tune harmonic complexes (0% mistuning), this resulted in 63 stimuli (3 fundamental frequencies ✕ 3 harmonic numbers ✕ 7 mistuning levels). Every stimulus was padded with 50-ms of silence.

#### Analysis (enumerative inference)

For each stimulus, we compared the hypothesis that there was one harmonic source versus the hypothesis that there was one harmonic source and one whistle source (one-interval, 2AFC task). For the single source hypothesis, we initialized a single harmonic source with the stimulus parameters of the in-tune harmonic complex. For the two source hypothesis, we made two initializations in order to aid in finding the best setting of the continuous latent variables for this hypothesis. Both initializations had a whistle source at the mistuned frequency and a harmonic source with the fundamental frequency of the stimulus, but they varied in the relative energy of the whistle and the component within the harmonic source that corresponded to the mistuned harmonic number. In the first initialization, the component corresponding to the mistuned harmonic was attenuated by 30 dB in the harmonic source, and the whistle source was set to 60 dB. In the second initialization, the component corresponding to the mistuned harmonic was only attenuated by 6 dB, and the whistle source was set to 50 dB. For each fundamental frequency and harmonic number, we computed the log odds as a function of mistuning percent then computed the detection threshold.

### Asynchronous onsets

#### Stimuli

We reproduced a subset of the stimuli from Experiment 1 in ref. (77). In the original experiment, participants heard short speech-like sounds and categorized them as either /I/ or /e/. The experiment included six participants, who had practice with the control conditions.

We generated four of the continuua from the original experiment. The first was the ‘basic’ continuum, which was composed of seven vowels for which the first formant was varied from 375 to 500 Hz in equal steps. The other three continuua were created by adding 500-Hz tones to the basic continuum. The first ‘shifted’ continuum was created by adding a tone with the same onset and offset as the vowel. The onset of the tone in the other two continuua was asynchronous with the vowel, either 32-ms before or 240-ms before the vowel. To generate the vowel sounds, we used a publicly available Python interface of the Klatt speech synthesizer, which was used in the original experiment (162, 163). To create the basic continuum, we generated seven 60-ms long vowels with a fundamental frequency of 125 Hz and overall level of 56 dB. In each vowel, formants 2-5 were centered at 2300, 2900, 3800, and 4600 Hz respectively. The bandwidths of these formants were unspecified in the original paper and were set to the Klatt synthesizer defaults (70, 150, 200 and 200 Hz for formants 2-5, respectively). The bandwidth of the first formant was 70-Hz and kept constant in each vowel. The first formant frequency varied across vowels, with values of 375, 296, 417, 438, 459, 480 and 500 Hz. These values are referred to as the “nominal first formant frequencies” for the other continuua. Each vowel had 16-ms linear onset and offset ramps. To generate the tones to add to each vowel in the basic continuum, we first measured the level of the 500 Hz component in each vowel. For each vowel, we generated a pure tone that was 6-dB higher in level than the 500-Hz component. This pure tone constructively interfered with the 500 Hz component of the vowel to produce a 9.5 dB increment from the original level at 500 Hz. After adding 16-ms linear onset and offset ramps to the tone, we added the tone to the vowel so as to produce the desired onset difference. This process resulted in 28 stimuli (4 continuua ✕ 7 vowels). All stimuli had the same overall duration, and were zero-padded so that the vowel had the same absolute onset time in each stimulus. The stimulus with the largest onset asynchrony was zero-padded with 50-ms of silence.

#### Analysis (enumerative inference)

For each stimulus, we found the frequencies where the first two ‘formant’ peaks occurred in the inferred spectrum of the harmonic source. We then computed classification probabilities according to empirical first and second formant distributions measured by (78). This enabled us to compute the vowel boundary (classification threshold) in terms of the nominal first formant frequency.

For each stimulus, we considered the case where the 500-Hz component is explained by the harmonic vowel source and the case where it is (at least partially) explained by a whistle source. For each stimulus in the basic and shifted continua, we optimized two structural hypotheses: (1) a harmonic source only and (2) a harmonic source and a simultaneous 500-Hz whistle source, initialized with the stimulus parameters. For the asynchronous onset continuua, we optimized a single structural hypothesis (a whistle source and a harmonic source) from two initializations. The first initialization included a harmonic source with the shifted spectrum and a whistle that ended at the beginning of the harmonic source. This corresponded to the possibility that the overlapping 500-Hz component is grouped with the harmonic source. The second initialization included a harmonic source with the basic spectrum and a whistle that ended synchronously with the harmonic source. This corresponded to the possibility that the 500-Hz component of the harmonic event is ‘captured’ by the asynchronous whistle.

After variational inference, for each initialization, we selected the batch of sampled scenes with the best score. To estimate the formants of the inferred vowel, we averaged the harmonic source’s latent spectrum over the batch, and then found the frequencies at which this inferred spectrum had peak amplitude in the ranges 325-500 Hz and 2000-2500 Hz, corresponding to the first and second formants. For each formant, we additionally took a weighted average of the formants arising from the different initializations using their importance-sampled marginal probability.

We then selected a subset of the data in (78) corresponding to the vowels /I/ and /e/. To compensate for potential differences between the speakers in the original vowel set and those which the synthetic vowel stimuli were modeled on, we normalized both the empirical formant distribution and inferred formant distribution (164) by z-scoring. We z-scored the inferred formants using the mean and standard deviation computed across all conditions and seeds. We z-scored the empirical formants using the mean and standard deviation of the whole selected subset of formants. We then separately computed the mean and covariance of the z-scored /I/ and /e/ formant distributions.

To derive classification probabilities for the model inferences, we computed the probability of each z-scored, inferred formant pair under two normal distributions with the empirical means and covariances. For each continuua, this provided a classification probability as a function of nominal first formant frequency. Finally, we computed the vowel boundary threshold. We took a mean of the vowel boundary over the different runs of the inference procedure and computed the standard error.

### Cancelled harmonics

#### Stimuli

We adapted the stimuli from Experiment 1 of ref. (79). In the original experiment, listeners matched the frequency of a comparison tone to their percept of a gated harmonic within a harmonic complex tone. The experiment included four male listeners between the ages of 21-65; two were the authors.

We created harmonic complex tones with fundamental frequencies spanning 190 to 210 Hz in five equal steps (in Hz). The duration of each tone was 750-ms, with 10-ms raised cosine onset and offset ramps. Each tone contained harmonics 1-30. The harmonics were each 45 dB and added in sine phase. For each fundamental frequency, we created a set of stimuli each with a different gated harmonic component (harmonics 1-3,10-12,18-20). We gated the component with 10-ms raised cosine ramps to create five tones of 100-ms each. The onset of the first tone and the offset of the last tone were aligned to those of the harmonic complex. Therefore, the interstimulus interval between the tones was 62.5-ms. The entire stimulus was padded with 50-ms of silence. This process led to a total of 45 stimuli (5 fundamental frequencies ✕ 9 harmonic numbers).

The stimuli for the model experiment were shorter than those for the original experiment. The original experiment used 9.1-s long harmonic complexes with four tones of 1.3-s. Inference with this stimulus duration would have been prohibitively computationally expensive, so we instead used 750-ms long harmonic complexes with five tones of 100-ms.

#### Analysis (sequential inference)

The test of whether the model succeeds in this illusion is whether it discovers any whistle sources that correspond to the gated components. Therefore, we used sequential inference, after which we selected the top-scoring hypothesis which contained any whistle sources. For any whistle source in that scene, we recorded its source-level mean fundamental frequency (µ_B_). We took the average *μ*_B_ across all whistle sources if there were more than one (because this typically corresponded to some of the gated tones being assigned to a different whistle source than others, but with similar frequencies). If no hypothesis contained a whistle source, we recorded this as a ‘no-match’ trial, as in the original experiment. For each trial (corresponding to a harmonic number and fundamental frequency), we calculated the percent matching error as a proportion of the gated component frequency. We also recorded the number of ‘no-match’ trials for each harmonic number summed across fundamental frequencies. We plotted the distribution of errors in Figure 6D.

### Frequency proximity

#### Stimuli

We adapted two tone sequences from Experiment 2 of (81), which are similar to the demonstrations in Track 17 of (67) titled “Failure of crossing trajectories to cross perceptually”. In the original sequences, tones in an ascending sequence are alternated with tones in a descending sequence. In the first clip, all the tones in both sequences are pure tones. In the second clip, the ascending sequence is composed of harmonic tones. Listeners judge whether they can hear the ascending and descending sequences, or whether they hear two ‘bouncing’ sequences.

We interleaved an ascending and descending sequence, each composed of 6 tones with frequencies evenly spaced on a log-frequency scale (504, 635, 800, 1008, 1270, 1600 Hz). Each tone was 100-ms in duration with 8-ms onset and offset ramps, with adjacent tones played back-to-back. In the first version (where all tones were pure tones), all tones were 70 dB. In the second version (where the ascending sequence contained harmonic complex tones), the harmonic complex tones contained the first four harmonics of the fundamental frequency (which were the pure tone frequencies from the first version). The pure tones were 73 dB while each component of the harmonic tone was 67 dB.

#### Analysis (sequential inference)

Since the stimuli from this experiment are often used as standalone demonstrations, we tested whether “bouncing” sequences are discovered in the first clip and “crossing” sequences are discovered in the second clip. Therefore, we only used sequential inference for these sounds and visually assessed the results.

### Bistability

#### Stimuli

We adapted the classic ABA sequences used in Track 3 of (67), titled “Loss of rhythmic information as a result of stream segregation”, in order to estimate the stimulus parameters Δ*f* and Δ*t* that lead to one- or two-stream perceptual organizations. These effects were first measured by Experiment 2.3.2 in ref. (82), but this experiment used 80-s sequences that are computationally infeasible for our model, and so we generated shorter sequences.

In the typical ABA tone sequence, three tones are followed by a silence of the same duration as the tone onset-to-onset interval. The first and third tone in the triplet (A) have the same frequency, which can be different from the frequency of the second tone (B). We generated versions of such sequences in which the onset-to-onset interval was 67, 83, 100, 117, or 150 ms. The A tone was always 1000 Hz, and the B tone was 3, 6, 9, or 12 semitones higher in frequency. The tones were all 70-dB and 50-ms long with 10-ms raised cosine onset and offset ramps. The ABA triplet was repeated four times. The stimuli were padded with silence so their total duration was equal (3.1 s). This resulted in 25 stimuli (5 frequency intervals ✕ 5 time intervals).

#### Analysis (enumerative inference)

For each tone sequence, we compared two hypotheses: one stream (all tones in one source) versus two streams (high tones in one source, low tones in a separate source). We initialized each hypothesis with a set of whistle events with the stimulus frequencies and timings that were organized into the appropriate sources. After variational inference, we computed the log odds of the hypotheses for each sequence. We took the average log odds across all runs of the inference procedure and computed the standard error.

### Buildup of streaming

#### Stimuli

We adapted the tone sequences used in ref. (84). In the original experiment, listeners heard a 12.5-s ABA sequences. At any point during each sequence, they could freely indicate whether they were hearing one or two sources. The experiment included eight listeners between 23-57 years old.

Listeners in (84) could respond at any point during a sequence. To obtain an analogous measure of the effect of time on the model’s inferred perceptual organization, we instead evaluated the model for multiple sequences, each with a different number of repetitions. We generated two sets of ABA sequences (with the tone arrangement described in the previous section, Bistability) one with a frequency difference Δ*f* of eight semitone and one with a frequency difference of four semitones. The A tone was 500 Hz. The tones were 50-ms in duration with 10-ms raised cosine onset and offset ramps, and with 125-ms onset-to-onset intervals. Each ABA triplet was thus 500-ms in duration. We generated sequences with 1-6 repetitions of the ABA triplet. This resulted in 12 tone sequences total (2 Δ*f* ✕ 6 sequence durations).

#### Analysis (enumerative inference)

The initialization of inference, and the analysis, was identical to the previous section, Bistability.

### Effects of context (1)

#### Stimuli

We adapted the stimuli used in Task 1 of Experiment 2 in ref. (14). In the original experiment, listeners first heard the standard AB tone pair in isolation and then 12 repetitions of the ABXY sequence. They rated whether the standard was audible as a separate pair in the sequence. The experiment included 16 young adult participants.

We generated the seven ABXY sequences in the original experiment, but only repeated them four times (because it was computationally prohibitive to run inference on longer stimuli). For the four “isolate” stimuli, the ABXY frequencies were (2800, 1556, 600, 333), (600, 333, 2800, 1556),(2800, 2642, 1556, 1468), and (333, 314, 600, 566) Hz, respectively. For the three “absorb” stimuli, the ABXY frequencies were (2800, 1556, 2642, 1468), (600, 333, 566, 314), and (2800, 600, 1468, 314) Hz, respectively. Each tone was 100-ms with 10-ms sine-squared onset and offset ramps, and 10-ms silences between tones. In the original experiment, all tones were 70-dB except for 333 and 314 Hz, which were 77-dB (to equate the loudness of the tones, based on equal loudness contours at 70 phon). This adjustment was unnecessary for the model, so we presented all tones at 70-dB. In the original human experiment, the sequence was faded in and out to prevent participants from adopting strategies based on the beginning or end of the sequence. This was not an issue with the model, and so all cycles of the sequence were presented at the same level.

#### Analysis (enumerative inference)

For each tone sequence, we compared two sets of hypotheses. In both sets, all hypotheses contained one source with a pair of tones, and one or two other sources. The first set contained the two hypotheses in which A and B were paired in their own source (one with X and Y in a second source, and one with X and Y in different sources). The other set contained the eight hypotheses in which A and B were in separate sources (each with a different assignment of X and Y to the sources containing A and B, or to separate sources). We initialized each hypothesis with a set of whistle events with the stimulus frequencies and timings that were organized into the appropriate sources. After variational inference, we summed the marginal probabilities within each set and then calculated the log odds for each sequence. We averaged the log odds across all runs of the inference procedure and computed the standard error.

### Effects of context (2)

#### Stimuli

We adapted the stimuli used in Task 1 of Experiment 2 in ref. (85). In the original experiment, listeners first heard a standard tone pair and then a longer sequence that contained a “target” tone pair, comprising tones of the same frequency. The target tones could be in the same order as the standard, or in reverse order. The longer sequence also contained two “distractor” tones, one immediately preceding the target pair and one following the target pair. Participants judged whether the standard and the target had the same order. The experiment included 13 participants from ages 16 to 26 years.

There were 4 stimuli in this experiment, each corresponding to a different “captor” condition that varied in the presence of captor tones that might cause distractor tones to segregate from the target tones. In the “none” captor condition, there were only distractor tones and target tones in the long sequence. In the other captor conditions, the distractor and target tones were in the same configuration, preceded by 3 captor tones and followed by 2 captor tones. The first target tone had frequency 2200-Hz and level 60-dB. The second target tone had frequency 2400-Hz and level 60-dB. The frequency of the distractor tones was 1460-Hz and their level was 65-dB. The frequencies and levels of the captor tones in the three conditions with captors were (590-Hz, 63-dB), (1030-Hz, 60-dB) and (1460-Hz, 65-dB) respectively. The duration of each tone was 70-ms (compared to 45-ms in the original experiment), with 7-ms on-ramps and 5-ms off-ramps. Each target tone and the first distractor was preceded by 9-ms of silence and followed by 0-ms of silence. The captor tones and second distractor tone were preceded by 9-ms of silence and followed by 64-ms of silence.

#### Analysis (enumerative inference)

For each tone sequence, we compared the pair of hypotheses where the target tones were in their own source to the hypothesis where they were grouped with the distractors. The pair of hypotheses with the target tones in their own source included the hypothesis where the captors and distractors were grouped and the hypothesis where the captors and distractors were segregated. Within this pair, we summed the marginal probabilities. Then we took the log odds of the targets-alone hypotheses and the targets-plus distractors hypothesis. We averaged the log odds across all runs of the inference procedure and computed the standard error.

## Human-model dissimilarity

To compare how well our model, the lesioned models, and the source-separation neural networks matched human perception (Figure 8A and D), we quantified the dissimilarity with human perception of classic illusions. The dissimilarity was based on correlations between model and human data, and the method for obtaining a correlation coefficient was dependent on the illusion (e.g., subjectively evaluated vs. threshold measurements) and the form of the model output (because the source-separation networks output a set of soundwaves – estimates of the premixture sounds – rather than the symbolic scene description provided by our model). We first give the overall form of the human-model dissimilarity metric, and then describe the method for obtaining each correlation from either a set of output soundwaves or from symbolic scene descriptions.

For each experiment, we computed a baseline correlation using the original stimulus as the “model output”, that is, as though a model inferred a single source identical to the stimulus itself. We then computed the human-model dissimilarity for that experiment normalized by the baseline dissimilarity:

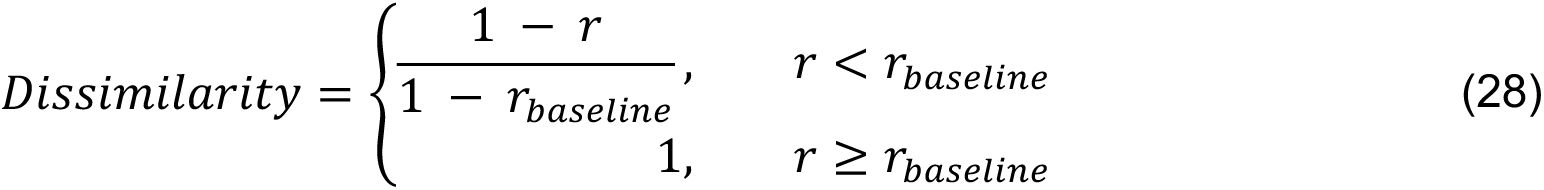

where *r* is the appropriate correlation coefficient as described below. Using this measure, a dissimilarity of zero corresponds to a perfect human-model correlation, while a dissimilarity of one corresponds to the baseline correlation or worse. We then averaged this dissimilarity across all experiments to obtain the aggregate dissimilarity measure plotted in Figure 8A and D.

We use one of three types of correlation coefficient *r* depending on the illusion. They are:

1. Pearson correlation between cochleagrams (across pixels). This correlation was used for illusions that are subjectively assessed (homophonic continuity, frequency modulation, frequency proximity).
2. Pearson correlation between continuous-valued experiment results (across stimulus conditions). This correlation was used for illusions assessed with continuous-valued average judgments: either the preference for one explanation over another (bistability, buildup of streaming, effect of context) or an estimate of a continuous variable (spectral completion, cancelled harmonics).
3. Spearman correlation between thresholds (across stimulus conditions). This correlation was used for experiments that measure thresholds (continuity illusion, co-modulation masking release, mistuned harmonic, and onset asynchrony) because thresholds were sometimes unmeasurable, and were set to a maximum value, violating the assumptions of the Pearson correlation.

Each of these will be explained in more detail below, along with the method for obtaining the correlation depending on the form of the model output. We first describe the forms of model output in more detail.

If the model output was a symbolic scene description (generative models represented in Figure 8A-C), we simulated psychophysical experiments as described above in Methods section *Classic auditory scene analysis illusions* (a combination of enumerative and sequential inference, depending on the illusion). For each lesioned model, we obtained analogous results to Figures 5-7.

If the model output was a soundwave (as for the baseline, the neural networks shown by grey bars in Figure 8D, and the generative model evaluated in the same way as neural networks, shown by the pink bar in Figure 8D), we require a method to simulate psychophysical experiments by obtaining psychophysical judgments from the output soundwaves. For this, we used “standard sets„ of source sounds. For each sound mixture, we defined standard sets that each consists of a set of sounds rendered from a scene hypothesis for that sound mixture. These sounds could be compared to the output soundwaves of a model to obtain psychophysical judgments. They were typically a set of sounds that reflect what human listeners hear (e.g., for frequency modulation: one sound with the frequency-modulated harmonics and one sound with the steady harmonics) or sounds rendered from the experimentally-defined hypotheses of enumerative inference, and are described below for each ASA illusion. For example, in the bistability experiment, there were two standard sets with two sounds each. The first represented the two-stream hypothesis: one sound had all the high frequency tones and the other sound had all the low frequency tones. The second represented the one-stream hypothesis: one sound was the mixture with both high and low tones, and the other sound was silence (because the networks always output at least two sources).

If there were multiple standard sets representing multiple hypotheses, as just described for the bistability experiment, we computed a model’s preference for one hypothesis over another by using the output soundwaves. To determine model preference, we computed the L2 distance between each of the model outputs *N* {*N_i_*}, and each standard set *S_k_* = {*S_kj_*}, in cochleagram space. We found the correspondence of model outputs *i* to standards *j* which minimized this L2 distance for each standard set. If the size of the model outputs was less than the size of the standard set we added a silent model source to allow this distance to be calculated. For experiments which compared two hypotheses, we then compared these minimal distances to each standard set, to provide a measure of the model’s preference for hypothesis H1 over hypothesis H2, specifically:

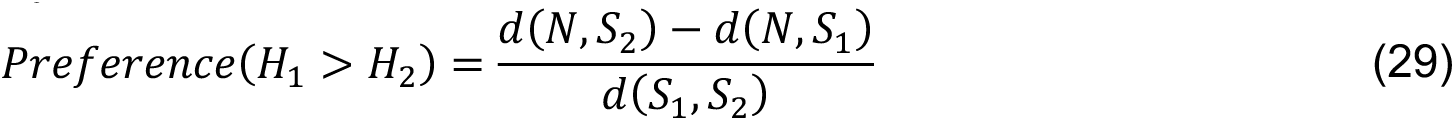

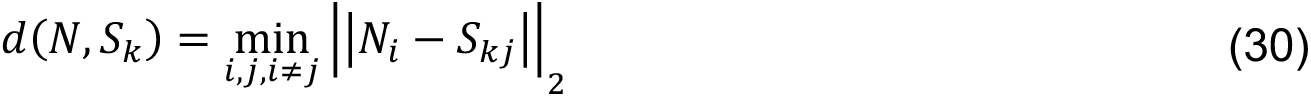

where the denominator is the minimal distance between each set of standards, used to normalize the distance metric. This model preference (based on estimated sound waveforms) was interpreted analogously to the log odds between symbolic scene descriptions in the generative models. This preference was used with correlation types 2 and 3.

We turn now to the three types of correlation coefficient introduced above. The subsequent section will cover each illusion in more detail.

### Correlation type 1, Comparing model outputs to standard cochleagrams

For subjectively evaluated illusions, we computed the maximum Pearson correlation between the cochleagram values of model outputs and the single set of standard sources that human listeners were considered to hear. For generative models with symbolic output, we used the cochleagrams rendered from the inferred sources.

### Correlation type 2, Comparing model and human continuous-valued experiment results

For each stimulus, the model output was used to determine a continuous-valued experiment result. We computed the Pearson correlation between the set of model judgments and the human judgments for all stimuli. In the case of the tone sequences (bistability, buildup of streaming, and effect of context), the model result was the model log odds (if symbolic output) or model preference (if soundwave output) for one explanation over another, and the human result was typically the average result across listeners (but see section on bistability below). In the case of spectral completion and cancelled harmonics, the result was an estimate of continuous variable (spectrum level and frequency). Note that for cancelled harmonics, defining a standard set was not required (see below).

### Correlation type 3, Comparing model and human thresholds

For continuity illusion, co-modulation masking release, and mistuned harmonic illusions, we compared model and human thresholds. For generative models with symbolic output, we used the thresholds as determined by the simulated psychophysical experiments described above. For models with soundwave output, we used the change in model preference as a function of an experimental parameter to estimate a threshold (Methods: Equation 27). The one exception was the onset asynchrony experiment, in which we used the change in an acoustic property (formant frequencies) of the preferred model output to estimate a threshold. The neural networks sometimes did not change the sign of their preference across all stimulus values. In these cases, we assigned the network judgment just above or below the range of stimulus parameters, analogous to setting a threshold to a ceiling or floor value when it cannot be measured in a human participant (we used Spearman rather than Pearson correlation because of the resulting truncated distribution of judgments).

We next define the standard sets for each experiment that were compared to the soundwave outputs to yield simulated experimental results (including for the baseline result), as well as the procedure used to obtain the experimental result. These next sections therefore exclusively apply to models with soundwave output (baseline, neural networks, generative model evaluated in same way as neural networks). The methods for evaluating the symbolic output from generative models have already been described in detail above (in Methods: *Classic auditory scene analysis illusions*).

#### Continuity illusion

For each continuity stimulus, we generated two pairs of standard sources. The first pair contained a sound with the noise bursts on their own and a sound with the (discontinuous) tones on their own. The second pair contained a sound with the noise bursts and a sound with a continuous constant-amplitude tone. For each masking stimulus, we also generated two pairs of standards. The first pair included the input mixture itself (no segregation of tones) and silence. The second pair included a sound with the tones only and a sound with the noise only.

For each condition and tone frequency, the preference measure (Methods: Equation 29) quantified whether continuity was preferred over discontinuity as a function of tone level and whether the tone was detected or not, as a function of tone level. Using correlation type 3, we obtained the Spearman correlation between the model and human thresholds.

#### Homophonic continuity

For the clip with the abrupt amplitude change, we generated a pair of standard sources: a sound comprising just the two short, louder noise bursts, and a sound comprising the long, quiet noise. For the clip with the gradual amplitude change, the standard was simply the stimulus itself. Using correlation type 1, we computed the Pearson correlation between the time-frequency bins of the cochleagrams of the model outputs and the standards, and selected the best match.

#### Spectral completion

The standards were the comparison stimuli described previously (Methods: Classic auditory scene analysis phenomena, Spectral Completion), with tab levels that matched the target and varying middle band spectrum levels. We found the standard which minimized the distance with the best-matching model output. The model’s judgment for a stimulus was chosen to be the middle band spectrum level of the selected standard. Using correlation type 2, we correlated the model and human judgments of the spectrum level.

#### Co-modulation masking release

The two pairs of standard sources were (1) the input mixture itself and silence and (2) one sound with the noise only and one sound with the tone only. For each noise type and noise bandwidth, the preference measure quantified whether the tone was detected or not as a function of tone level (i.e., whether a model inferred a tone source as separate from the noise). Using correlation type 3, we obtained the Spearman correlation between the model and human thresholds.

#### Frequency modulation

We generated a pair of standards: a sound with only the frequency-modulated components and a sound with only the constant-frequency components. Using correlation type 1, we computed the Pearson correlation between the time-frequency bins of the cochleagrams of the model outputs and the standards, and selected the best match.

#### Mistuned harmonic

The two pairs of standard sources were (1) the input mixture itself and silence and (2) one sound with the harmonic components only and one sound with the mistuned tone only. For each fundamental frequency and harmonic index, the preference measure quantified whether the tone was detected or not as a function of mistuning. Using correlation type 3, we obtained the Spearman correlation between the model and median human thresholds.

#### Asynchronous onsets

The two standards were the ‘basic’ vowel and the ‘shifted’ vowel with the same nominal first formant frequency as the input stimulus. We selected the model output which minimized the distance to either standard (finding the model output that best captured the vowel). Then, we estimated the first and second formant frequencies from the selected output using linear predictive coding (165, 166). The rest of the analysis followed the same steps as for the model with enumerative inference in order to derive vowel thresholds. Using correlation type 3, we computed the Spearman correlation between the model and human vowel thresholds.

#### Cancelled harmonics

The logic for the analysis of this experiment is based on the idea that if the model successfully separated out a whistle source, then the hypothetical whistle source should have a higher amplitude at one frequency than the rest. We computed the spectrum of each model output and found the peaks in the spectrum that were spaced at least 95% of the fundamental frequency apart. We then computed the amplitude ratio of the highest peak to the second highest peak. We selected the model output with the highest peak-to-peak amplitude ratio. The model’s judgment was selected to be the frequency of the highest peak in this output. We then calculated the proportion of trials for which the percent error was less than or equal to two percent, as a function of harmonic number. Using correlation type 2, we correlated this proportion with the human results. This method is conceptually similar to using standards with pure tones of varying frequencies, but it allowed us to obtain a more precise pitch judgment.

#### Frequency proximity

For the pure-tone stimulus, the pair of standard sources was the low-frequency bouncing melody and the high-frequency bouncing melody. For the alternating pure-complex tone stimulus, the pair of standard sources was the ascending melody and the descending melody. Using correlation type 1, we computed the Pearson correlation between the time-frequency bins of the cochleagrams of the model outputs and the standards, and selected the best match.

#### Bistability

For each tone sequence stimulus, we generated two pairs of standards, for the two-source and one-source explanation. The standard set for the two-source explanation contained one sound with all the high-frequency tones and another sound with all the low frequency tones. The standard set for the one-source explanation contained the mixture itself and silence. The preference measure quantified whether two sources were preferred over one source for each setting of Δ*f* and Δ*t*. To compare with human data, we labeled each (Δ*f*, Δ*t*) point as above the human one source threshold (1), in the bistable region (0.5), or below the human two-source threshold (0). Using correlation type 2, we computed the Pearson correlation between the model preferences and the human judgments as reflected in these labels.

#### Buildup of streaming

We computed the model preferences as for bistability but correlated them with the human proportion of two-stream responses.

#### Effects of context (1 and 2)

Both of these experiments included comparing hypotheses, for which multiple sequences corresponded to a hypothesis. Therefore, for each hypothesis, we generated sets of standards corresponding to the sets of structural hypotheses described previously (Methods: Classic auditory scene analysis phenomena, Effects of context). Since multiple sets of standards corresponded to a single hypothesis, we took the minimum distance across all sets of standards to calculate the distance of the model outputs to a hypothesis. The preference measure quantified which hypothesis was preferred (Expt. 1: A and B paired in their own source versus a non-AB pairing; Expt. 2: target tones in their own source versus grouped with distractors). Using correlation type 2, we computed the Pearson correlation between the model preferences and the human judgments.

#### Statistics (Figure 8A and D)

To generate error bars and assess statistical significance for the human-model dissimilarity analysis, we used bootstrap. For each experiment, we bootstrap-resampled the datapoints to obtain uncertainty over the dissimilarity between human and model results (yielding one dissimilarity measure for each experiment and each model, for each bootstrap sample). For each bootstrap sample (n=1000), we averaged the human-model dissimilarity for a model across all experiments. We use the standard deviation of the bootstrap distribution of this average dissimilarity to approximate the standard error (plotted as error bars in Figure 8A and D).

To assess whether the human-model dissimilarity for our model was statistically significantly different from each of the source separation neural networks, we computed the bootstrap distribution of the difference in dissimilarity between the models being compared. We then calculated 99% two-sided confidence intervals for this difference using the “empirical„ bootstrap method (also called the “basic„ bootstrap method in (167)). We used the same procedure to assess whether the model lesions produced statistically significant worse human-model dissimilarity.

## Model lesions

We computed the human-model dissimilarity for our full generative model by simulating psychophysical experiments as described above in Methods section *Classic auditory scene analysis illusions* (blue bar in Figure 8A and D). We assessed the four model lesions in the same way. We created the lesioned models as follows: (1) For the Fixed sources lesion, we set each set of source parameters to the modes of the temporal source priors and each of the variance and lengthscale source priors, specific to that source’s sound type. (2) For the Uniform lesion, we set the distributions over variance and lengthscale to a uniform distribution for each Gaussian Process (frequency trajectory, amplitude trajectory, and spectral shape). (3) For the spectral swap lesion, we used an Ornstein-Uhlenbeck kernel for the harmonic source model and a squared exponential kernel for the noise source model, and then remeasured the source priors as described previously (Methods: Source priors). We only tested ASA results which had a noise or harmonic source in them because the tone sequences are not affected by this lesion. (4) For the stationary covariance lesion, we only altered the whistle source in order to investigate the effect of the non-stationary kernel on tone sequence grouping. We fixed the non-stationary kernel parameter β to zero and re-measured the source priors as described previously (Methods: Source priors).

## Source-separation neural networks

We selected a set of source-separation neural networks to compare to our model. Selection was based on 1) public availability of pre-trained weights, 2) good performance in machine hearing competitions and 3) the goal of spanning a variety of training methods, tasks, network architectures and natural sound datasets. These criteria yielded a set of seven networks:

- ConvTasNet for two-speaker separation, trained on LibriMix (90, 92)
- the same ConvTasNet trained with background noise
- TDCN++ for open domain sound separation, trained on FUSS (91)
- Open-Unmix RNN for music separation, trained on MUSDB18 (93)
- Open-Unmix RNN for music separation, trained on a much larger, private dataset (168)
- Open-Unmix RNN for speech enhancement, trained on the Voicebank+DEMAND corpus (94)
- MIXIT network, trained on YFCC100m (95) (MIXIT is the only network optimized with an unsupervised training objective)

The dissimilarity results for these networks are shown as the striped grey bars in Figure 8D.

The enumerative inference method of obtaining results is arguably most similar to what human participants do when they complete experiments, but was only possible with our model, as it leverages the generative model to constrain inference (blue bar in Figure 8A and D). Therefore, as an additional comparison, for every illusion, we used sequential inference to obtain the most likely scene description (sequential inference only, pink bar in Figure 8D). We analyzed the sounds rendered from this description exactly as for the sounds output by the source separation networks. This is arguably the fairest comparison to the source separation networks, but would be expected to produce a worse match to human results. We ran only 1 seed of sequential inference for each phenomenon.

We also trained an additional source separation network on samples from the generative model. We chose to use the TCDN++ network because it was designed for open domain sound separation with more than two premixture sounds, which is most similar to our model. We used the same network architecture as the TCDN++ reported in (91), training sixteen networks with different hyperparameters (batch size, learning rate) and datasets (higher density of events than the prior or the same density, and the use of domain randomization in which samples were obtained from uniform distributions instead of the model’s prior). From these several TCDN++ networks, we selected the one with the highest similarity to human results. This network was trained on a dataset consisting of approximately 440-h of sound mixtures (n=677170) from the model prior with a higher density of events, limited to contain 1-4 sources. The networks were trained until the validation error converged (for the selected network: 264136 iterations, batch size=20, learning rate=1e-5). This result is shown as the solid grey bar in Figure 8D.

## Everyday sound experiments

Experiments were run online using Amazon Mechanical Turk. Prior to each experiment, potential participants gave consent and indicated that they were wearing earphones or headphones. They used a calibration sound to set their volume to a comfortable level. Participants were initially screened with a short experiment to check that they were wearing earphones or headphones (169). If participants failed the headphone check, they were compensated and did not continue to the main experiment. All experiments were approved by the Committee on the use of Humans as

Experimental Subjects at the Massachusetts Institute of Technology, and were conducted with the informed consent of the participants.

### Stimuli

To assess the ability of the model to infer perceptually valid scenes from naturalistic sounds, we sourced a small subset of sounds from the Free Universal Sound Separation dataset (FUSS) (91). The FUSS dataset contains mixtures generated by adding together audio clips of everyday sounds and then simulating reverberation. FUSS was designed for open domain source separation, with each premixture clip derived from one of over 300 sound categories.

The mixture clips in FUSS are 10-s long, composed of 1-4 sounds from the FSD50K dataset with simulated reverberation. There is always one background sound in each mixture clip, defined to be a sound which extends the entire duration of the clip. We randomly selected 50 2-s clips from the training set of FUSS, subject to a few constraints. The main constraint was that each mixture clip should contain three pre-mixture recordings of at least 200-ms in duration. Second, although sounds in FUSS are not explicitly labeled, we recovered labels from FSD50K for each pre-mixture sound. This allowed us to exclude four categories out of the 357 categories included in FUSS: “Speech”, “Scratching (performance technique)”, “Mechanisms” and “Human group actions”. We excluded speech and scratching because we knew that our model would be poorly suited to the variable spectra that occurs in these sounds. We excluded Mechanisms and Human group actions because their names suggested that they contained more than one perceptual stream. We used sequential inference to obtain full scene descriptions for each sound mixture, and then rendered each inferred source sound into audio using the maximum a posteriori scene description. We used these rendered audio signals in the experiments. In addition, the original mixture clips were used in Experiment 1, and their corresponding pre-mixture clips were used in Experiments 1 and 2.

For both experiments, the maximum level across experimental stimuli was set to +5 dB and -4 dB relative to the calibration level for Experiments 1 and 2 respectively. We maintained the relative levels of the recorded audio and the inferred sources. We excluded 7 model sounds out of 144 because they did not reach a threshold sound level and would have been inaudible over typical headphones. Pilot participants reported that a few sounds remained which were difficult to hear (presumably low-frequency sounds).

### Experiment 1 procedure

For the main experiment, we randomly split the 50 mixture sounds into two halves. For each participant, one of these splits was randomly assigned to the model condition, and the other was assigned to the recorded audio condition. On each trial, participants heard a mixture sound followed by two additional sounds that they had to choose between. Participants were instructed to select which of the two sounds was part of the initial mixture. On recorded sound trials, the correct sound was a pre-mixture sound from the mixture, and the incorrect sound was a pre mixture sound from a different mixture randomly chosen from FUSS. We selected the incorrect option to not share a class label with any of the premixture sounds in the mixture for that trial. On model sound trials, the correct sound was a source inferred by the model from the mixture. The incorrect sound was a randomly selected source inferred by the model from a mixture in the other split of mixture sounds. The incorrect sounds were randomly chosen for each participant. Participants were told that the correct answer could either be the exact sound from the mixture or a computer imitation of a sound present in the mixture. Participants could listen to the sounds multiple times and did not receive feedback.

### Experiment 1 participants

115 participants were recruited through Amazon Mechanical Turk. The main experiment included 10 catch trials, which were not included in the main analysis. The catch trials comprised an independent set of recorded sound trials and were the same across all participants. 35 participants failed the headphone check and 16 participants answered less than 7 out of 10 catch trials correctly. The data from these 51 participants was removed. We analyzed the data of the remaining participants (N=64, 34 male, 29 female, 1 non-binary or other based on self-report, mean age = 43.7 years, S.D. = 11.6 years).

### Experiment 1 sample size

We determined sample sizes a priori based on a pilot study with 40 participants. We computed the split-half reliability of the participant’s accuracy on individual trials, varying the size of the splits and extrapolating to estimate the sample size needed to obtain a split-half reliability of 0.9, yielding a target sample size of 60 participants.

### Experiment 1 analysis/statistics

For Figure 9B, we computed the percent correct for each participant in each condition (recorded, model) by taking the proportion over trials. The figure reports the average and standard error over all participants (regardless of the half of the data they were assigned to). We also report the two tailed 95% confidence interval over all participants for each condition, using the function `DescrStatsW.tconfint_mean` from the Python statsmodels module version 0.13.2. For Figure 9C, we computed the percent correct for each model inference by taking the proportion of correct answers across participants who completed a trial with that sound.

### Experiment 2 procedure

Upon successful completion of the headphone check, participants proceeded to the experiment instructions and a set of six practice trials. Participants received feedback on the first four practice trials. The last two practice trials did not have feedback. If participants did not correctly answer the last two practice trials, they were compensated and did not proceed to the main experiment.

Each trial presented two sets of sounds: source sounds inferred by the model (“row sounds”) and pre-mixture sounds (“column sounds”), which were arranged in the headers of a grid. All sounds on a trial corresponded to a single mixture. The number of row sounds and column sounds varied from mixture to mixture depending on the number of premixture sounds and inferred sources that were recognizable to human listeners (the number of inferred sources also varied from mixture to mixture). Participants were instructed to mark the corresponding checkbox within the grid if any part of a row sound matched part of a column sound. Participants were told that the purpose of the task was to evaluate a computer sound-synthesis algorithm. They were warned that the computer-generated sounds were meant to imitate the column sounds but that they might not always be perfect resemblances. They were explicitly told that multiple row sounds might match to the same column sound or vice versa, or that a column sound might not have any matches (the practice trials also demonstrated these possibilities).

For each mixture, we excluded model sounds and premixture sounds for which the average performance of participants in Experiment 1 did not exceed 60%. The premixture sounds that were unrecognizable tended to be those that were completely masked in the mixture. The model sounds that were unrecognizable had already been measured in Experiment 1, and we excluded them to instead measure other types of deviations in perceptual organization. These criteria resulted in the exclusion of 20 model sounds (out of 124) and 8 premixture sounds (out of 140).

Because the experiment only presented inferred sources that were recognizable in the original mixture, it was reasonable to assume that each inferred source should correspond to at least one pre-mixture sound. Participants were thus not allowed to proceed to the next trial until they placed at least one checkmark for each row sound.

Participants performed 50 trials (plus ten catch trials – see below), one for each of the included mixtures. Participants were allowed to listen to the sounds as many times as they wanted and in any order.

### Experiment 2 sample size

Based on a pilot experiment with 7 participants, we found that the over-combination deviations were the most rare and so this proportion required the largest sample size to estimate with sufficient precision (relative to the small effect size). For a range of sample sizes, we calculated the 95% confidence interval on the mean by bootstrap with the pilot data. We determined the sample size required to obtain a 95% confidence interval with width = 0.05, which yielded a target sample size of 21.

### Experiment 2 participants

An initial group of 49 participants were recruited through Amazon Mechanical Turk. The main experiment included 10 catch trials which were not included in the main analysis. The catch trials had the same three audio recordings for the row and column sounds, and the correct answer was to match a sound with only itself. 22 participants failed the headphone check and 2 participants answered less than 7 of the 10 catch trials correctly. We analyzed the data of the remaining participants (N=25, 14 male, 9 female, 2 non-binary or other based on self-report, mean age = 41.8 years, S.D. = 11.6 years).

### Experiment 2 analysis/statistics

We estimated the overall reliability of participant responses using the intra-class correlation, considering each target to be an inferred source/premixture sound pair. We used the function `intraclass_corr` from the Python pingouin module version 0.5.1.

Of the four deviations analyzed in the everyday sounds experiments, Experiment 2 was meant to measure absence, oversegmentation, and overcombination deviations. We tallied the total number of these three deviations per premixture sound for each participant. This allowed us to compute the five quantities displayed in Figure 9G. Bars 1-3, labeled ‘Absent’, ‘Oversegment’, and ‘Overcombine’, were computed as the proportion of premixture sounds with each of the corresponding deviation type, averaged across participants. Bar 4, labeled ‘No deviations’, was computed by finding the proportion of premixture sounds for which a participant checked only one model inference and then averaging that proportion across participants. Bar 5, labeled ‘No deviations + Most common’, is based on a subset of the proportion indicated by Bar 4. We determined which response (i.e. pattern of checkmarks across model inferences) was most common for each premixture sound. This allowed us to compute the average proportion of premixture sounds for which a participant checked only one model inference and for which that concurred with the most common response. We plotted the standard error across participants.

To derive the chance level for each of these average proportions (except overcombine deviations - see below), we randomly permuted our data 5000 times within each worker’s responses, subject to the constraint that each row contains at least one checkmark. We then computed the same statistics on the permuted datasets. We plotted the average proportion of premixture sounds across permutations and workers as well as the standard error across workers. We use a permutation test to determine statistical significance: we report the quantile of the empirical average proportions (across workers) with respect to the distribution of 5000 permuted average proportions (across workers). The proportion of overcombine deviations was an exception because by design they occur at the same rate in the permuted and original data. Instead, we computed the two-tailed 95% confidence interval of the proportion of overcombine deviations by bootstrap with 5000 samples.

We estimated the reliability of each proportion via the split-half reliability, using 10 random splits of the participants. For each premixture sound, we computed the proportion of participants responding with each deviation type. Then, for each deviation type and each split, we computed the Pearson correlation (across premixture sounds) of that proportion between each half of participants. We averaged this correlation across the 10 splits. The average split-half correlation for each deviation type is plotted as a bar in Supplementary Figure 4. To obtain the expected chance level of this reliability for each deviation type, we computed the split-half correlation for 100 of the permutations described in the previous paragraph (averaged across 10 random splits for each permutation). We computed the p-value for the actual split-half reliability as the proportion of 100 permutations for which the average permuted split-half correlation exceeded the average split-half correlation on the real data.

## Acknowledgments

This work was supported by NSF Grant No. BCS1921501. We thank Ted Adelson, Dan Ellis, and Josh Tenenbaum for helpful discussions, Alex Lew for advice on inference, and the McDermott Lab for comments on an earlier draft of the manuscript. We also thank the creators of the icons and sound examples in our figures. We provide a full list of creators along with licenses and links to the original work on our project website: https://mcdermottlab.mit.edu/mcusi/bass/attributions.html.

## Supplementary Information

**Supplementary Algorithm 1.**
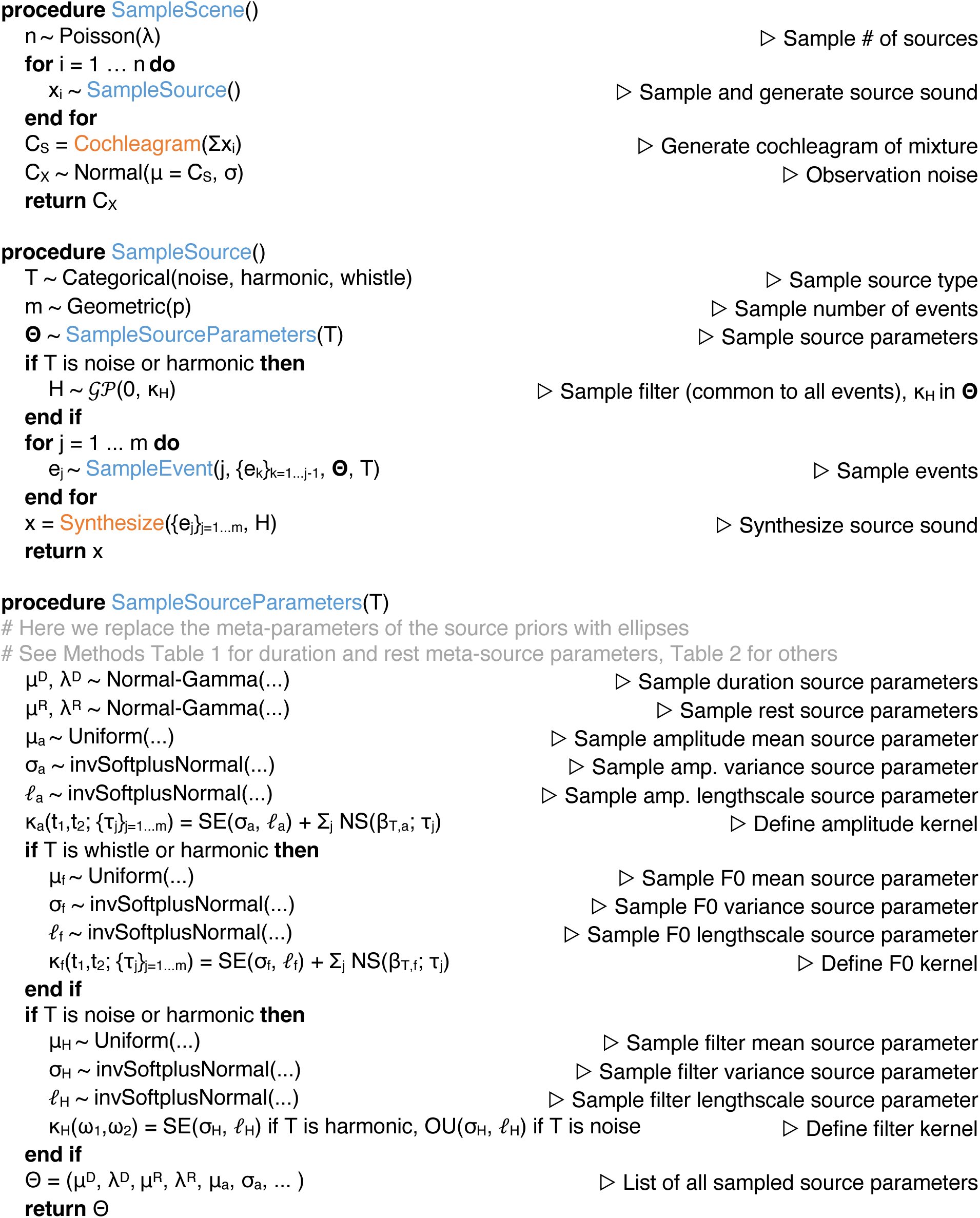
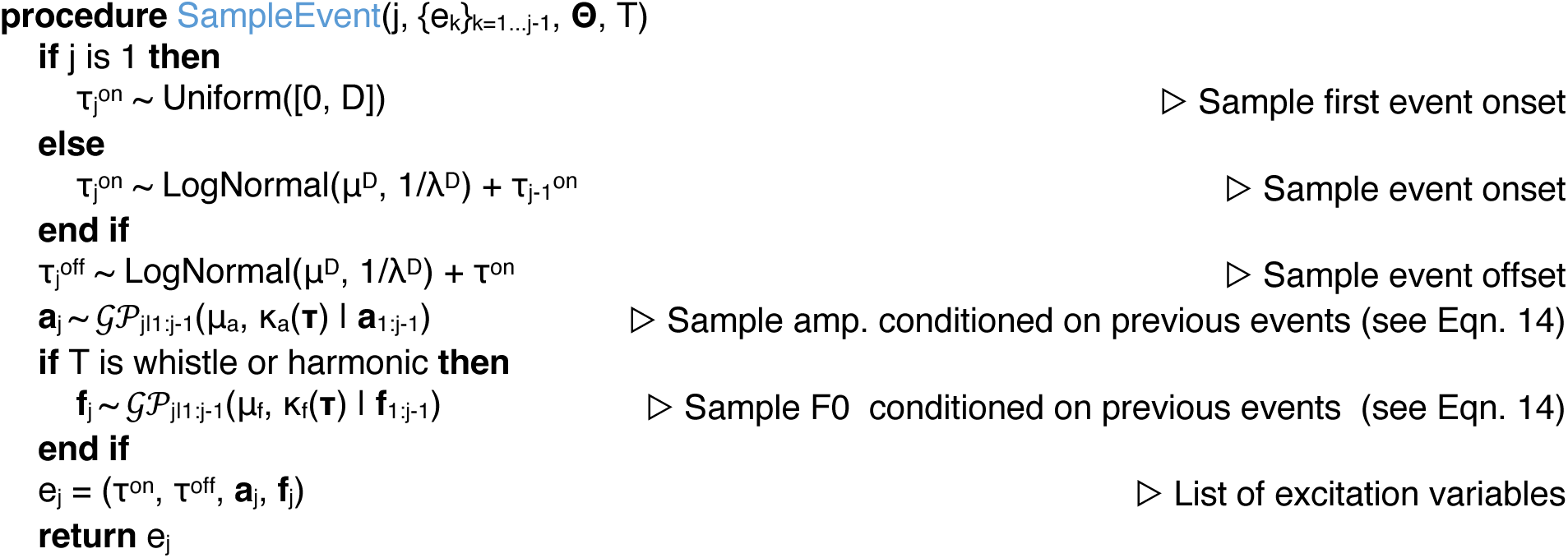
Probabilistic program that defines the generative model. Blue: high-level sampling functions. Orange: rendering functions. Methods Tables 1 and 2 specify meta source parameters.

**Supplementary Figure 1.**
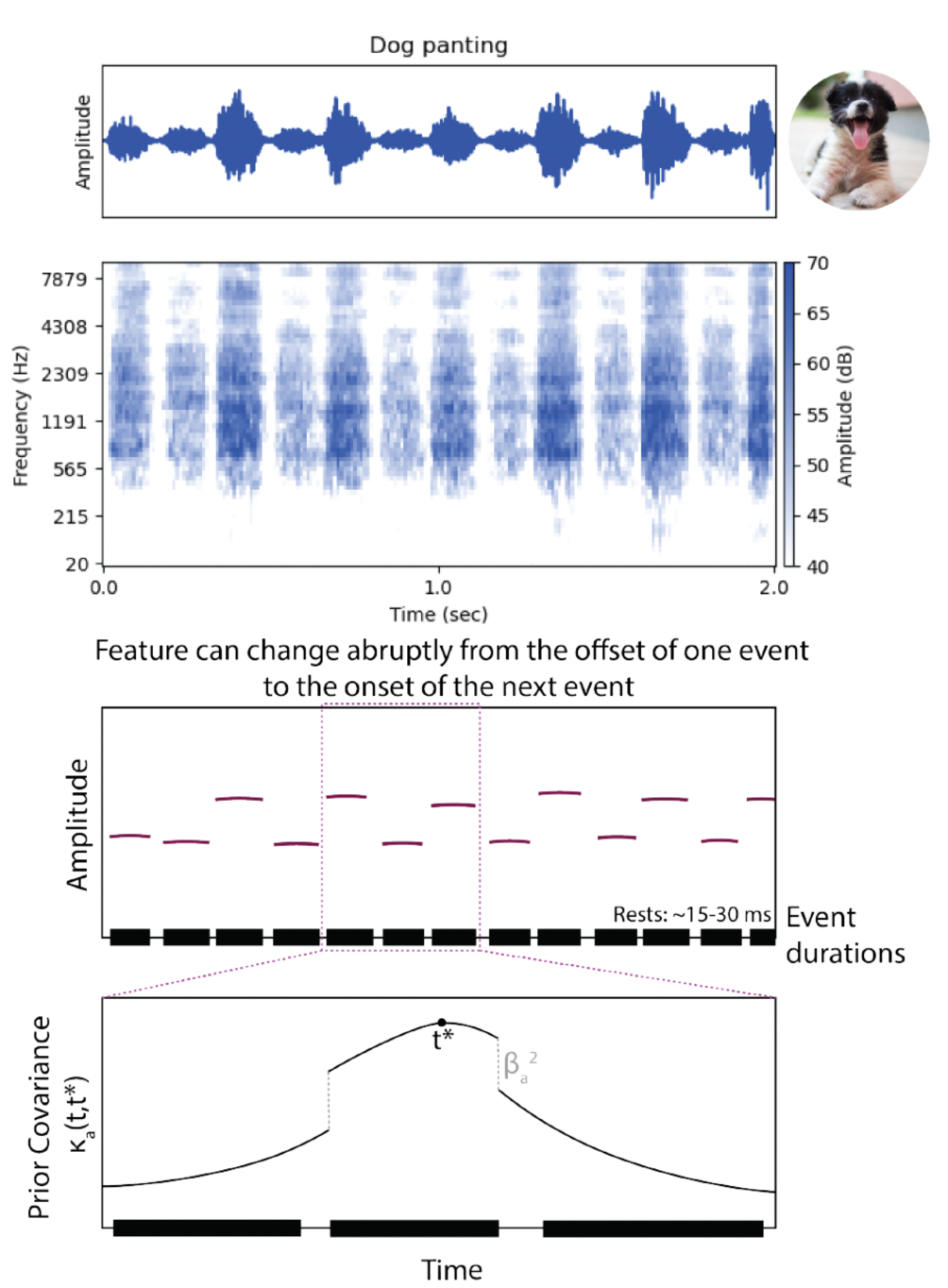
Event structure as implemented by the non-stationary kernel. The generative model can account for a situation where a single source slightly changes its sound generation process between events. Top two panels: sound and cochleagram of a dog panting, alternating between the in- and out-breath. Middle: Schematic latent variable description of the sound. The red lines indicate the amplitude trajectory, y-axis is amplitude level and x-axis is time. The black rectangles at the bottom depict each event duration. Bottom: Non-stationary prior covariance kernel. Black rectangles at the bottom depict event durations. Timepoints are more correlated within an event than between events.

**Supplementary Figure 2.**
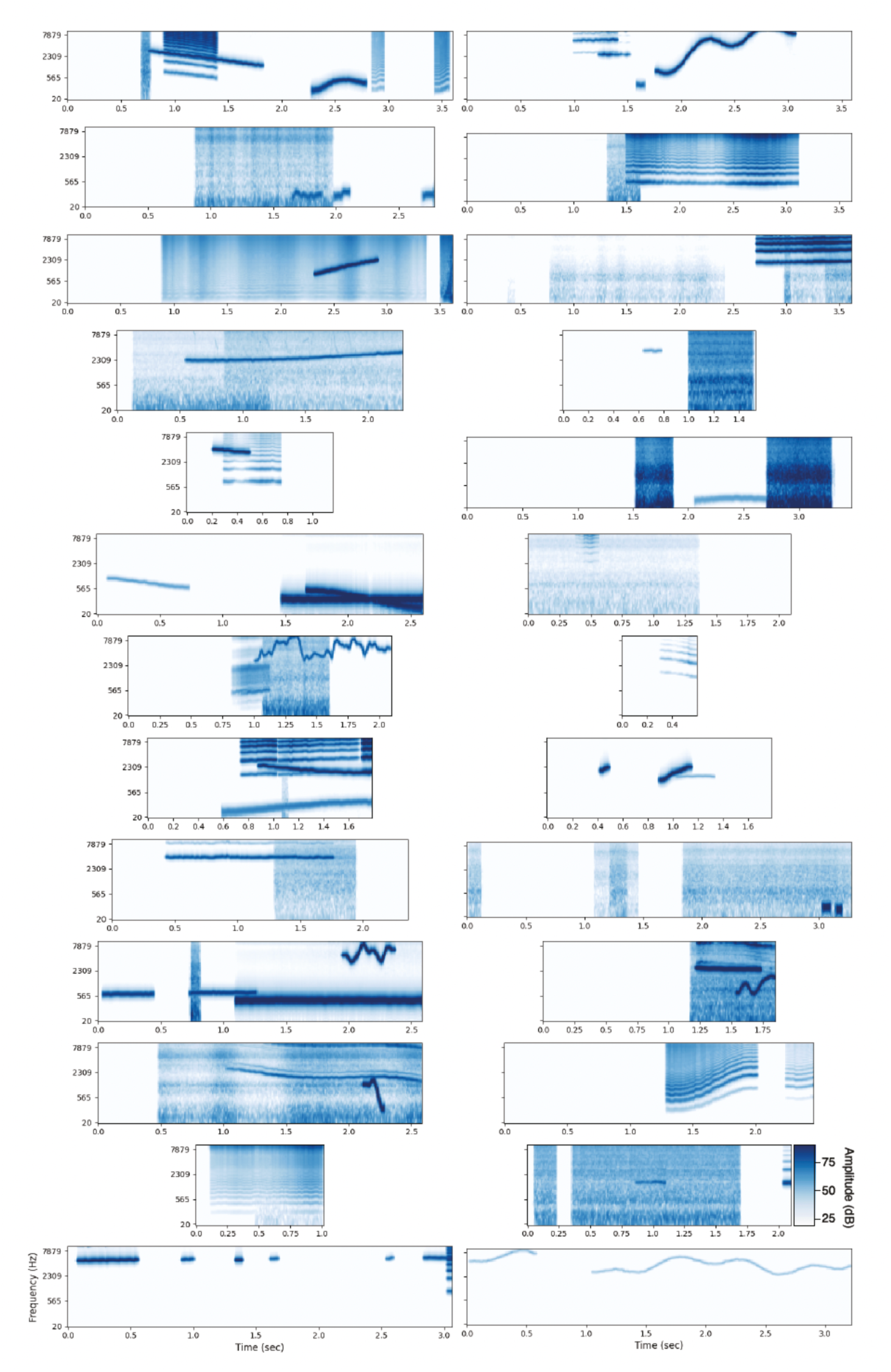
Scenes sampled from prior p(S), rendered as cochleagrams. Scenes sampled from prior show variation in the number of sources and events, event timing, frequency and amplitude modulation, and spectral shape. These sounds demonstrate the model’s expressivity, while also revealing its simplifying assumptions (e.g. time-symmetry, raised cosine ramps for all events, etc.).

**Supplementary Table 1.** Classic auditory scene analysis illusions.

|  | **Citation** | **Result** | **Experiment** |
| --- | --- | --- | --- |
| Masking and perceptual filling in | Continuity illusion (66) | Perceived temporal continuity of sounds through masker depends on masker level | Masking and continuity thresholds |
|  | Homophonic continuity (67) | Abrupt but not gradual amplitude changes cause segregation | Subjective report |
|  | Spectral completion (68) | Analogous perceptual completion in frequency domain | Subjective matching of spectra |
|  | Co-modulation masking release (69) | Detection thresholds of tones decreases as bandwidth of co-modulated noise increases | Tone detection (2AFC) |
| Simultaneous grouping | Frequency modulation (72) | Frequency-modulated harmonics segregate | Subjective report |
|  | Mistuned harmonic (76) | Mistuned harmonic component segregates from complex tone | Subjective report (1AFC) |
|  | Asynchronous onsets (77) | Harmonic component with asynchronous onset segregates | Vowel categorization |
|  | Cancelled harmonics (79) | Amplitude gated harmonic component segregates from complex tone | Pitch matching |
| Sequential grouping | Frequency proximity (81) | Crossing tone sequence segregates into non-overlapping bouncing streams | Subjective report |
|  | Bistability (82) | Increasing frequency difference and decreasing interstimulus interval increases segregation, with a region of bistability | Subjective report |
|  | Buildup of streaming (84) | Segregation increases with repetition | Subjective report |
|  | Effects of context (14, 85) | Context can promote segregation of grouping of two tones | Subjective report |

**Supplementary Table 2.**
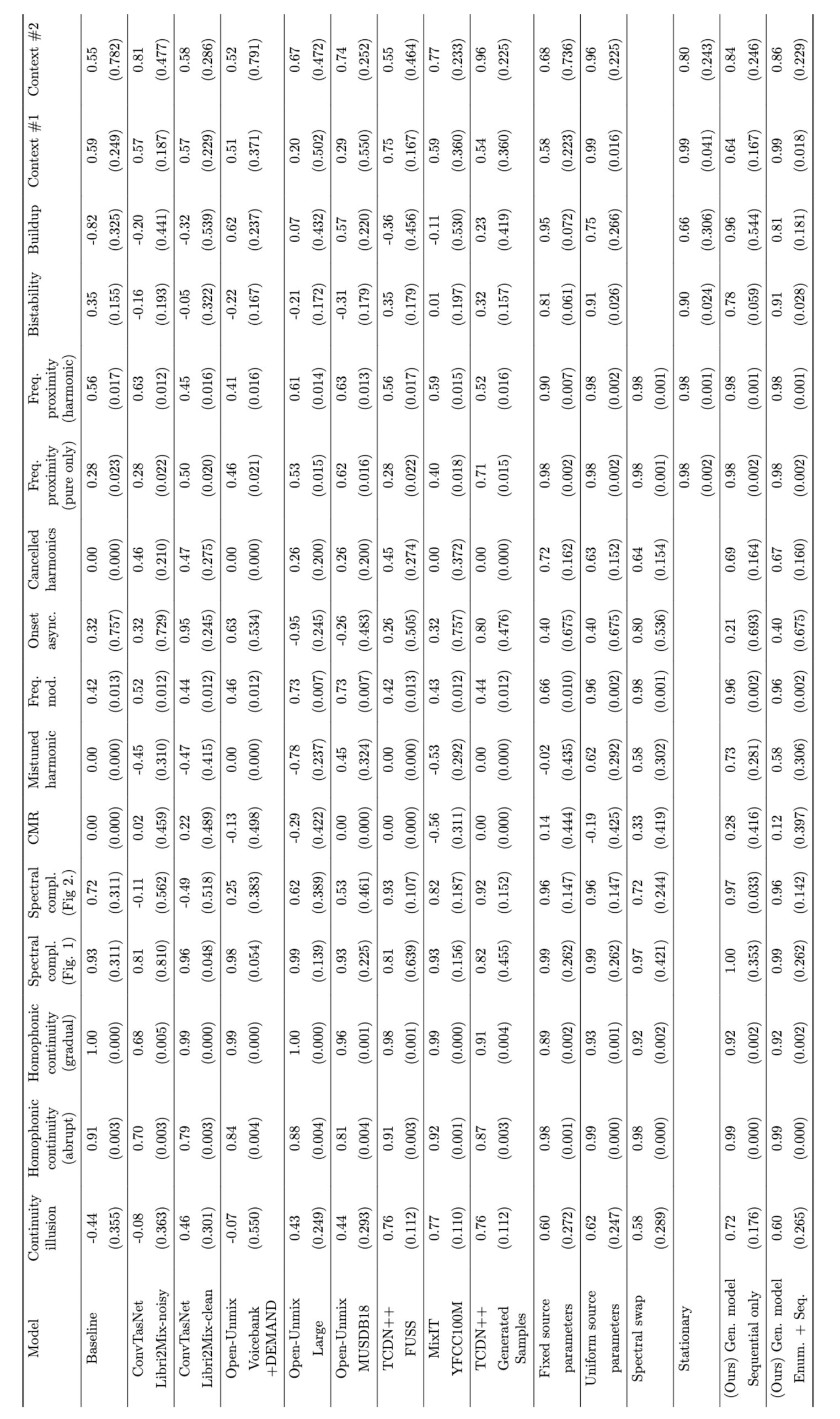
Human-model correlations for individual illusions. Used to compute dissimilarity across models and illusions. Bootstrapped standard errors in parentheses.

**Supplementary Figure 3.**
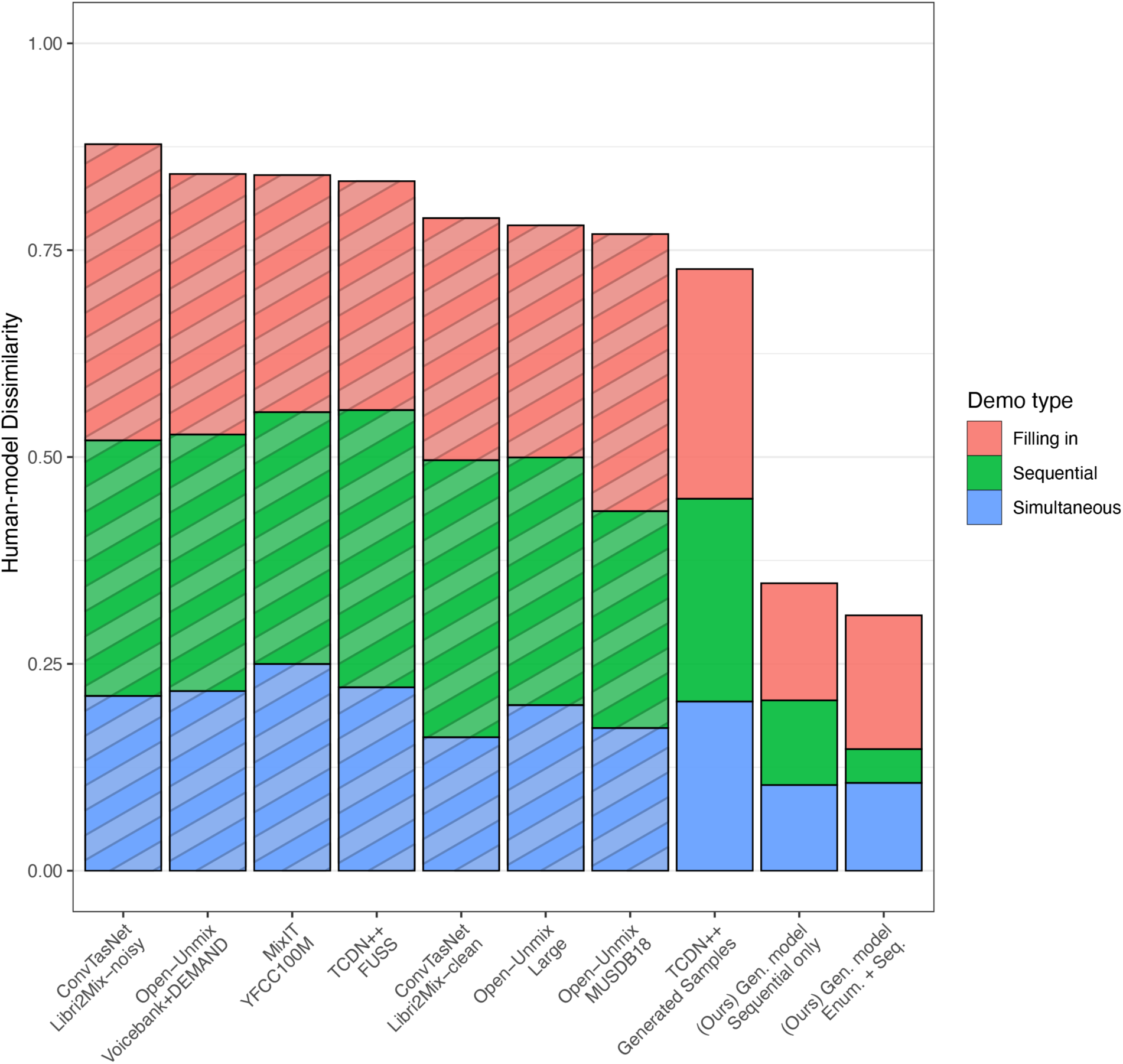
Comparison of model-human dissimilarity for different classes of illusions.

**Supplementary Figure 4.**
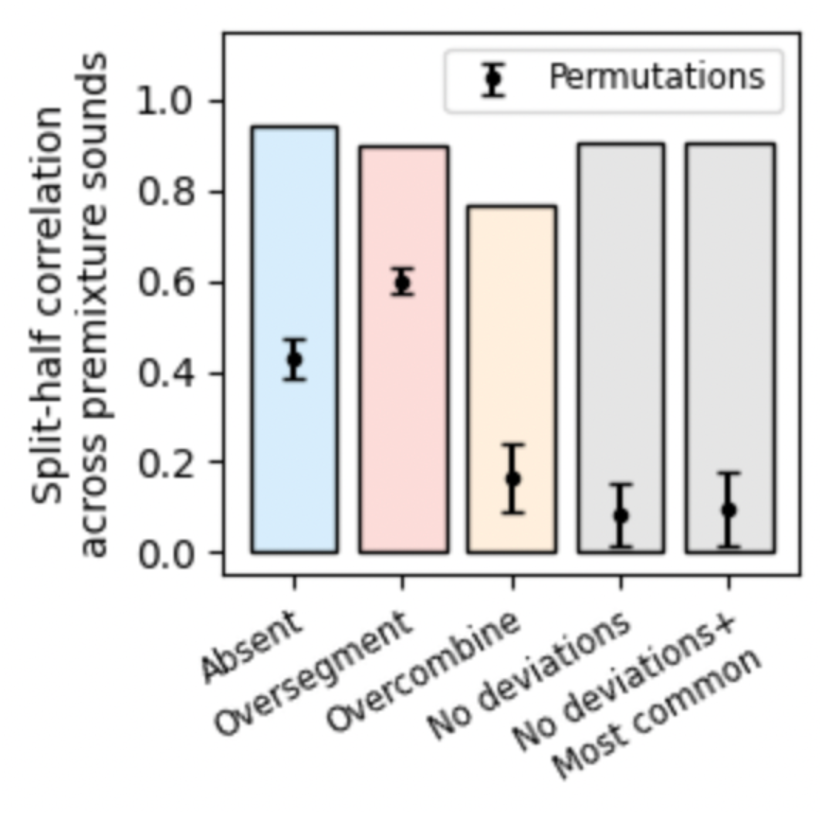
Split-half reliability of the incidence of perceptual organization deviations for pre-mixture sounds. For each deviation type, we computed the Pearson’s correlation across premixture sounds, of the proportion of workers reporting a deviation for the premixture sound, between split-halves of participants. These correlations show that the same mixtures tend to produce the same types of deviations across participants. The error bars plot ±1 standard deviation over permutations of participant responses.

**Supplementary Figure 5.**
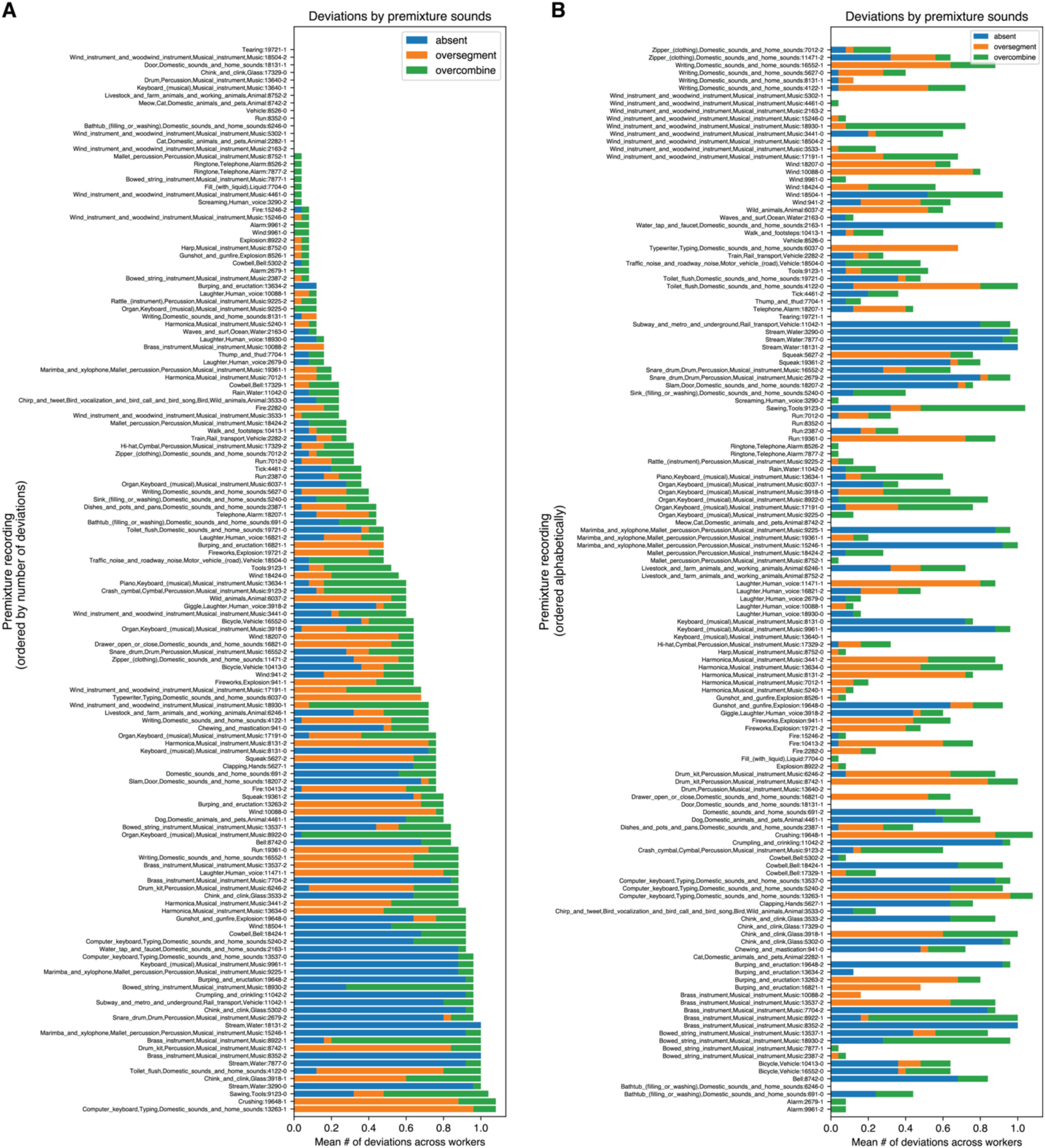
Experiment 2 data per pre-mixture sound (n=21). Zoom on digital copy to see labels. A) Results by premixture clip in Experiment 2. Each bar displays the average number of deviations across participants, tallied for a particular premixture clip. The bars are ordered by total number of deviations, increasing from top to bottom. B) Same data, but organized alphabetically by FUSS category label.

**Supplementary Figure 6.**
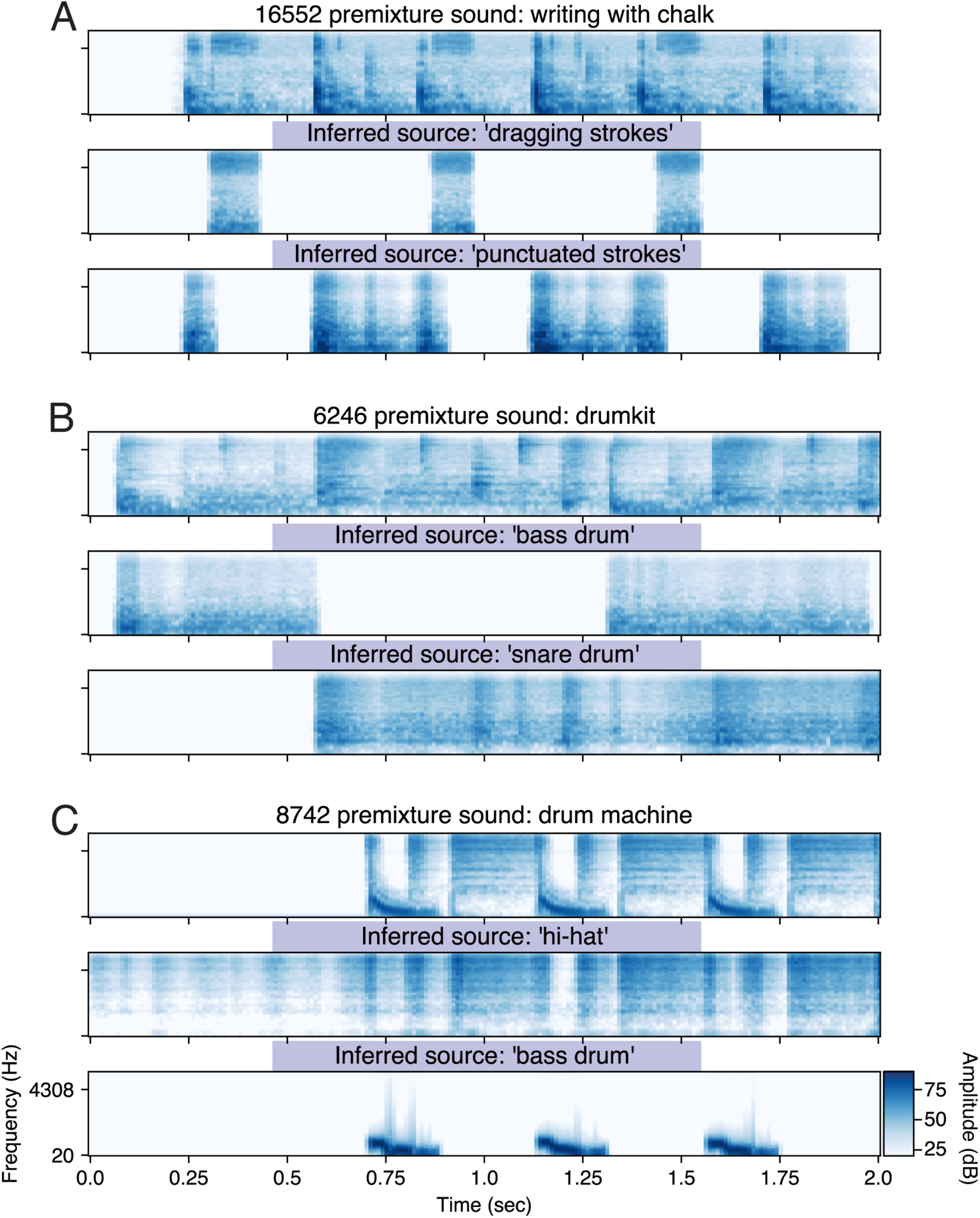
Over-segmentation deviations that occur when various sound generating processes within a single premixture clip have some common causal factor, involving sequences. A) An over-segmentation deviation for a premixture clip of writing on a chalk board, in which dragging the chalk across the board is punctuated by shorter strokes. The sustained contact sounds and punctuated contact sounds are explained as separate sources. B) A similar over-segmentation deviation occurs in a premixture clip of a drumkit, for which the model infers a noise source for the bass drum and a different noise source for the snare. C) Another drum machine in which the hi-hat and bass are inferred separately, is also linked to an over segmentation deviation. We suggest that percepts for these sound examples are akin to hierarchical grouping in vision. Such hierarchy cannot be captured by our generative model, and represent a direction for future research.

**Supplementary Figure 7.**
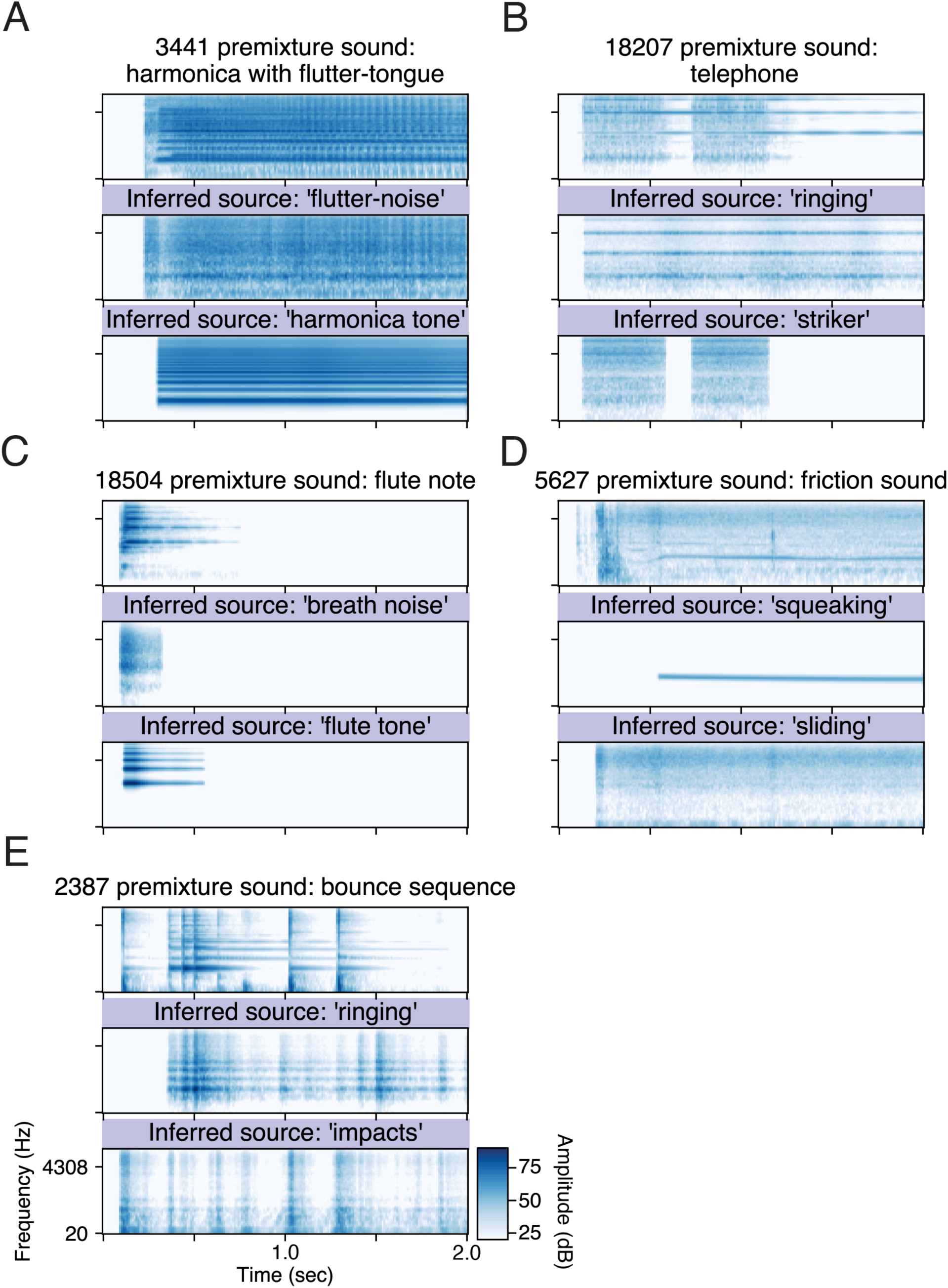
Over-segmentation deviations that occur when distinct excitation processes occur within a single premixture clip. A) Pre-mixture clip: a harmonica played with fluttertongue, in which the player rolls their tongue while playing a pitched note. The model separates this into the noisy sound of the tongue-roll and the pitched sound of the harmonica tone. B) Pre-mixture clip: a telephone ringing. The model separates the resonances of the bell and the noisy sound of the striker. C) Pre-mixture clip: a flute note. The model separates the breath noise from the pitched tone. D) Pre-mixture clip: a sound produced by friction. The model separates the pitched squeak and noisy sliding sound. E) Pre-mixture clip: a series of bounces. On the second to fifth bounce, the object hits a resonant metal surface (you can see the modal resonances in the sound). The modal separates the sequence into ringing sounds and the impact ’thuds’. The event at the end of the ringing source (starting at 1.5-sec) corresponds to a sound in the mixture that is not in the pre-mixture clip (see next Supplementary Figure). We again suggest that hierarchy in the source models may be necessary to explain human perception in these examples.

**Supplementary Figure 8.**
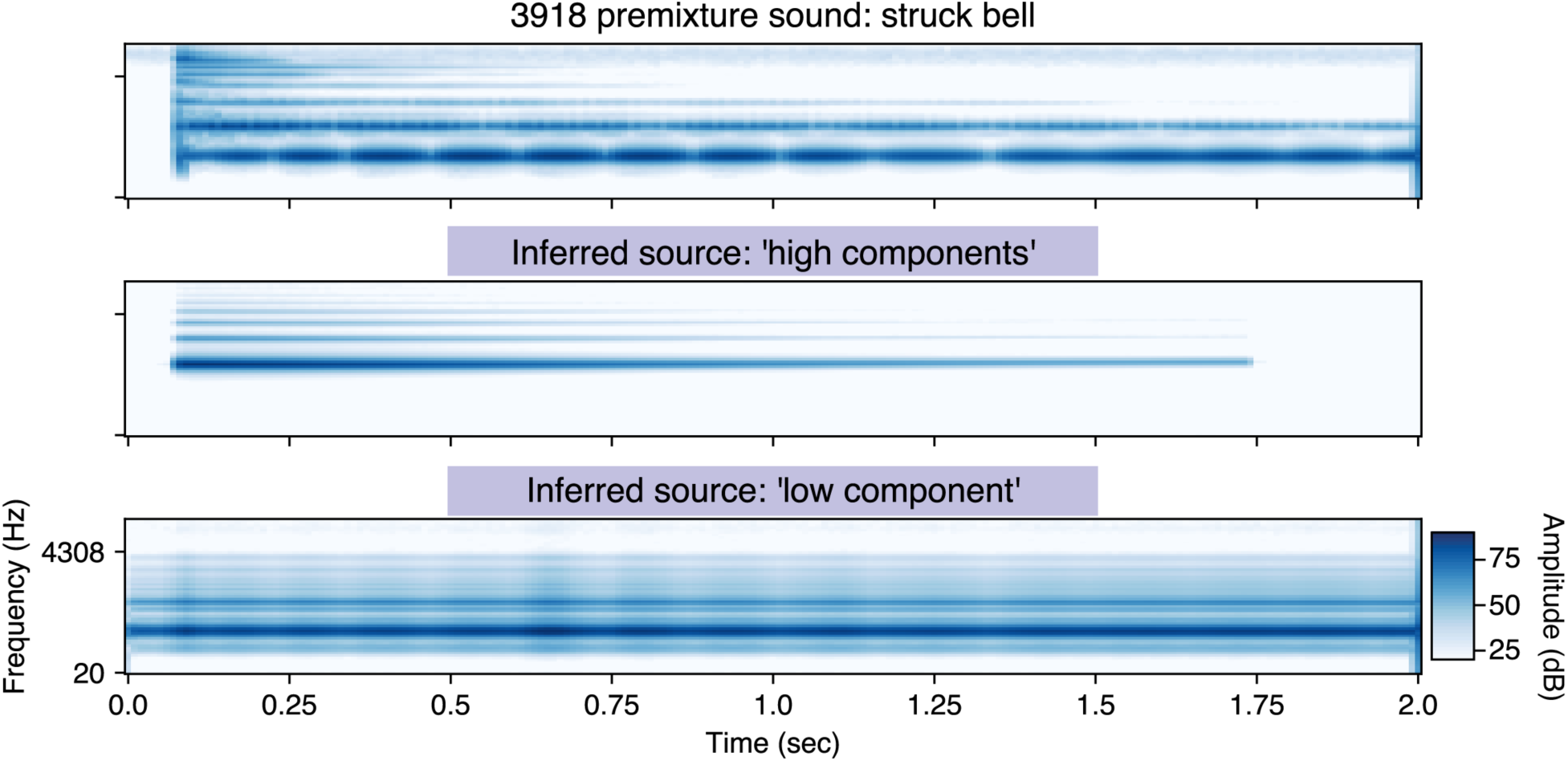
Impact sounds illustrate the need for more nuanced modelling of spectrum and amplitude. In this premixture impact sound of a struck bell, the model over segments higher and lower frequency sound components which decay at different rates. Other issues for the model include the bell sound’s inharmonicity and its transient.

**Supplementary Figure 9.**
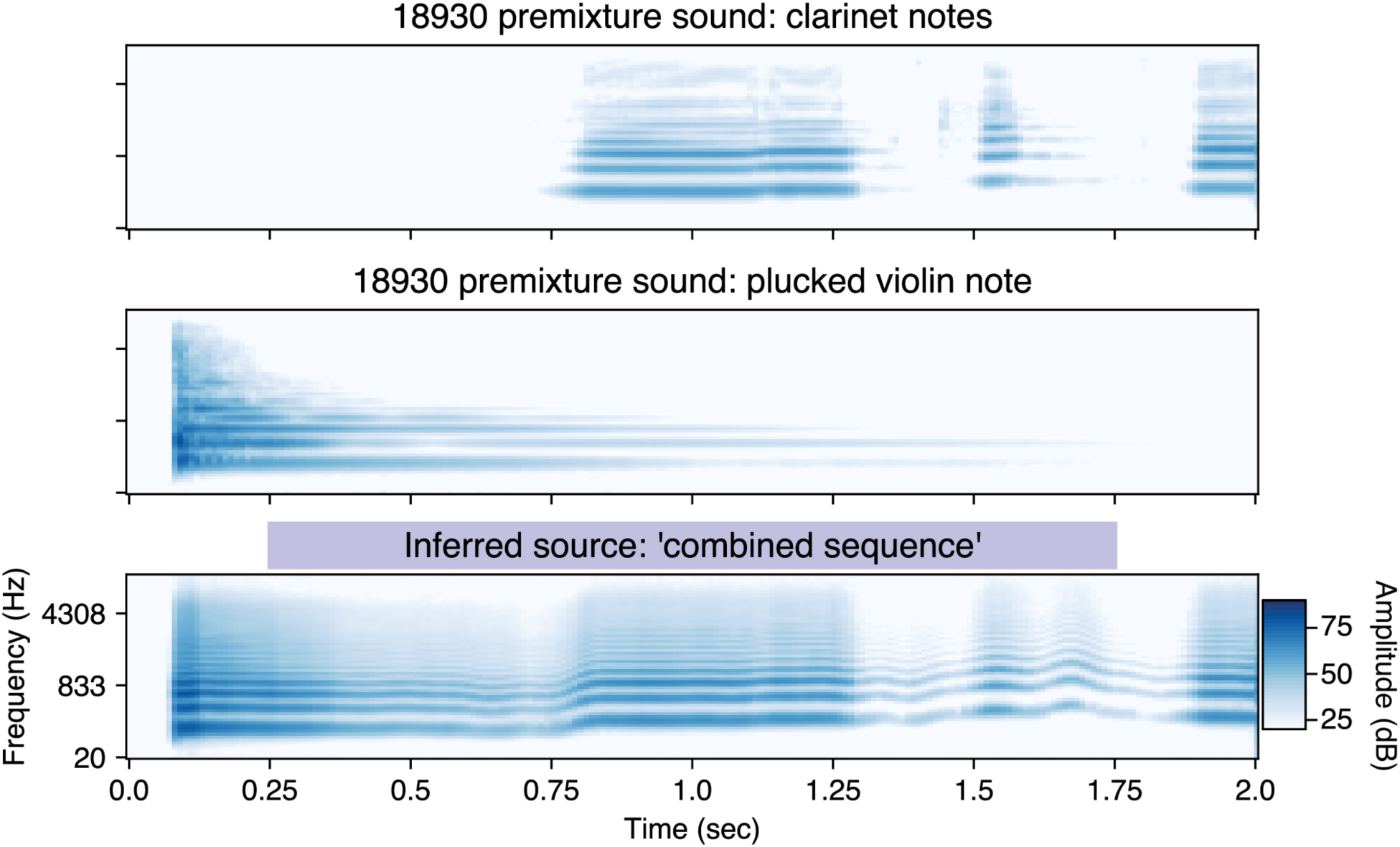
Instrument sounds illustrate the need for more nuanced modelling of spectrum and amplitude. An over-combination deviation occurs when the model sequentially groups a plucked violin note with the subsequent clarinet notes. Since both sources have similar fundamental frequencies, spectral envelopes, and amplitudes, the model cannot separate these perceptually distinct sources.

**Supplementary Figure 10.**
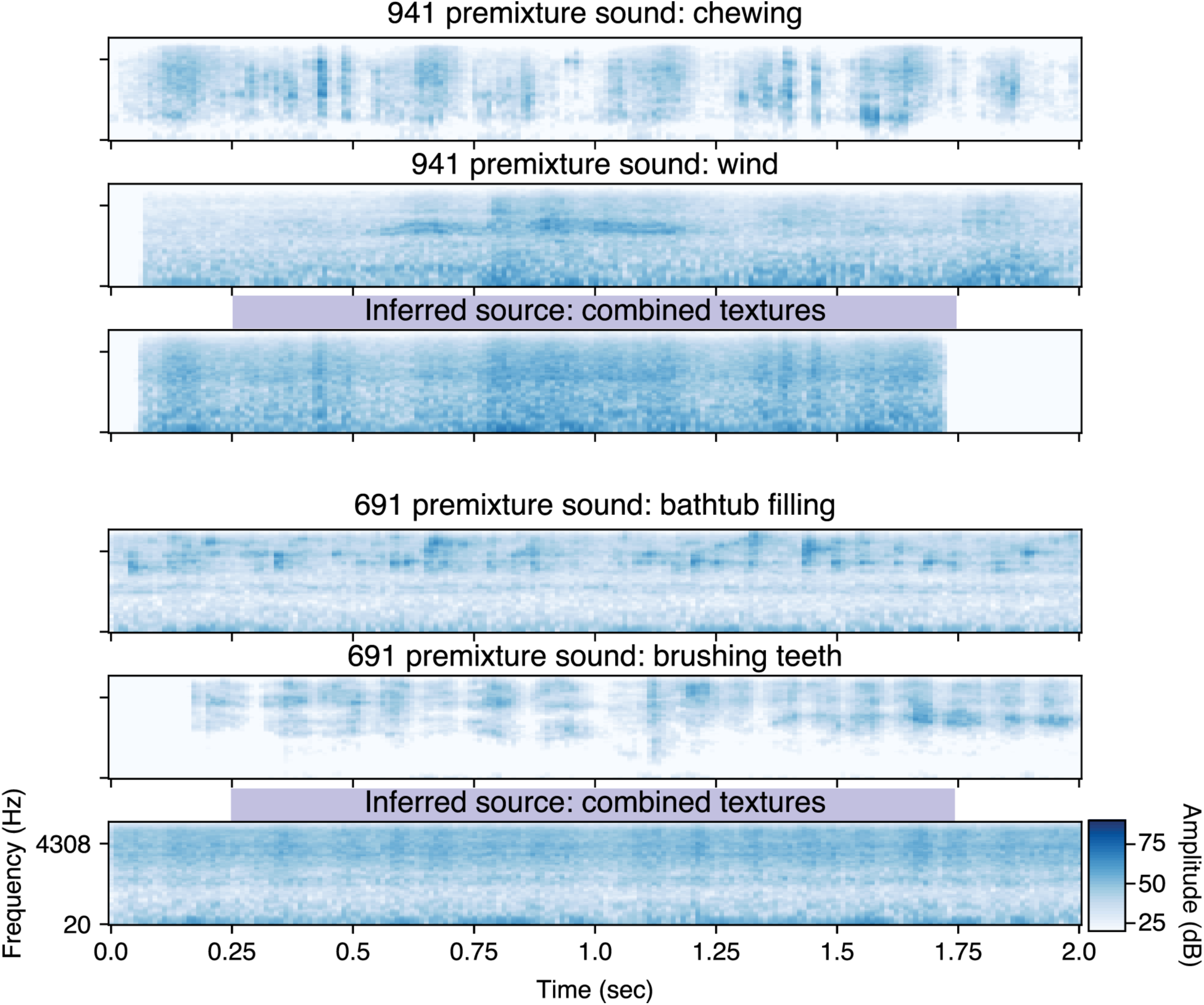
Sound textures illustrate limitations of our generative model (1). In these two examples, two premixture sound textures are explained with a single noisy source. These examples are associated two kinds of deviations, over-combination and absence.

**Supplementary Figure 11.**
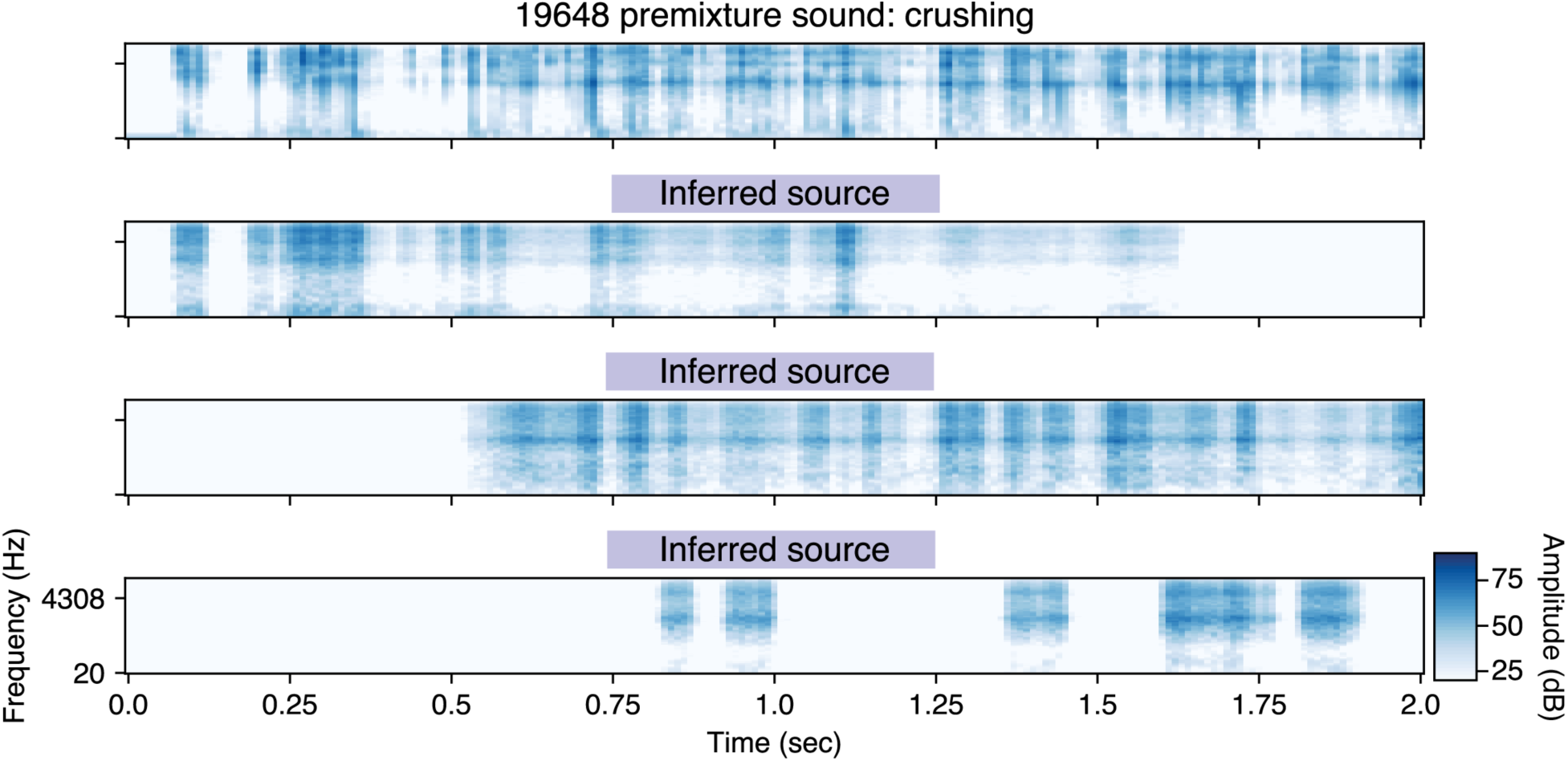
Sound textures illustrate limitations of our generative model(2). Multiple inferred sources are used to explain this complex sound texture. This premixture sound is associated with an over-segmentation deviation.

**Supplementary Figure 12.**
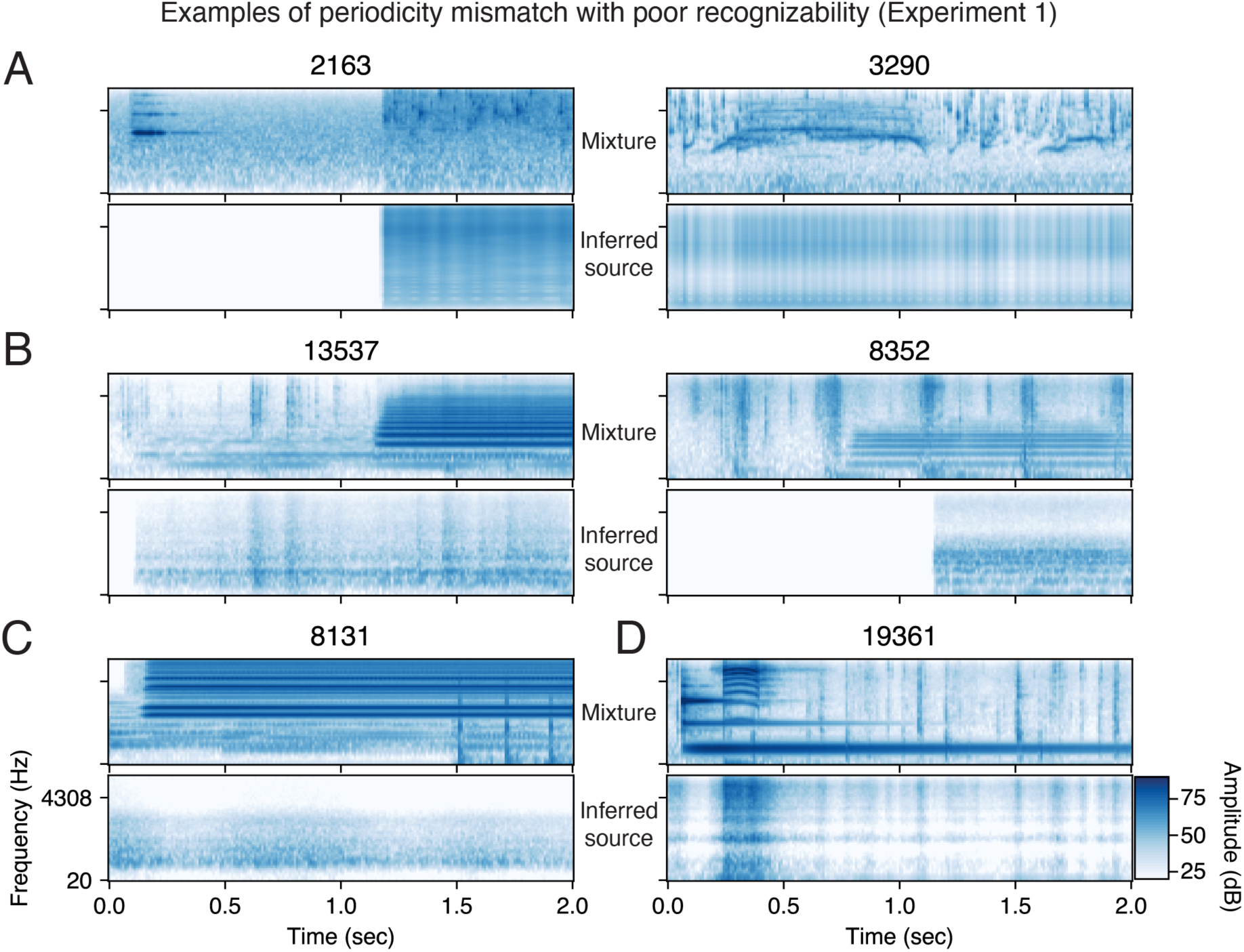
Examples of source inferences with poor recognizability in Experiment 1, due to periodicity mismatch in the mixture and inferred source. Title numbers refer to FUSS filename. A) Noisy sounds are explained as harmonics. 2163 contains rain pattering during the duration of the inferred harmonic source. 3290 contains water gurgling during the duration of the inferred harmonic source. B) A note on a low-frequency instrument is explained as noises rather than as a periodic sound. 13537 contains a single note on the cello being bowed and 8352 contains a trumpet note. C) Low-frequency instrument sounds with overlapping notes are explained as noises rather than periodic sounds. 8131 contains overlapping piano tones. D) High-frequency harmonic squeak is explained as a noise in mixture 19361.

**Supplementary Figure 13.**
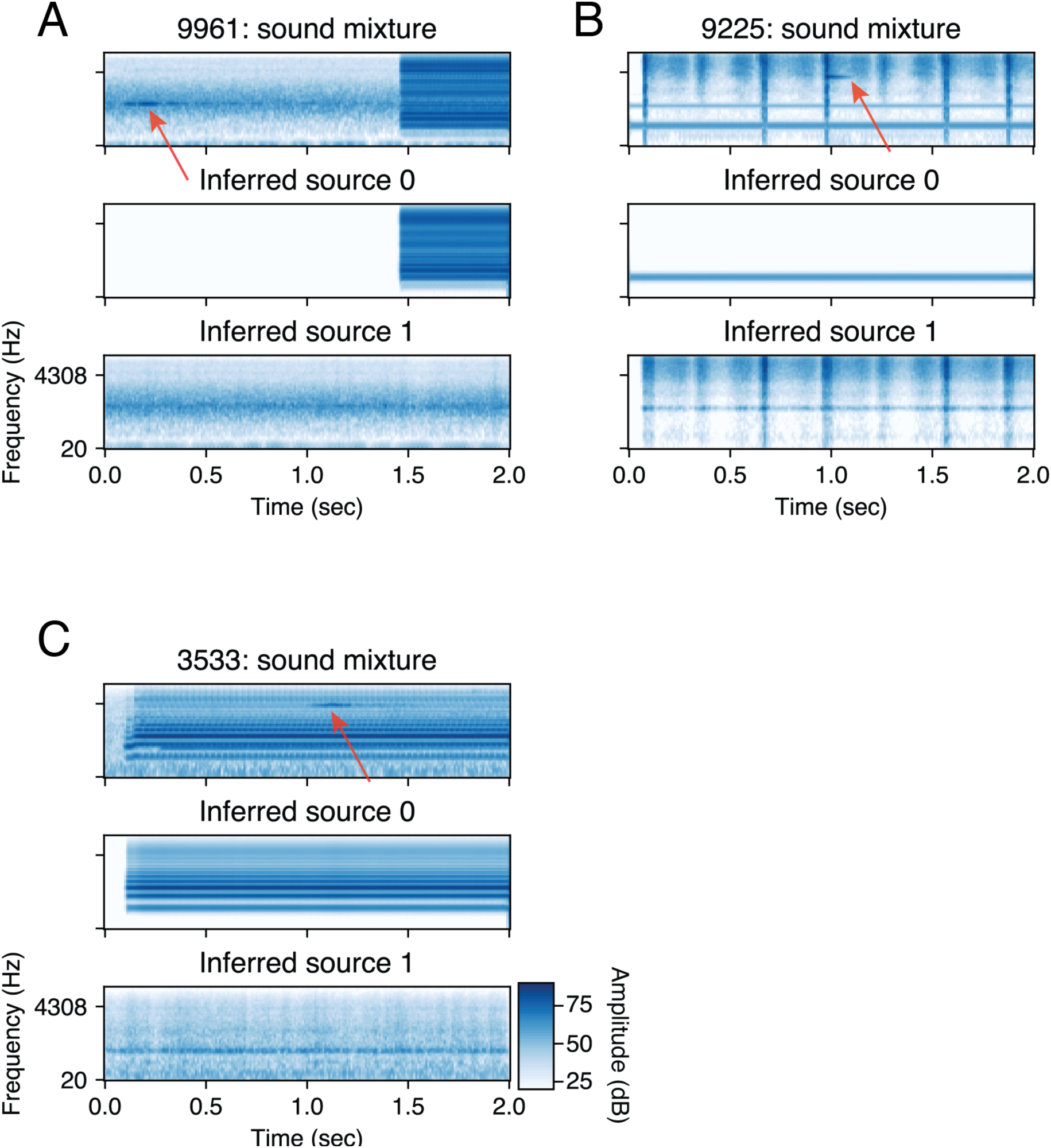
Examples of inferences in scenes containing a quiet tone in noise. Red arrows indicate where the short, quiet tones occur in the sound mixtures. We suggest that the lack of a periodicity representation hinders its ability to detect these tones.

**Supplementary Figure 14.**
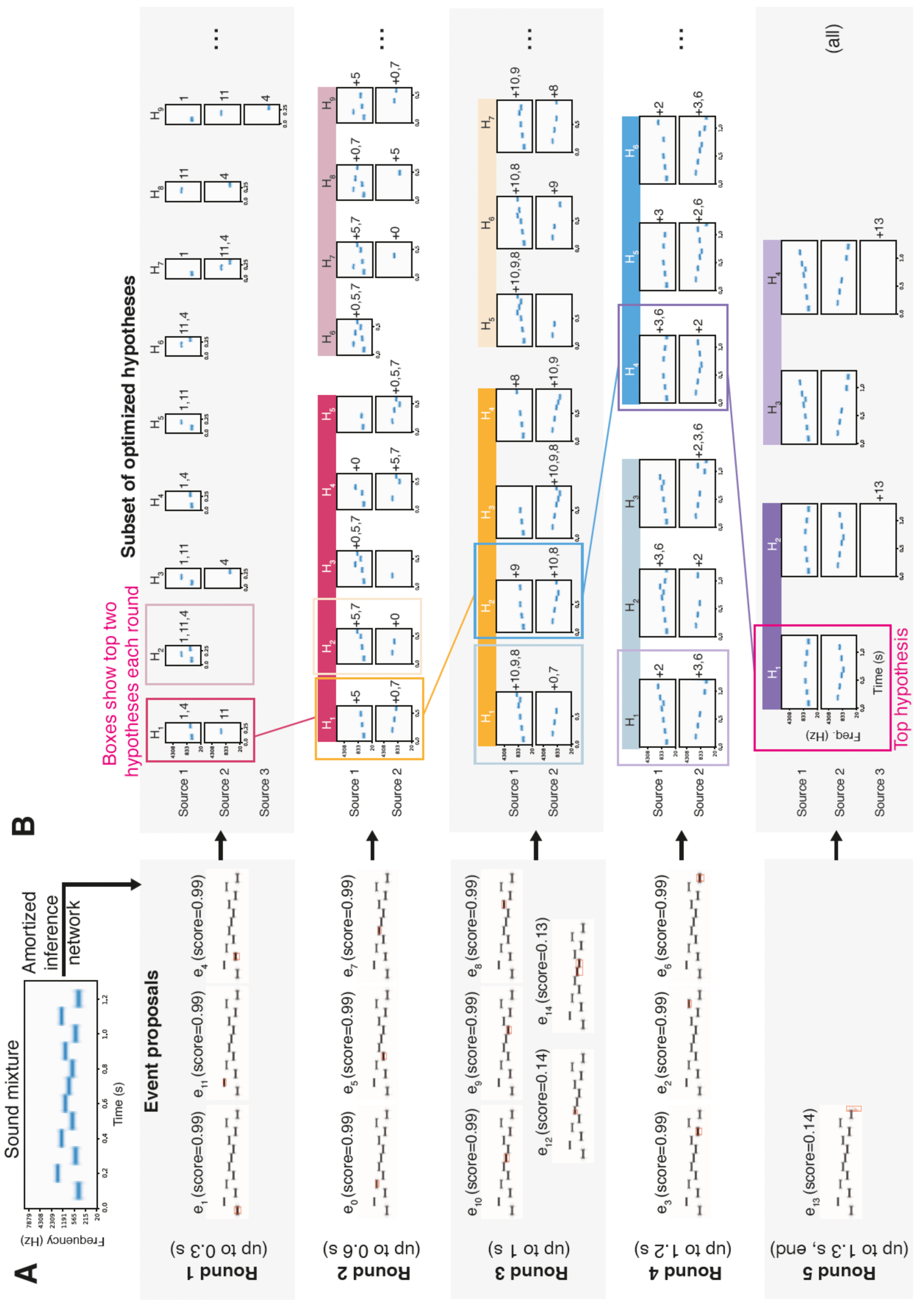
Example of sequential inference algorithm with a tone sequence. A) The observed sound mixture is 1.3 seconds long. Given the sound mixture as input, the amortized inference network returns a set of event proposals. Each event proposal is depicted as a red mask overlaid on the sound mixture cochleagram, subtitled with its network confidence, and grouped based on onset interval. For example, the first three event proposals (by onset) are grouped together and used for source construction in Round 1 (which considers the observed sound up to 0.3 sec). B) A subset of optimized hypotheses for each round, with boxes that encircle the two hypotheses with the highest posterior probability each round. The high-saturation path demonstrates how the top hypothesis for the whole observation (“Top hypothesis” with pink box, Round 5) is gradually built up over multiple rounds. Each hypothesis (indicated by Hi) is composed of combining candidate events into sources. The candidate events used for each source are indicated to the right of the source cochleagram, with the numbers referring to the title (“ei”) of the event proposal in A. Both top hypotheses on a round are used for source construction in the next round, indicated by the matching color of the hypothesis box and the thick bar on the next round. Only a subset of the hypotheses are depicted for Rounds 1-4 (“…” on right). Round 5 shows all of the hypotheses optimized (“all” on right).

**Supplementary Figure 15.**
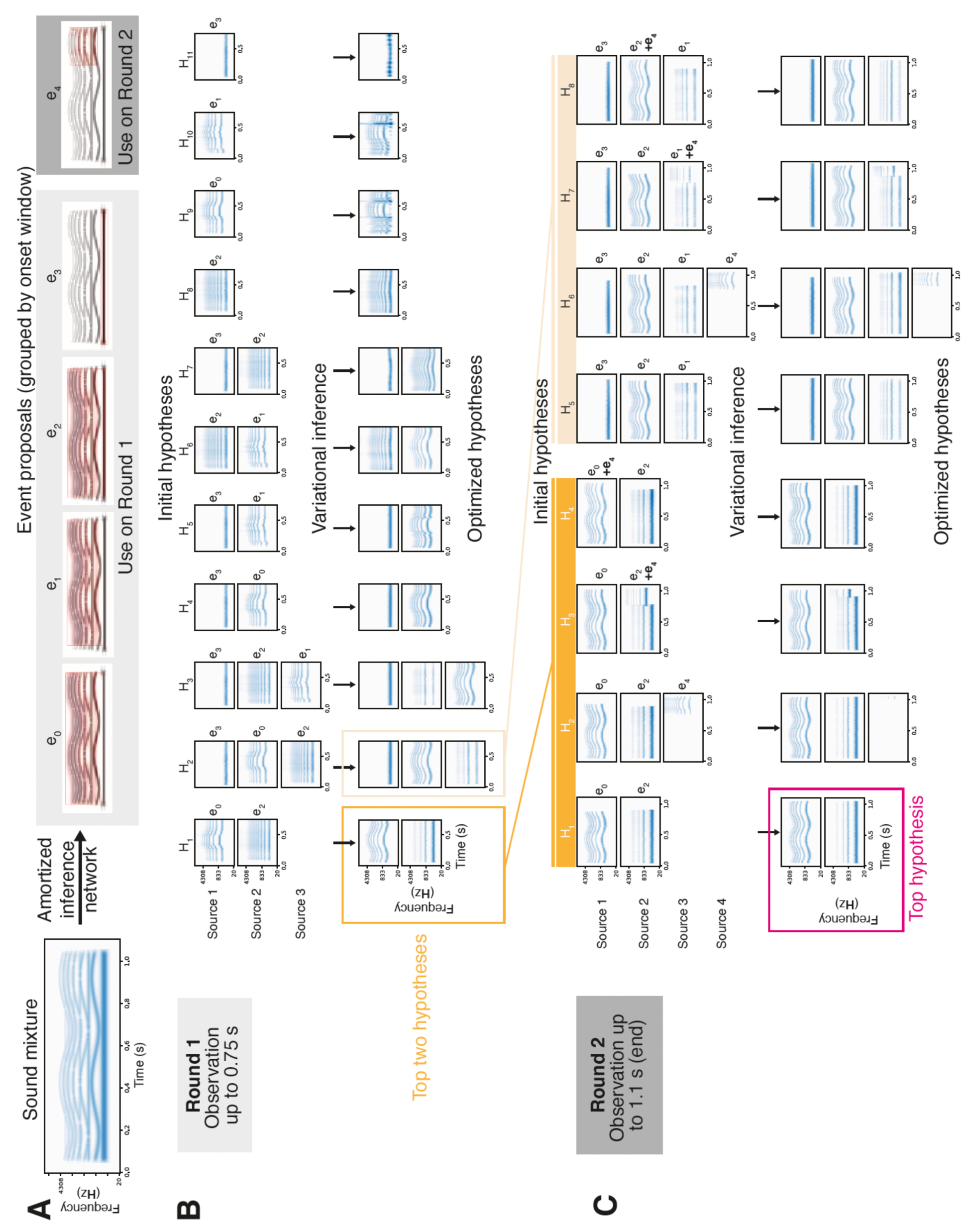
Example of sequential inference algorithm with frequency modulation. A) Event proposal network. The observed sound mixture is 1.1 seconds long. Given the sound mixture as input, the amortized inference network returns a set of five event proposals. Each event proposal ei is depicted as a red mask overlaid on the sound mixture cochleagram, and is associated with a set of event latent variables (not shown). Event proposals with nearby onsets are considered in the same round of sequential inference. B) Round 1 of sequential inference. On Round 1 of inference, the first 0.75 seconds of the observation are considered and the first four event proposals are utilized. Initial hypotheses are constructed by combining different sets of event proposals subject to the heuristic constraints described in Methods (Source Construction) resulting in eleven initial hypotheses {Hi}; note that all of the actual hypotheses used during inference are depicted here, in contrast to Supplementary Figure 14. The initial hypotheses are optimized with variational inference, resulting in optimized event-level variables (as reflected by the change in the rendered cochleagrams) and source-level variables (not shown). The two hypotheses with the highest posterior probability are used in source construction for Round 2. C) The second round of sequential inference. The full observation is considered and therefore this is the last round. Initial hypotheses are constructed by using H1 and H2 of round one (bright and dim yellow, respectively), potentially combining them with event proposal e4. The initial hypotheses are optimized with variational inference. After variational inference, the hypothesis with the highest posterior probability is selected.

## Footnotes

1 https://mcdermottlab.mit.edu/mcusi/bass/

2 https://mcdermottlab.mit.edu/mcusi/bass/

